# Principles of gene regulation quantitatively connect DNA to RNA and proteins in bacteria

**DOI:** 10.1101/2021.05.24.445329

**Authors:** Rohan Balakrishnan, Matteo Mori, Igor Segota, Zhongge Zhang, Ruedi Aebersold, Christina Ludwig, Terence Hwa

## Abstract

Bacteria allocate their proteome to cellular functions differently in different growth conditions. It is largely unknown how such allocation arises from known mechanisms of gene regulation while constrained by limited translation capacity and fixed protein density. Here, we performed absolute transcriptomic and proteomic analysis for *E. coli* across many conditions, obtaining a plethora of results on promoters and mRNAs characteristics that clash with conventional expectations: the majority of mRNAs exhibit similar translational efficiencies, while the promoter strengths are vastly different across genes. These characteristics prescribe two principles of gene regulation guiding bacteria to attain the desired protein allocation under global constraints: Total transcriptional output is tightly coordinated with ribosomal activity, and the concentrations of individual proteins are largely set by transcription. These two principles lead to a quantitative formulation of Central Dogma which unravels the complex relationship between gene regulatory activities and mRNA/protein concentrations across conditions. The knowledge obtained will be invaluable for accurately inferring gene regulatory interactions from ‘omics data, as well as for guiding the design of genetic circuits for synthetic biology applications in *E. coli* and other organisms.

## INTRODUCTION

Gene expression involves the transcription of genes into mRNAs, followed by the translation of mRNAs into proteins. Concentrations of proteins in cells are in turn determined by the balance between protein synthesis and dilution (Fig. 1A, Fig. S1) for exponentially growing bacteria where protein degradation is negligible [1, 2]. According to the canonical picture of bacterial gene expression [3–5], doubling the transcription initiation rate of a specific gene by an activator would result in doubling the concentration of the corresponding mRNA and protein in the absence of post-transcriptional processes. However, this simple picture breaks down at the global level: if we attempt to double the transcription initiation rate of every gene by a global activator, clearly protein concentrations cannot double, since the synthesis of proteins is constrained by the translational capacity of the ribosomes [6, 7]. Moreover, it is known for the best quantitatively characterized model bacterium *E. coli* that the total number of proteins per cell volume is approximately constant, ∼3 × 10^6^/μm^3^, across many different growth conditions (Fig. S2; see also [8]). Thus, the canonical single-gene picture of bacterial gene expression is at odds with global constraints, making it difficult to predict the effects of transcriptional and translational regulation on mRNA and protein levels, or conversely, to infer underlying regulatory processes from observed changes in mRNA and protein levels. These difficulties are expected to be generic whenever global changes in gene expression are encountered, e.g., upon changes in nutrient conditions or exposure to antibiotics.

**Figure 1.**
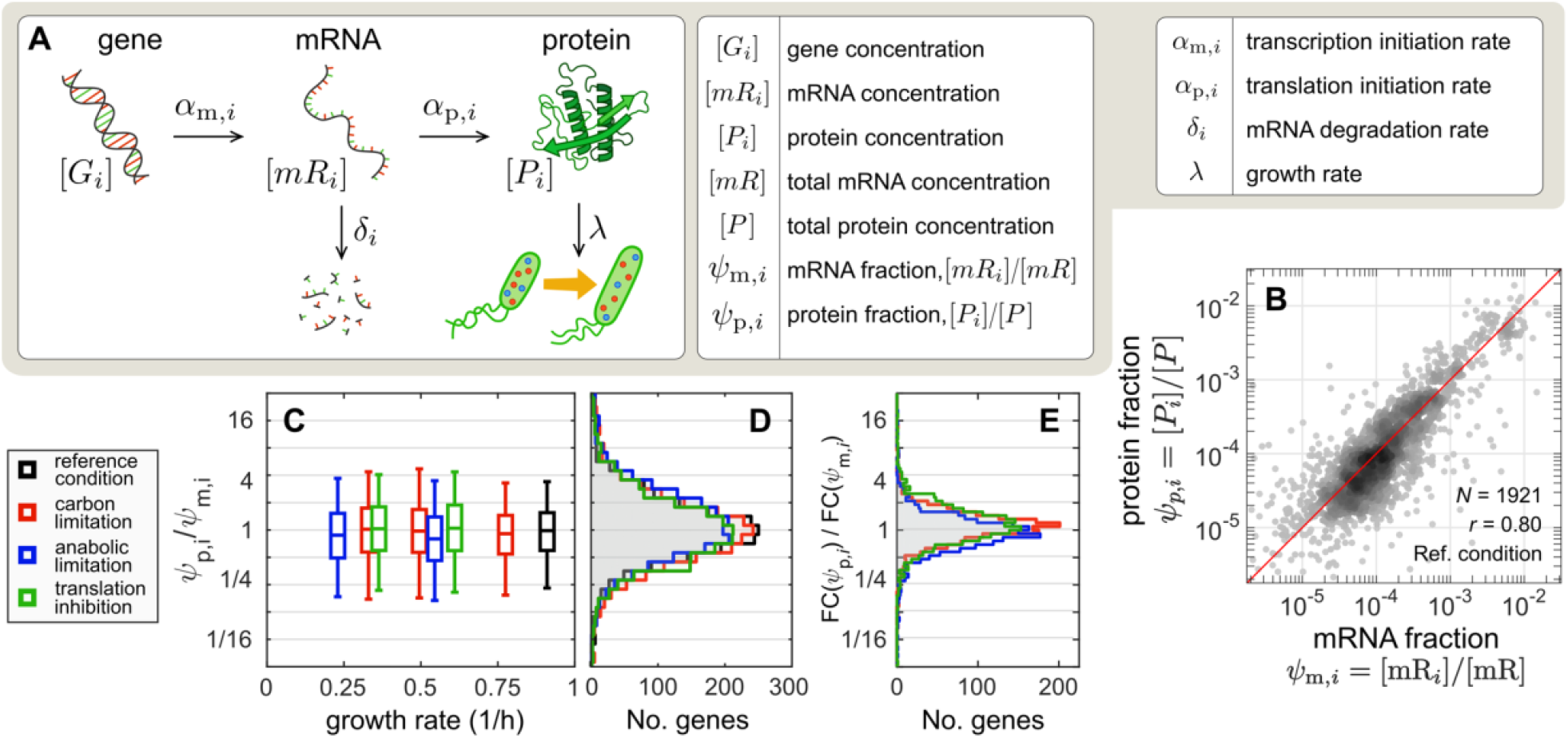
Genome-wide mRNA and protein comparison. **(A)** Schematic illustration of the basic processes determining mRNA and protein concentrations in exponentially growing bacteria; the symbols used throughout the study are described alongside the respective cellular processes (see also Figure S1). **(B)** For *E. coli* K-12 strain NCM3722 growing exponentially in glucose minimal medium (reference condition, growth rate 0.91/h), the fractional number abundances of proteins (*ψ*_p,*i*_, obtained from DIA/SWATH mass spectrometry [11] and of mRNAs (*ψ*_m,*i*_, obtained from RNA-sequencing; see Methods) for each gene *i* are shown as scatter plot (number of genes and Pearson correlation coefficient in figure). The red line represents the diagonal, *ψ*_p,*i*_ = *ψ*_m,*i*_. **(C)** The ratios of protein and mRNA fractions, *ψ*_p,*i*_ *ψ*_m,*i*_, are distributed around 1 for exponentially growing cultures under all growth conditions studied (Fig. S3A-P). These include the reference condition (black), as well as conditions of reduced growth, achieved by limiting carbon catabolism (red), anabolism (blue), or inhibiting translation (green); see SI Methods. Boxes and the whiskers represent 50% and 90% of the genes, respectively; x-axis values give the corresponding growth rates. See Table S1 for list of strains and conditions in this study and Table S2 for transcriptomics and proteomics data. **(D)** Distributions of the ratios *ψ*_p,*i*_ *ψ*_m,*i*_ obtained in reference condition and the slowest-growing of each of the three types of limitations; same color code as (C). The same plots also give the distributions of the relative translational initiation rate, 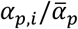; see text. **(E)** The fold-changes in protein and mRNA fractions for each gene *i* between the reference condition and the slowest growth condition, FC(*ψ*_p,*i*_) and FC(*ψ*_m,*i*_), were computed as described in Fig. S4 for each one of the three growth limitations; the distribution of their ratio FC(*ψ*_p,*i*_) FC(*ψ*_m,*i*_) is shown using the same color code as (C). The histograms are narrowly distributed around 1, with more than half of the genes within 35% from the median. See Table S3 for the fold changes in translation efficiency for each gene.

As the expression of each gene is ultimately determined by the rates at which the respective mRNA and proteins are synthesized and diluted (Fig. S1), we designed a battery of experiments to determine these rates through measuring the absolute mRNA and protein concentrations and their fluxes, for *E. coli* growing exponentially under a variety of conditions [9, 10]. Our findings establish characteristics of promoters and mRNAs that defy conventional expectations, and reveal design principles underlying *E. coli*’s gene regulation program which enable the cell to allocate its proteome in accordance to functional needs while complying with global constraints. In addition to identifying a key regulator implementing the global regulatory program, we established a simple, quantitative relation which connects gene regulatory activities to mRNA and protein levels, and reconciles the canonical gene-specific view with global constraints.

## RESULTS

### Translation initiation rates are similar across mRNAs and growth conditions

Using a recently developed proteomics workflow that accurately quantifies the abundance of individual *E. coli* proteins [11] by combining the versatility of data-independent acquisition (DIA) mass spectrometry [12, 13] and the accuracy of ribosome profiling [14], we determined the protein number fractions *ψ*_p,*i*_ ≡ [*P*_*i*_]/[*P*] for >1900 proteins (labeled by *i*), with [*P*] = ∑_*i*_[*P*_*i*_] being the total protein concentration, defined here as number of proteins per cell volume. Similarly, transcriptomics (RNA-seq) was used to determine the mRNA number fractions *ψ*_m,*i*_ ≡ [*mR*_*i*_]/[*mR*] for the corresponding mRNAs, with [*mR*] ≡ ∑_*i*_ [*mR*_*i*_] being the total mRNA concentration; see SI Methods. The result for *E. coli* K-12 cells growing exponentially in glucose minimal medium is shown as a scatter plot of *ψ*_p,*i*_ vs *ψ*_m,*i*_ in Fig. 1B. We observed a strong correlation (*r* = 0.80) along the diagonal (red line) across a vast range of abundances (10^−2^ to 10^−6^). The histogram of *ψ*_p,*i*_ /*ψ*_m,*i*_ is peaked around 1, with 50% of the genes within 1.7-fold (Fig. S3B). We next repeated the proteomics and transcriptomics measurements for cells growing exponentially in three types of growth limiting conditions in minimal medium (carbon limitation, anabolic limitation, and translational inhibition [9, 10], with growth rates ranging between 0.3/h and 0.9/h. A similar number of gene products are detected in these conditions and the resulting scatter plots and histograms (Fig. S3C-P) look very similar to those in glucose minimal medium (Fig. S3AB). These results, summarized in Fig. 1C-D, strongly indicate that the fractional abundances of mRNA and proteins are approximately the same, i.e.,

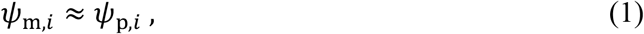

for the vast majority of expressed genes in all growth conditions tested.

The strong correlation between mRNA and protein fractions observed here, seemingly at odds with numerous earlier reports that emphasize discordance between the two [15–21], is in fact also embedded (although not articulated) in several recent quantitative measurements of *E. coli* protein expression using ribosome profiling (Fig. S3Q-T). To probe how changes in protein and mRNA fractions are related to each other across growth conditions, we generated additional proteomics and transcriptomics datasets for more conditions under each type of growth limitations (Fig. S4A), so that a smooth growth-rate dependence can be obtained individually for the mRNA and protein fractions (Fig. S4B). We then extrapolated these data to compute the fold-change (FC) in the protein and mRNA fractions, FC(*ψ*_p,*i*_) and FC(*ψ*_m,*i*_) respectively, for each gene *i* (Fig. S4B). The fold-change was calculated between the “reference condition” (WT cells grown in glucose minimal medium) and one with ∼3x slower growth, for each of the three types of growth limitation imposed. Their ratio, FC(*ψ*_p,*i*_)/FC(*ψ*_m,*i*_), is even more tightly distributed than *ψ*_p,*i*_ /*ψ*_m,*i*_ for each type of growth limitation (compare Fig. 1E with Fig. 1D), indicating that the mRNA and protein fractions tightly co-vary for the majority of genes. The few exceptions which do not co-vary usually occur in only one of the growth limitations, and mostly correspond to known targets of post-transcriptional regulation (Fig. S4C-E, Table S3).

### Total mRNA abundance matches the translational capacity

From the steady state relation between concentrations of individual mRNAs and proteins (Fig. S1), i.e.

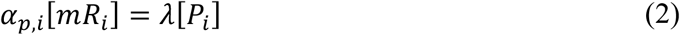

where *α*_*p,i*_ is the translation initiation rate of each mRNA *mR*_*i*_ and *λ* denotes the growth rate, we can sum over contributions from all genes to obtain a relation between the flux of total protein synthesis and dilution,

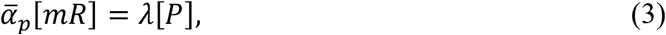

with 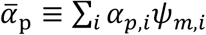 being the average translational initiation rate (over all mRNAs). Since the ratio of Eq. (2) and Eq. (3) gives

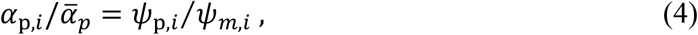

we see that Fig. 1D also provides the distribution of the relative translation initiation rates 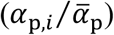. The observed similarity between mRNA and protein fractions, Eq. (1), implies that the translation initiation rates *α*_*p,i*_ are similar for the majority of mRNAs for each growth condition. Thus, the average translational initiation rate 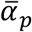 can be taken as representative of the majority of mRNAs.

Eq. (3) describes a constraint between the total mRNA concentration [*mR*] and the total protein synthesis flux *λ*[*P*] across all growth conditions, since [*P*] ≈ 3 × 10^6^/μm^3^ (Fig. S2F) is approximately constant. To understand how this constraint is accommodated by gene regulatory processes, we quantified the total mRNA amount, an important quantity which is often left out of transcriptomics analysis (Fig. S5, SI Methods). This was done by hybridizing ^3^H-uracil labeled RNA to genomic DNA and quantifying the radioactivity signal relative to that in glucose minimal media (Fig. S5A-C), and further quantifying the absolute abundance by calibrating RNA-seq data on multiple genes using quantitative Northern Blotting (Fig. S5D-H, SI Methods). The result, shown in Fig. S5H for carbon-limited growth, was then converted to cellular concentration, i.e., [*mR*] (Fig. S2C and SI Note S1), and shown as the red symbols in Fig. 2A (left vertical axis). This data allowed us to use Eq. (3) to obtain the average translation initiation rate 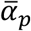. The approximately linear growth-rate dependence of [*mR*] makes 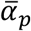 only weakly growth-rate dependent; see red symbols in Fig. 2B (left axis). The value of 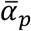 in turn allows us to obtain the distribution of *α*_*p,i*_, the translational initiation rate of individual mRNAs, using Eq. (4) and the distributions of *ψ*_*p,i*_ /*ψ*_*m,i*_ (Fig. 1D). The results, shown either as a fraction of genes or mRNAs (Figs. 2C, 2D respectively) for the reference and a slow, carbon-limiting growth condition (black and red lines, respectively), exhibit weak dependence of *α*_*p,i*_ on both the mRNA species and growth condition.

**Figure 2.**
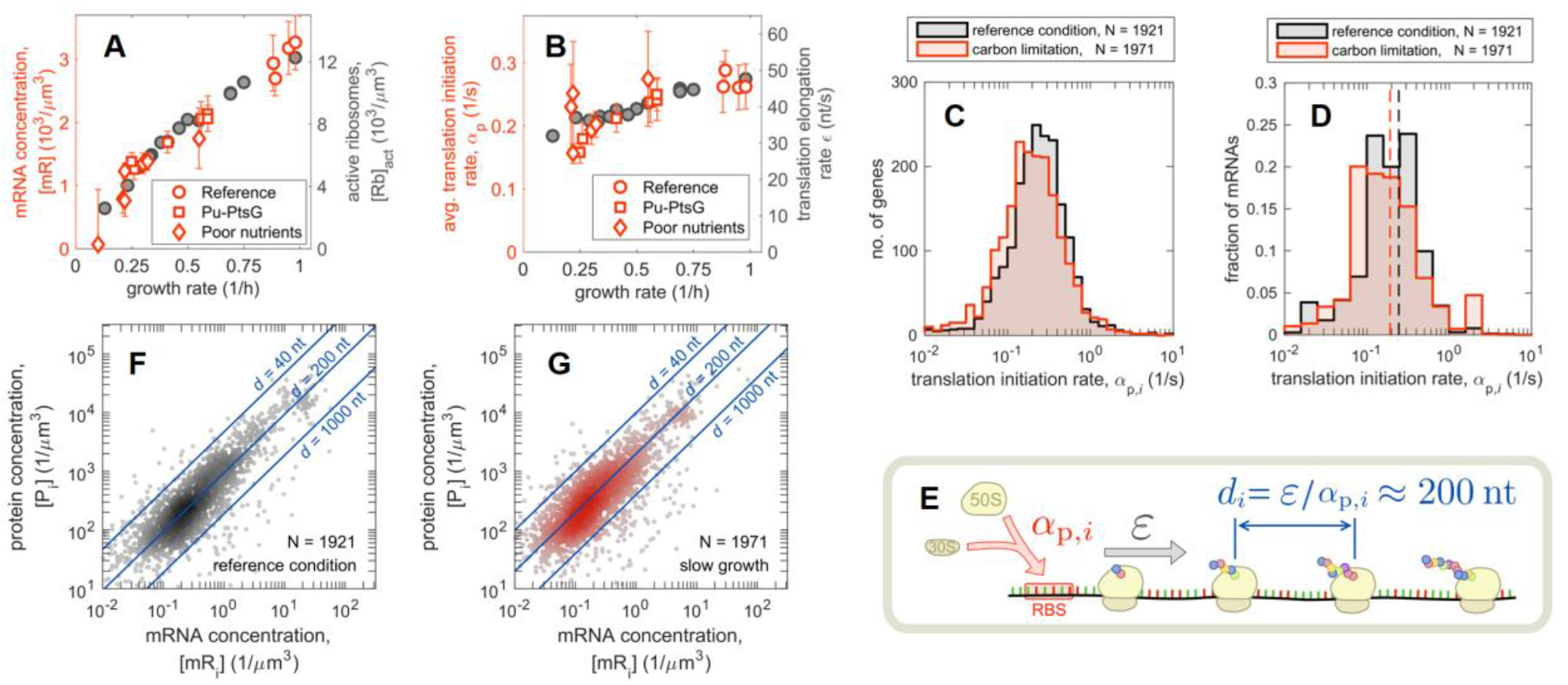
Coordination of mRNA and ribosome abundances. **(A)** Left axis (red symbols): total concentration of mRNA is plotted against the growth rate. Total mRNA abundance was obtained as described in Fig. S5 and SI Methods. The measurements were performed for a range of growth conditions, including reference, glucose uptake titration (Pu-*ptsG*, see Table S1) and a host of poor carbon sources. Right axis (grey symbols): concentration of active ribosomes in nutrient-limited conditions, converted from the data in Ref. [7] (reported in per culture volume) using the total cellular volume shown in Fig. S2C. **(B)** Left axis (red symbols): average translation initiation rate,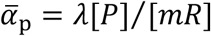, as a function of growth rate. Right axis (grey symbols): translational elongation rate from Ref. [7]. **(C)** Distribution of translation initiation rates, 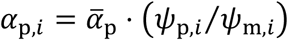, for reference condition and slow glucose-limited growth. **(D)** Same as panel (C), but weighting the histogram by mRNA abundance. The dashed lines indicate the values of the average initiation rates 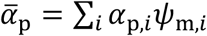 in the two conditions, 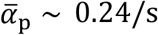 in reference condition and 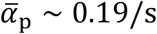 at slow growth. **(E)** The spacing between consecutive translating ribosomes on an mRNA is given by the ratio between the ribosome elongation rate (similar across mRNAs, [7]) and the translation initiation rate *α*_*p,i*_, which is also narrowly distributed (see panel C). Our data give an average ribosome spacing of *d* ≈ 200 nt; see Figure S6B. **(F)** Absolute mRNA and protein concentration for each gene in reference condition, computed by combining the fractional abundances *ψ*_m,*i*_ and *ψ*_p,*i*_ with total mRNA abundances (panel A), total protein abundances and cell volume (see Figure S2 and Note S1). Blue lines indicate the corresponding values of inter-ribosome spacing *d*, calculated from the known elongation rates (∼15.3 aa/s). **(G)** Same as panel (F), but for slow growth in the most C-limiting condition (growth rate ∼ 0.35/h, elongation rate ∼12 aa/s).

### Constancy of ribosome spacing across mRNA and nutrient conditions

To understand how the relation between the total mRNA concentration and the total protein synthesis flux (Eq. (3)) arises in molecular terms, we note that the total flux of peptide synthesis, is given by 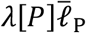, where 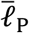 is the average length of a protein, approximately 250 aa across conditions (Fig. S2E). This flux corresponds to the product of the concentration of actively translating ribosomes ([*Rb*]_*act*_) and the speed of translational elongation (*ε*) as depicted in Fig. S6A, i.e.,

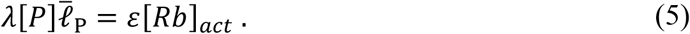

Both [*Rb*]_*act*_ and *ε* have been characterized previously for a broad range of nutrient conditions [7]. The active ribosome concentration is plotted as grey symbols in Fig. 2A (right vertical axis). The data exhibits a striking congruence with the total mRNA concentration (compare grey and red symbols in Fig. 2A), revealing a global coordination of mRNA abundance and the cellular translational capacity, with an average spacing between translating ribosomes close to 200 nt (Fig. S6B), or about 4 ribosomes for a typical mRNA 750 nt long.

The dependence of the average number of translating ribosomes per mRNA,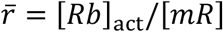,on molecular parameters can be obtained by combining Eq. (3) and (5), leading to 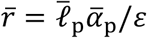. Hence, the proportionality between total mRNA and active ribosome concentrations (Fig. 2A and Fig. S6B) implies that the average translational initiation rate 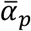 and elongation speed *ε* are proportional. This is consistent with our data (Fig. 2B), where the values of *ε* (taken from [7] for cells grown in various nutrient conditions) are shown as the grey symbols (right vertical axis).

Since the translational initiation rates of individual mRNAs (*α*_*p,i*_) are distributed tightly around the average (Figs. 2C, 2D), this implies that *α*_*p,i*_ ∝ *ε*, which leads to the observed constant spacing of 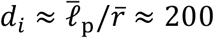 nt between the translating ribosome along the length of mRNAs (Fig. 2E).

We examined the predicted relation between ribosome and mRNA by working out the concentrations of proteins and mRNAs (Fig. S6CD) and making scatter plots of these quantities for the reference and slowest carbon-limiting conditions (Figs. 2F, 2G). The blue lines show the expected number of proteins per mRNA for different ribosome spacing on the mRNA (SI Note S2). Thus, ribosome spacing is indeed clustered around 200 nt in both growth conditions for typical mRNA species. Distributions of ribosome spacing over genes and mRNAs (Figs. S6E, S6F), show that the data are bounded by ∼40 nt per ribosome in accordance to the physical packing limit [22].

### mRNA degradation is largely condition independent

We next investigated the mechanism behind the observed proportionality between the concentrations of total mRNA and the active ribosomes (Fig. 2A). The mRNA concentration is set by the balance between its synthesis and degradation (Fig. S1). We performed kinetic experiments to determine the mRNA degradation rates *δ*_*i*_ genome-wide in the reference and the slowest carbon-limiting condition by inhibiting transcription initiation using rifampicin and quantifying the relative mRNA levels at short time intervals using RNA-seq (Fig. S7A-C, SI Methods, Table S2). As an example, we show in Figure 3A-C time courses of changes in the relative mRNA levels of genes of the *nuo* operon in the two growth conditions. The time course can be described as a delayed exponential decay, with the lag time reflecting the time needed for the RNA polymerase to reach the gene from the transcription start (Fig. S7D), and the decay rate attributed to the turnover of that mRNA. This analysis yielded degradation rates for ∼2700 mRNAs (SI Methods, Table S4). At the genome-wide level, we found the mRNA degradation rates to be strongly correlated in the two growth conditions (Fig. S7E). The average degradation rate is hardly different (Fig. S7F, vertical dashed lines), even after weighting by mRNA abundances (Fig. S7G). In particular, the fold-change in *δ*_*i*_ is sharply peaked, with 90% of genes in the range 0.50 to 1.57 (Fig. 3D). The data indicates the lack of dependence of degradation rates on either the mRNA species or the growth condition for most mRNAs, as is consistent with earlier studies [23, 24].

**Figure 3.**
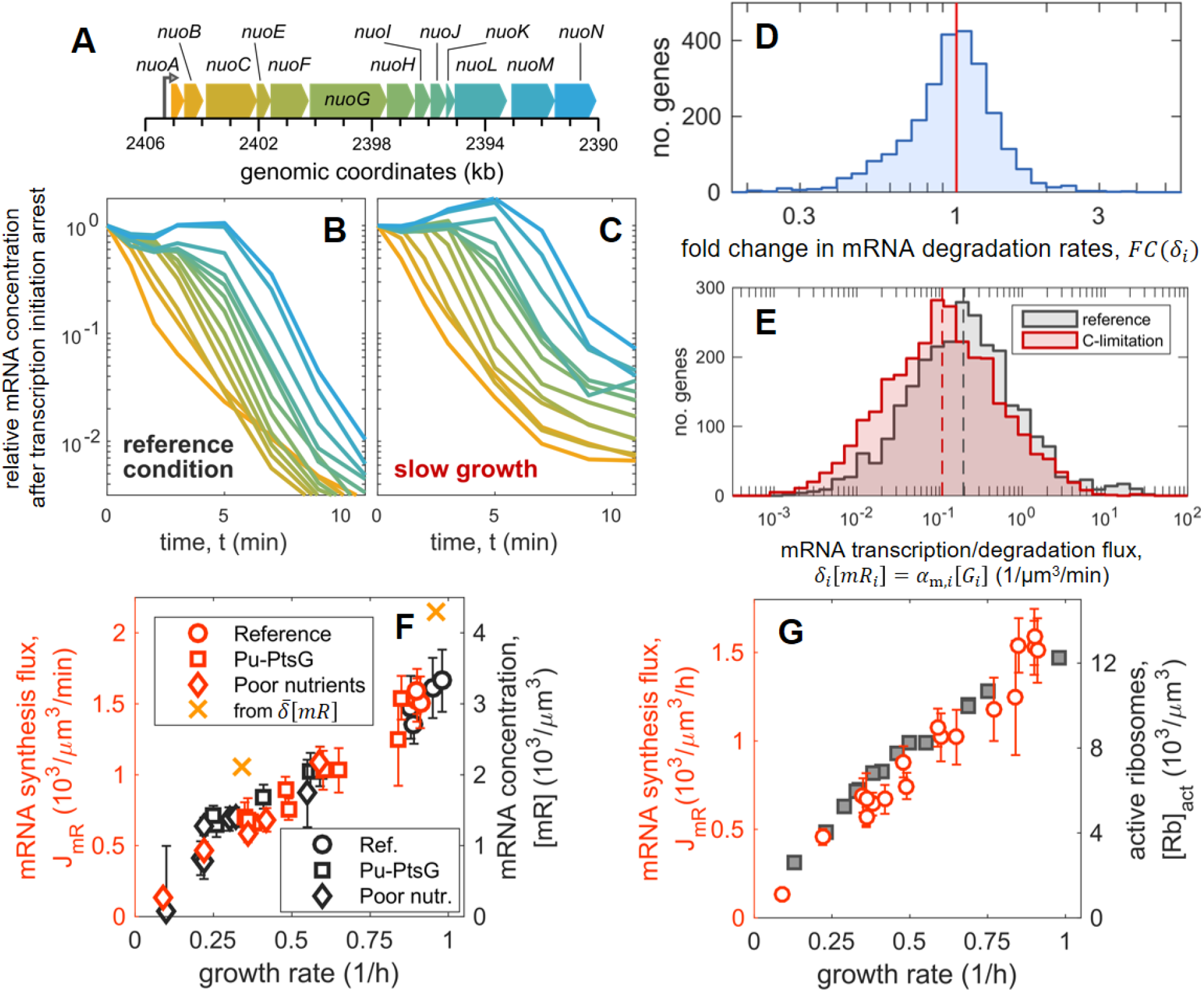
mRNA degradation and synthesis. **(A-C)** Degradation of mRNA transcribed from the long *nuo* operon (A) in reference condition (B) and carbon-limited condition (C). The abundance of mRNA was measured by RNA-seq over the course of 11 minutes following the blockage of transcription initiation by rifampicin (SI Methods, Figure S7). While the abundance of the mRNA of genes proximal to the promoter (*nuoA*, orange) drops immediately after rifampicin treatment (at time *t* = 0), a lag is observed for genes progressively more distant from the promoter (from orange to blue). The lag time corresponds to the time elapsed between the transcription of the proximal and distant genes by RNAPs which initiated transcription before the application of rifampicin (Fig. S7D). **(D)** Histogram of fold-change of the mRNA degradation rates, *FC*(*δ* _*i*_), between carbon limited medium and reference condition for *N* = 2550 genes. Half of the fold changes are within 25% from unity, and 90% of the fold changes are in the range 0.50 to 1.57, implying that the degradation rates for most mRNAs do not change significantly between the reference and carbon-limited growth conditions. **(E)** Distribution of the mRNA degradation fluxes, *δ*_*i*_ [*mR*_*i*_], computed from the mRNA concentration and degradation rates. These quantities should equate the mRNA synthesis fluxes, *α*_m,*i*_ [*G*_*i*_], in steady state conditions. Dashed lines indicate the median fluxes, 0 1 μm^3^/m in reference condition and 0 10 μm^3^/min at slow growth. **(F)** Left axis (red symbols): total mRNA synthesis flux *J*_mR_ = ∑_*i*_ *α* _m,*i*_ [*G*_*i*_] (transcripts synthesized per cell volume per unit time), for a variety of growth conditions as indicated (see Table S1 for growth conditions). This was obtained from the measured total mRNA flux per culture volume by pulse-labeling RNA with ^3^H-uracil, followed by hybridization to genomic DNA (Figure S10). The orange crosses indicate the total mRNA synthesis flux obtained from summing *δ*_*i*_ [*mR*_*i*_] using the data in (E). Right axis (black symbols): absolute mRNA abundances (same data as Fig. 2A). **(G)** Left axis (red symbols): total RNA synthesis flux vs. growth rate (same data as in panel (F)). Right axis (grey symbols): concentration of active ribosomes (same data as Fig. 2A).

The lack of large changes in mRNA degradation rates does not mean the absence of post-transcriptional regulatory processes, which often leave their signatures as changes in mRNA stability [25–27]. A closer comparison of the distribution of the degradation rates in the two conditions suggests a group of mRNAs with *δ*_*i*_ ∼1/min in reference condition and ∼0.5/min in the slow-growth condition (indicated by the triangular region in Fig. S7E or the arrow in Fig. S7F). A further look into those mRNAs with altered stability (Fig. S7HI) found a number of known effects of post-transcriptional regulation, where changes in mRNA stability contributed significantly to changes in mRNA concentrations. However, our data indicates that, at the genome level, the magnitude of such post-transcriptional regulation is limited, often not more than 2-fold. Moreover, even those mRNAs affected at this modest level constitute a minority, at least for exponentially growing cells under the conditions tested.

### Total mRNA synthesis flux is adjusted to match translational capacity

From the concentration and degradation rate of each mRNA species, [*mR*_*i*_] and *δ*_*i*_ respectively, we can obtain the mRNA degradation flux, *δ*_*i*_ [*mR*_*i*_], whose distributions are shown in Fig. 3E for the reference and slow growth conditions. A clear shift in the median of the two distributions is seen (vertical dashed lines), reflecting growth dependence of the total degradation flux,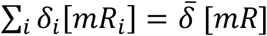, where 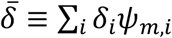 is the average degradation rate across mRNAs. By the balance of mRNA synthesis and degradation in steady state growth (Fig. S1; see also SI Note S3), the total mRNA synthesis flux *J*_*mR*_ can be expressed as

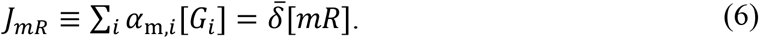

Since the average degradation rate 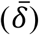 is hardly affected by growth conditions (Fig. S7G), Eq. (6) predicts that the observed growth dependence of the total mRNA concentration [*mR*] (grey symbols in Fig. 2A) arises primarily due to changes in the mRNA synthesis flux, *J* _*mR*_.

We tested this prediction by directly measuring the total mRNA synthesis flux *J* _*mR*_ across the range of carbon-limited growth conditions, by pulse-labelling cultures with ^3^H-uracil and hybridizing the labelled RNA to genomic DNA over short time intervals (Fig. S8). The data shows a strong growth-rate dependence Fig. 3F (red circles, left vertical axis), closely matching the observed growth dependence of the total mRNA concentration (reproduced as grey symbols in Fig. 3F, right vertical axis) as predicted. Note that the total mRNA fluxes inferred from the degradation rates, 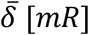 (orange crosses) are within 20% of the directly measured synthesis fluxes, showcasing the consistency of these two very different measurement approaches. Putting together, the results in Figs. 2 and 3 show that the global constraint Eq. (3) is enforced primarily by matching the total mRNA synthesis flux *J* _*mR*_ with the translational capacity (Fig. 3G).

### mRNA synthesis flux and transcriptional regulation

The synthesis flux of each mRNA species is given molecularly by the product of the transcription initiation rate per gene, *α*_*m,i*_, and the “gene concentration”, [*G* _*i*_]; see Fig. 1A and Fig. S1. The growth-rate dependence of gene concentration is in turn given by the product of number of chromosome replication origins (Ori) per cell volume, [Ori], and the “gene dose” relative to the Ori, *g*_*i*_ ≡ [*G* _*i*_]/[*Ori*]. Thus, the total mRNA synthesis flux can be expressed as

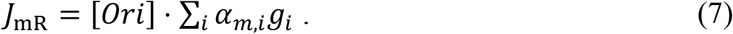

Since the relative gene dose *g*_*i*_ depends on the chromosome replication time (the C-period) and the chromosomal position of the gene [28, 29], the growth-rate dependence of *g* _*i*_ can be obtained through the growth-rate dependence of the C-period (Fig. S9AB). Further including a weak growth-rate dependence of the Ori concentration [30] (Fig. S9C-E, we obtain negative growth-rate dependences for the concentration of genes [*G* _*i*_] = [*Ori*] · *g* _*i*_ at all chromosomal positions; see Fig. 4A. It is then clear from Eq. (7) that the strong positive growth-rate dependence seen for the total mRNA synthesis flux *J*_mR_ (Fig. 3F) cannot be accounted for by the opposite growth-rate dependences of gene concentrations and must involve systematic changes in the promoter activities *α*_m,*i*_. This is seen more explicitly by computing the distributions of the promoter activity *α*_m,*i*_, obtained for each gene using the known degradation fluxes *δ*_*i*_ [*mR*_*i*_] and gene concentrations [*G* _*i*_] at steady state (Eq. S3 in Fig. S1; see also SI Note S3; data in Table S4). The results (Fig. 4B) show a broad range of promoter activity, spanning 4 orders of magnitude, with the high-end (∼0.3/s in reference condition) approaching the maximum of ∼1/s given the transcriptional elongation speed of ∼50 nt/s and a transcription elongation complex footprint of ∼40 nt [31–33]. A clear difference is seen between the reference and C-limited growth conditions (grey and red curves, respectively), with a median reduction of about 3.5-fold in *α*_*m,i*_ for ∼3-fold reduction in growth rate (Fig. 4C). Thus, the coordination of mRNA synthesis flux with the growth rate (Fig. 3F) is likely a result of global changes in transcription initiation between these conditions.

**Figure 4.**
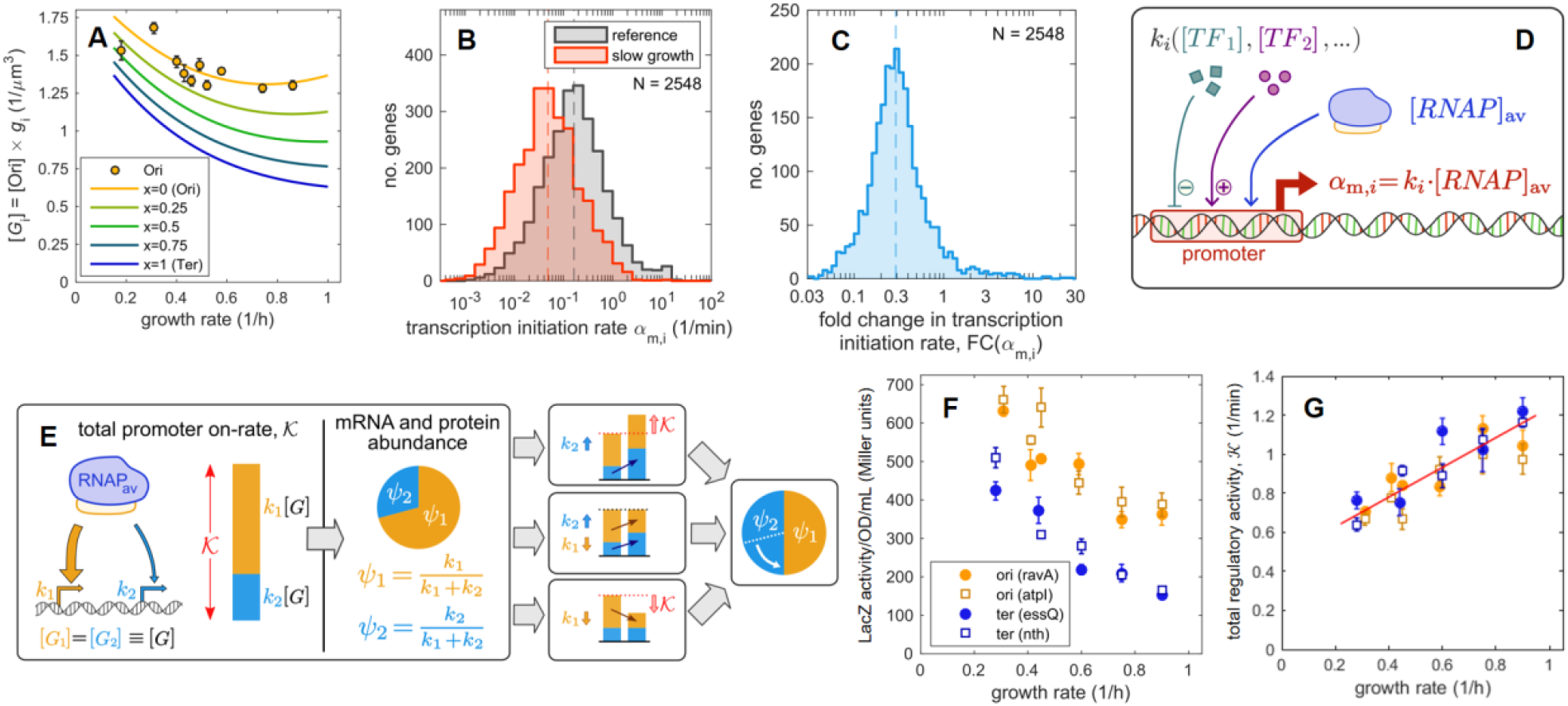
Quantitative relations between promoter on-rates and mRNA, protein abundances. **(A)** Growth rate dependence of gene concentration [*G*_*i*_] at various distances from the origin of replication Ori (solid lines). These are computed as the product of the Ori concentration [*Ori*] (orange circles, shown in Fig. S9E with raw data from [30] and the gene dose *g*_*i*_ = [*G*_*i*_]/[*Ori*] (Fig. S9B); see Figure S9 for details. **(B)** Distribution of transcription initiation rates *α*_m,*i*_ in reference condition (black) and slow growth (red), computed using the available mRNA abundances and degradation rates (see Note S3 for details). Dashed lines indicate the median initiation rates in the two conditions (2.64/min for reference condition, 0.87/min for slow growth). **(C)** Fold change of the transcription initiation rates *FC* (*α*_m,*i*_) between reference condition and slow growth. The data shows a generalized decrease of initiation rates, with a median reduction of 0.29 (dashed line) at slow growth (*λ* = 0 3/ h) compared to the reference condition (*λ* = 0 91/ h). **(D)** Illustration of a canonical model of transcriptional regulation [34, 35], with the transcription initiation rate for gene *i, α*_m,*i*_, depending on the promoter on-rate *k*_*i*_, which is modulated by transcription factors (*TF*_1_, *TF*_2_, …), as well as on the cellular concentration of available RNA polymerases ([*RNAP*] _av_), as described by Eq. (8). **(E)** Cartoon illustrating the dependence of mRNA and protein abundances on the promoter on-rates, as described by Eq. (12). Consider two genes with promoter on-rates *k*_1_ (orange) and *k*_2_ (blue) and identical gene concentration [*G*_1_] = [*G* _2_] ≡ [*G*]; the corresponding mRNA and protein fractions (*ψ*_m,1_ = *ψ*_p,1_ ≡ *ψ*_1_ and *ψ*_m,2_ = *ψ*_p,2_ ≡ *ψ*_2_, respectively) depend on both promoter on-rates via the total regulatory activity 𝒦 = *k* _1_ + *k* _2_)[*G*] (in red). Three possible scenarios are illustrated. Top: If *k* _2_ increases, while *k* _1_ remains constant, then 𝒦 increases, resulting in the reduction of protein and mRNA abundances for the orange gene despite it not being downregulated at the transcriptional level. Bottom: If only *k* _1_ decreases while *k* _2_ remains constant (bottom), then the proteins and mRNAs for the blue gene increase despite the lack of change at its promoter level. Middle: If 𝒦 is unchanged (due to compensating changes in *k* _1_ and *k*_2_ in this case), then the changes in protein and mRNA fractions would reflect changes at the regulatory level. **(F)** *E. coli* strains harboring constitutive expression of *lacZ* at various locations near *oriC* (orange) and near *terC* (blue; loci listed in the legend) were grown in carbon-limited conditions (see Table S1 for strains and conditions). LacZ protein abundance per culture volume (OD·mL), reflected by *β* -gal activity (Miller units), is shown; error bars indicate standard error on the mean of two independent replicates. **(G)** The relative change in the total regulatory activity 𝒦 across growth rates was estimated from the relative change in LacZ abundance using the data in panel (F) and Eq. (14) in the text. To do so, the LacZ abundance per culture volume was converted to protein fraction by dividing by total protein mass per culture volume (Fig. S2D). The result shows a linear dependence of the total regulatory activity on the growth rate (red line). The absolute scale 𝒦 was set for the reference condition using Eq. (10) with the values for the total mRNA synthesis flux *J*_mR_ obtained from Fig. 3F, the *oriC* concentration from Fig. 4A, and the available RNAP concentration estimated as described in SI Note S5.

To look further into the determinants of transcription initiation, we turn to a canonical model of transcription regulation (Fig. 4D) [34, 35] where the transcription initiation rate *α*_*m,i*_ for gene *i* is given by the product of the available RNAP concentration ([RNAP] _av_) and the promoter on-rate *k*_*i*_, i.e.,

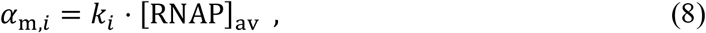

where *k* _*i*_ captures the regulatory activities of all transcription factors acting on the promoter driving gene *i* [34, 35]. Using this expression for *α*_*m,i*_, the balance of mRNA synthesis and degradation (Eq. (S3) in Figure S1) can be written as

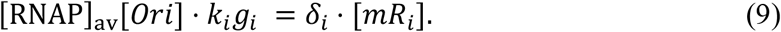

### Quantitative relations connect transcriptional regulation to gene expression

From Eq. (9), we can derive two fundamental relations connecting transcription regulation to gene expression (see also SI Note S4). Summing Eq. (9) over all genes, the balance of the total transcription flux becomes

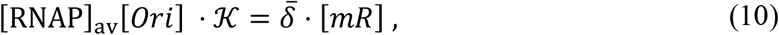

where 𝒦 ≡ ∑_*i*_ *k* _*i*_ *g* _*i*_ describes the total on-rate for promoters across the genome and is a measure of the total regulatory activity on transcription (weighted by gene dose). Given the proportionality between active ribosome and total mRNA concentrations 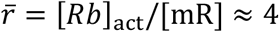 (Fig. 2A), Eq. (10) can be written as:

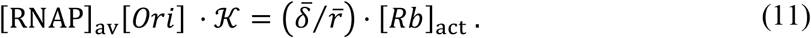

This relation represent a fundamental constraint between the overall transcription activity ([*RNAP*] _av_ 𝒦), the DNA content (via [*Ori*]) and the translational activity of the cell.

Another important relation can be obtained by taking the ratio of Eqs. (9) and (10). Noting that the mRNA degradation rates are closely distributed around the average 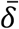 and independent of growth conditions, i.e., 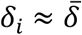 (Fig. S7FG), we obtain *k*_*i*_ *g*_*i*_ /𝒦 ≈ [*mR*_*i*_]/[*mR*] = *ψ*_*m,i*_. This relation extends further to the fractional protein abundances *ψ*_p,*i*_ = [*P*_*i*_]/[*P*] due to the established relation between protein and mRNA fractions (Eq. (1) and Fig. 1), leading to

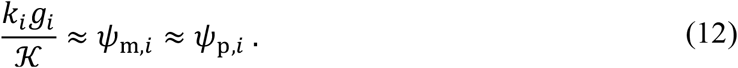

This expression relates the (gene-dose weighted) regulatory activity on specific promoters (*k* _*i*_ *g*_*i*_) to the mRNA and protein levels as determined by transcriptomics (*ψ*_m,*i*_) and proteomics (*ψ*_p,*i*_). Importantly, *ψ*_p,*i*_ = [*P*_*i*_]/[*P*] gives approximately the cellular protein concentration [*P*_*i*_] since the total protein concentration [*P*] ≈ 3 · 10^6^/μm^3^ independently of growth rate (Fig. S2F). Thus, Eq. (12) quantitatively connects regulatory activities at the promoter level (*k*_*i*_) to cellular protein concentrations [*P*_*i*_], without explicit reference to the macroscopic machineries of gene expression. Eqs. (11) and (12) are the central quantitative results of this study. We suggest Eq. (12) be viewed as a quantitative statement of the Central Dogma of bacterial gene expression, with Eq. (11) describing a global constraint on transcription and translation. In the following, we shall separately explore some consequences of these two central relations.

### Global coupling in gene expression

According to Eq. (12), the mRNA and protein levels of a given gene *i* are dependent not only on the regulatory activity on that gene, *k*_*i*_ *g*_*i*_, but also on the total regulatory activity, 𝒦 ≡ ∑_*j*_ *k*_*j*_ *g*_*j*_. The latter dependence couples gene expression globally as illustrated in Fig. 4E. This dependence is explicitly seen when comparing fold-changes in gene expression across two different conditions:

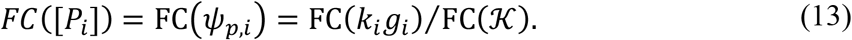

In different growth conditions where the promoter on-rates *k*_*i*_ of many genes are affected, we generally expect the total rate 𝒦 to vary, i.e. FC(𝒦) ≠ 1. Consequently, changes in the regulatory activity of a gene are generally expected to be different from the changes in the fractional abundances of the corresponding mRNA and protein. In fact, the latter might change even if the corresponding regulatory activity *k*_*i*_ *g*_*i*_ is unchanged, due to the overall change in regulatory activity 𝒦; see illustrations in Fig. 4E.

To determine how the total regulatory activity 𝒦 may change across growth conditions, we return to the spectrum of carbon-limited growth conditions. The growth-rate dependence of 𝒦 can be deduced by applying the relation (13) to “constitutively expressed” (i.e., unregulated) genes, for which *k*_*i*_ is constant. For this purpose, we inserted a constitutively expressed *lacZ* gene in the vicinity of Ori, for which *g*_*ori*_ = 1. The total rate 𝒦 can then be obtained by measuring the LacZ protein level, [*LacZ* _*ori*_] for different growth rates, as

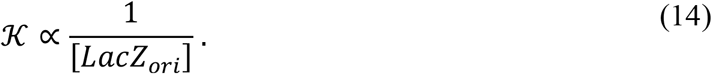

The relative LacZ protein levels obtained are shown as orange symbols in Fig. 4F across the range of carbon-limited growth. The data shows that [*LacZ* _*ori*_] increased as growth slowed down, indicating that 𝒦 has a positive growth-rate dependence (orange symbols, Fig. 4G). As a further test, we quantified the relative abundance of the same constitutively driven *lacZ* gene inserted near *terC*. In this case, the measured LacZ level exhibits a stronger negative growth-rate dependence (Fig. 4F, blue symbols). The values of 𝒦 calculated according to 𝒦 ∝ *g*_*ter*_ /[*LacZ* _*ter*_] (blue symbols, Fig. 4G) is seen to match well with the estimates from the reporter at Ori. While this set of experiments establishes the relative changes in 𝒦 across conditions, the absolute scale of 𝒦 can be determined from Eq. (10) using the measured [*Ori*] (Fig. 4A) and the measured *J*_mR_ (Fig. 3F) for 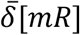 (Eq. 5). As discussed in SI Note S5, the abundance of available RNAP, [*RNAP*] _av_can be estimated in reference condition to be approximately 1000/μm^3^, leading to 𝒦 ∼ 1.2 μm^3^/m in reference condition.

### Identifying promoter on-rates

Knowledge of the magnitude of 𝒦, together with the mRNA abundances, allows us to compute the promoter on-rate *k*_*i*_ for most genes across growth conditions (see SI Note S4). The results for ∼2500 genes, shown in Fig. 5A, display a broad range across more than 3 orders of magnitude. The advantage of looking at promoter on-rates *k*_*i*_ rather than the protein concentrations (or protein fractions) is that the former provides a more direct, molecular view of transcriptional control. This is illustrated in Fig. 5B, where, as expected, we found the variation in *k*_*i*_ to be smaller than that in protein concentrations [*P*_*i*_] for genes belonging to the same operon. The difference between the two is an indicator of possible post-transcriptional control: This is, for example, clearly seen for the *sdh* operon (Fig. S10A), which contains a target of regulation by the small RNA RyhB in the translational initiation region of *sdhD*, the second gene of the operon [36]. Similarly, intra-operon variation in protein fractions, but not promoter on-rates, is seen for the *rbs* operon (Fig. S10B), which contains a regulatory target of the small RNA DsrA in the coding region of the gene, *rbsD* [37]. Unexpectedly, we also encountered a case where the intra-operon variation in promoter on-rates well exceeded that of protein fractions. This is the case of the *waa* operon (Fig. S10C), formerly known as *rfa*, encoding core-oligosaccharide assembly enzymes [38]. A steady reduction in the promoter on-rates *k*_*i*_ is seen along the length of the operon. The magnitude of this reduction, approximately 2-fold every 2500 nt, is reminiscent of the attenuation reported for premature transcriptional termination [33]. In this context, it is interesting to note that the *waa* operon requires the transcriptional anti-termination RfaH to reduce transcriptional pausing [39]. Given that the protein levels are similar across the whole operon, premature transcription termination is likely compensated post-transcriptionally, reminiscent of mechanisms discussed in Ref. [40].

**Figure 5.**
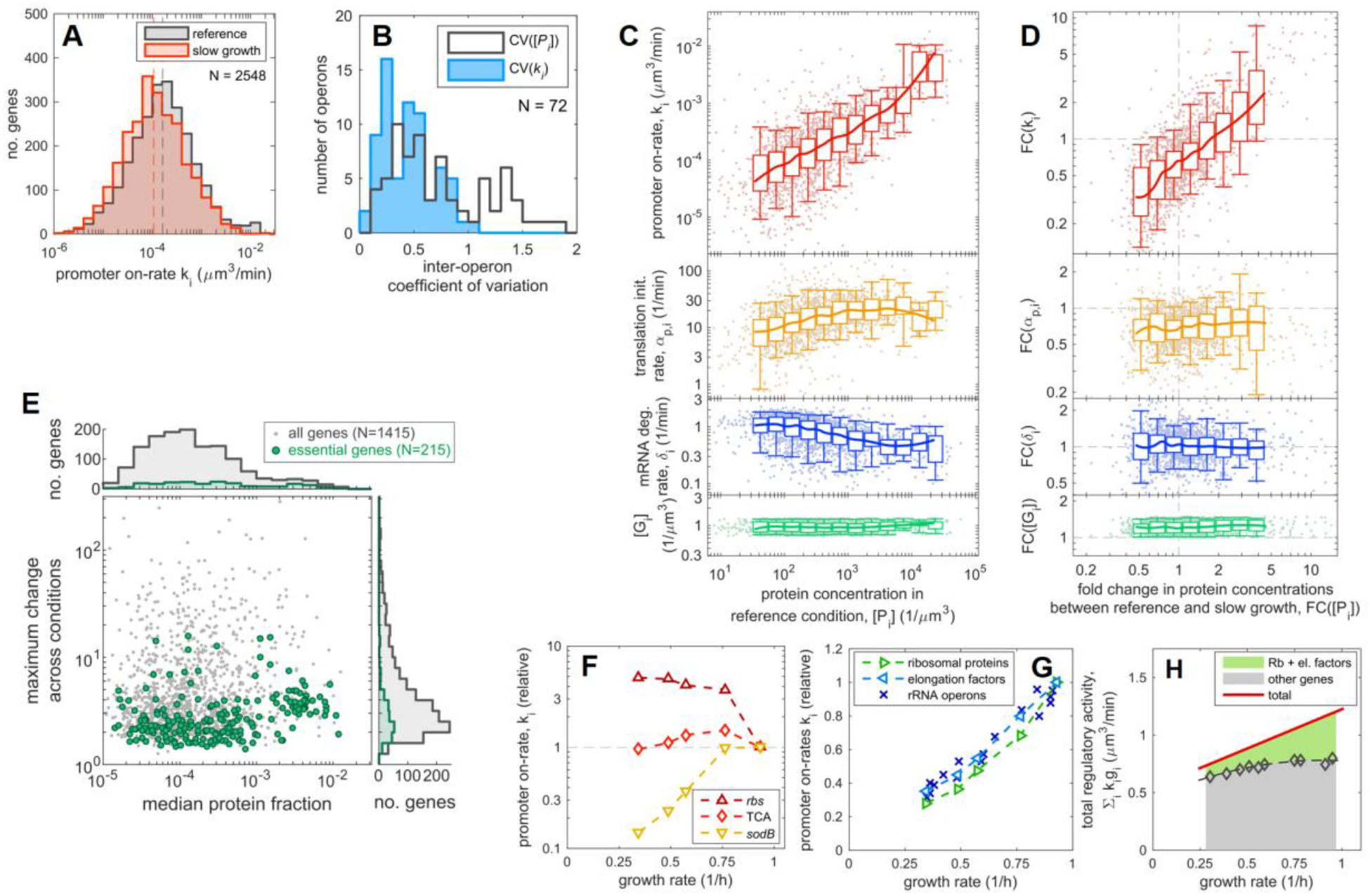
Gene expression is primarily determined by the promoter on-rates. **(A)** Distribution of promoter on-rates *k*_*i*_ in the reference and slow growth condition, obtained from the distribution of the translation initiation rate and the concentrations of available RNAP, *k*_*i*_ = *α*_*m,i*_ /[*RNAP*] _*av*_ (see Eq. (8)), as described in SI Note S4. The median promoter-on rate (vertical dashed lines) shifts from 1.63 · 10^−4^ μm^3^/min in reference condition (*λ* ∼ 0.3/ h) to 1.07 · 10^−4^ μm^3^/min in slow growth (*λ* ∼ 0 3/ h). This change is much less than the ∼3-fold change in both the growth rate and the median transcription initiation rates (Fig. 4BC). **(B)** For a set of 72 operons containing at least 3 genes according to the annotation in Ecocyc [73], we computed the coefficient of variation in protein concentrations [*P*_*i*_] and in the promoter on-rates *k*_*i*_ for genes within the operon in reference condition. In absence of post-transcriptional regulation, the inferred promoter on-rates for genes within the same operon are expected to be the same. Indeed, the promoter on-rates are more narrowly distributed (lower coefficient of variation) compared to protein concentrations. **(C)** Promoter on-rates *k*_*i*_, translation initiation rates *α*_p,*i*_, mRNA degradation rates *δ*_*i*_ and gene concentrations [*G*_*i*_] are the four molecular parameters determining cellular concentration of a protein in a given growth condition (Fig. 1A, with the transcription initiation rate *α*_m,*i*_ given by *k*_*i*_ via Eq. (8)). These four molecular parameters are plotted against the protein concentrations [*P*_*i*_] in reference condition, binned according to the observed protein concentrations. Box and whiskers indicate 50% and 90% central intervals for the binned data; the solid lines represent moving averages. **(D)** Same as panel (C), but for the fold changes of each quantity across growth conditions (slow growth compared to reference). All molecular parameters and concentrations shown in panels A-D are listed in Table S4. **(E)** 1415 genes across 19 vastly different growth conditions (Table S5). Here, we plotted for each protein the maximum fold-change of the protein fraction among any pair of conditions against the median abundance across all the conditions (gray dots). The set of genes that are annotated as essential [73] is highlighted in green. Histograms show distributions along both the axes. **(F)** Growth rate dependence of promoter on-rates summed over a few sets of genes encoding for the ribose uptake system (“rbs”, *rbsABCDK*), TCA enzymes (“TCA”, including *sucABCD, sdhACBD* and *acnAB*) and *sodB*, that encodes an iron-dependent superoxide dismutase. For all these genes protein levels increase in carbon-limited conditions (Figure S13). **(G)** Growth rate dependence of promoter on-rates summed over different groups of genes: ribosomal proteins, elongation factors (encoded by *fusA, tufAB* and *tsf*), and the rRNA operons. The activity of the rRNA operons is estimated from the synthesis flux of stable RNA (Supp. Note S5); see also Figure S14. **(H)** The sum of promoter on-rates weighted by gene dose, 𝒦 = ∑ _*i*_ *k*_*i*_ *g*_*i*_ (red line; same as in Fig. 4G) is partitioned between the contribution from ribosomal proteins and elongation factors (green) and the rest of genes (grey area). Symbols indicate the partitioning obtained from the computed *k*_*i*_ across growth rates. The growth rate dependence of 𝒦 largely stems from that of the promoter on-rates of the translational genes.

The complete set of gene expression rates generated in this work, including the promoter on-rates, mRNA degradation rates and translation initiation rates (Table S4) allowed us to investigate at the genome scale which factors control the observed protein levels in *E. coli*, as well as their fold change across conditions. By plotting the promoter on-rates *k*_*i*_ against the protein concentrations in reference condition (Fig. 5C, top), we see that the vast span of protein levels observed can be largely attributed to differences in the promoter on-rates, i.e., transcriptional regulatory activities, as opposed to other factors such as mRNA degradation rate, translation initiation rate, or the gene dose (other panels in Fig. 5C), in agreement with the simple scenario expressed by Eq. (12). The large disparity in promoter on-rates is an important characteristics of *E. coli*’s gene regulatory processes which we will return to shortly below.

At a finer level, we note that proteins present at lower concentrations tend to have lower translation initiation rates *α*_p,*i*_ and larger mRNA degradation rates *δ*_*i*_ compared to those at high concentrations (middle panels in Fig. 5C). The ratio of these two rates yields the translational “burstiness” parameter *b*_*i*_ = *α*_*p,i*_ /*δ*_*i*_, which sets the average number of proteins produced during the lifetime of a single mRNA, and is a key contributor to the stochasticity of gene expression given the low copy number of most mRNAs in bacteria (average < 1/cell; Fig. S11A); see also [3]. While the majority of mRNAs have translational burstiness ≫ 1 (within two-fold of 20, Fig. S11B), the burstiness decreases with the concentration of the expressed proteins (Fig. S11CD). Hence, low protein concentrations are attained by both reduced transcription and reduced translational burstiness (Fig. S11E).

### Unraveling the innate and regulatory effects on gene expression

When looking at the fold change in protein abundance across conditions, we see an almost perfect correlation between the fold changes in protein abundance and that in promoter on-rates (Fig. 5D, top), while the effect of post-transcriptional regulation, or gene copy number is negligible (Fig. 5D, other panels; see also Fig. S11F). This confirms that protein levels are typically adjusted across conditions by modulating the promoter on-rate as described by Eq. (13), while post-transcriptional effects do not play a major role in setting the protein concentration across conditions, except for a minority of genes as described earlier (Fig. S4 and S7HI). Comparing the promoter on-rates shown in Fig. 5C and 5D, we note that the typical magnitude of change in promoter-on rates (a few-fold) is very small compared to the full range of *k*_*i*_ across genes (> 3 orders of magnitude); see also Fig. S11GH. Hence, transcriptional regulation, which is itself the main source of gene regulation, offers a comparatively small change to protein abundances compared to the vast innate differences in promoter on-rates.

This point appears to extend well beyond C-limited growth which we have been focused on here due to the limited availability of all data sets (mRNA fluxes and degradation rates). Looking over the proteome of cells under A-limited growth and translation limited growth, we see the same theme of moderate changes across condition vs. vast differences across genes (compare Fig. S13BC with Fig. S13A). The same is seen for cells grown under osmotic stress and in rich medium (Fig. S13D, E). We repeated this comparison across 19 different growth conditions for which absolute proteomic abundance data are available for our strain [11], and computed the maximum fold-differences in protein fractions between any 2 out of the 19 conditions for >1400 proteins, giving a sense of the magnitude of their regulation. A scatter plot of maximum fold-change vs. median protein abundance for these genes (Fig. 5E, gray points) show that across the >3-order of magnitude difference in the median abundance of these proteins, the maximum variability across the 19 conditions is very limited (67% of proteins change at most by 5-fold). Thus, regulatory effects seem to be limited generally compared to differences in the magnitude of expression, which is largely set by the innate promoter characteristics as shown in Fig. 5C.

Interestingly, proteins encoded by essential genes, as well as proteins associated to “housekeeping” functions such as DNA replication and cell membrane/wall biosynthesis display remarkably small change across the 19 conditions tested (green points and histograms in Fig. 5E; see also Fig. S12F-J), suggesting that the expression of essential genes and house-keeping genes are almost entirely set innately. Indeed, some of these genes (*dnaA, ftsZ, polA*) are among the ones with smallest variation in promoter on-rates in carbon-limited conditions (Fig. S10D-F), despite the presence of known regulators [41, 42]. On the contrary, the expression of proteins involved in many metabolic functions (TCA cycle, nutrient uptake, biosynthesis) can change strongly across conditions (Fig. S12K-O).

Quantitative determination of the promoter on-rates across conditions allows us delve more deeply into transcriptional control. As an example, different growth-rate dependences of the promoter on-rate are found for the set of proteins that are up-regulated under C-limitation (Fig. S13), suggesting different regulatory effects. A group of genes (*acs* and operons encoding various sugar uptake systems, Figure S13A-D) showed increase in promoter on-rate as growth rate decreases, reflecting the well-known regulation by cAMP-Crp in carbon limited growth [9, 43]. In contrast, the promoter on-rates of genes encoding TCA enzymes tend to have non-monotonic dependence on growth rate, peaking at intermediate growth rate of 0.7-0.8/h (Fig. S13E-G). This dependence is consistent with the result of multiple regulatory factors acting on these promoters, including the aforementioned effect of cAMP-Crp, as well as the opposing effect of repression by ArcA, whose level also increased under C-limitation (Fig. S13H). More intriguingly, a group of genes exhibited decrease in promoter on-rate despite increase in protein levels. These include *sodB* and *proP* (Fig. S13IJ). While the promoters of these genes are reported to be acted upon by various regulators, what they have in common is repression by cAMP-Crp [9, 43]. The three types of regulatory patterns are summarized in Fig. 5F.

For proteins involved in translation, although their abundances generally increase at increasing growth rates, the magnitude of the change shows quite some differences (black symbols, Fig. S14A-D), despite the fact that many of these genes are co-transcribed [44]. Reassuringly, these differences largely disappear when considering promoter on-rates, as summarized in Fig. 5G (green and light blue triangles). Moreover, the growth rate dependence of these promoter on-rates is also similar to that of the on-rates of the rRNA operons (blue crosses), obtained from the rRNA synthesis flux and the measured elongation rate of RNAP on the *rrn* operons [33]; see SI Methods. These results suggest the existence of post-transcriptional effects acting on ribosomal proteins and elongation factors. This is well known for ribosomal proteins [45] but largely uncharacterized for elongation factors. On the other hand, a number of other translation-related genes, e.g., those encoding tRNA synthases and translational initiation factors (Fig. S14E-H), exhibit similar growth-rate dependences at the protein level despite being organized in different transcriptional units. Plotting the promoter on-rate of the latter genes indeed shows different growth-rate dependencies (red symbols, Fig. S14E-H), suggesting that their protein levels are primarily shaped by post-transcriptional effects. Overall, the promoter on-rate for translational genes increase with the growth rate, and this increase largely accounts for the increase in the total promoter on-rate 𝒦 (Fig. 5H).

### Control of global mRNA synthesis by the anti-sigma factor Rsd

Having explored the consequences of the quantitative central dogma, Eq. (12), we next return to the global constraint between transcription and translation, Eq. (11). The combination [*Ori*] · 𝒦 in Eq. (11) has some moderate growth-rate dependence as shown in Fig. 6A. The concentration of available RNAP, estimated as the ratio of the mRNA synthesis flux (*J* _mR_) and [*Ori*] · 𝒦 based on Eq. (6) and (10), exhibits a stronger growth rate dependence (left axis in Fig. 6B), approximately matching that of the concentration of active ribosomes (right axis).

**Figure 6.**
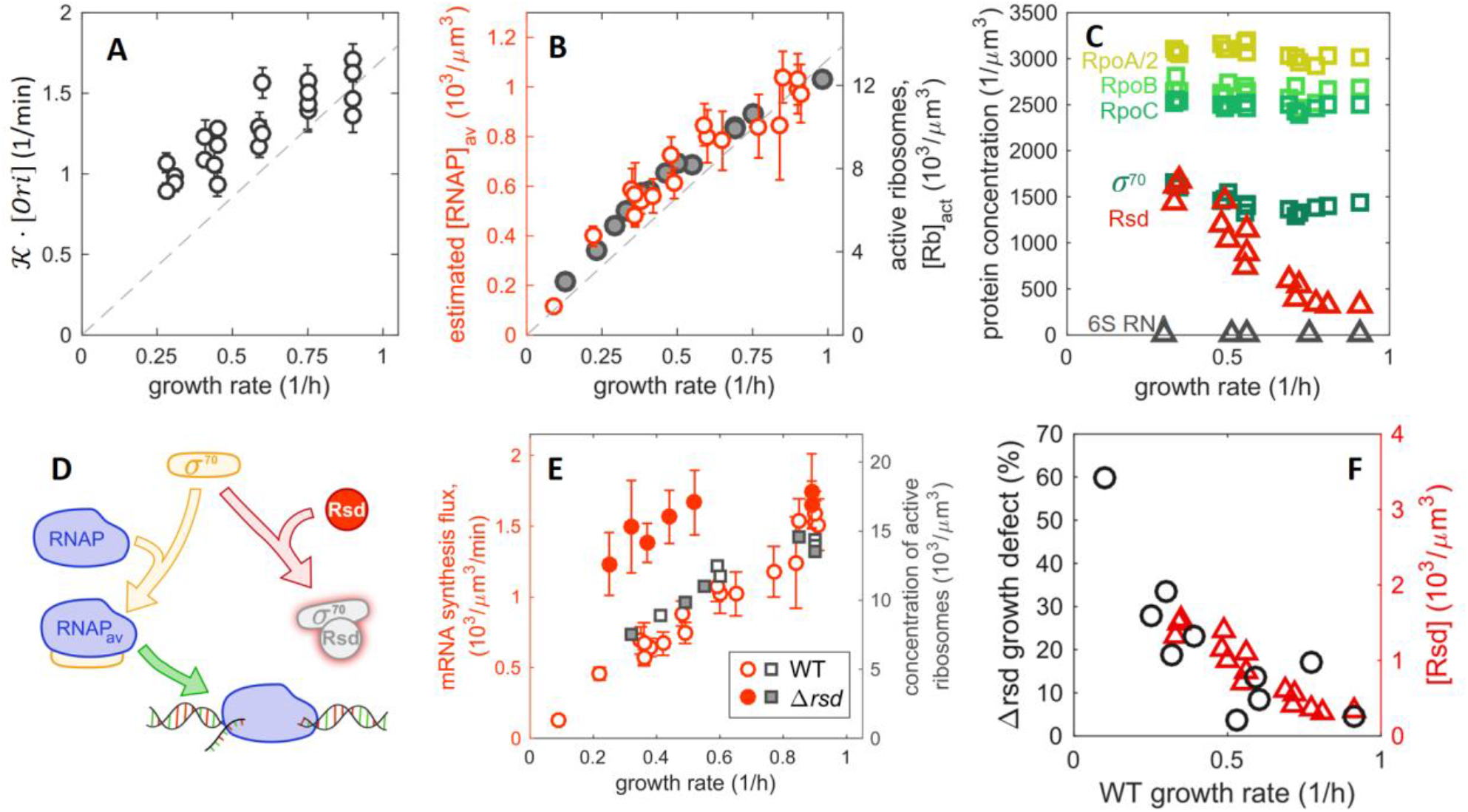
The role of the anti-sigma factor Rsd in global regulation of mRNA synthesis. **(A)** Value of 𝒦 · [*Ori*] across growth rates, obtained from the values of the Ori concentration and the total regulatory activity 𝒦 shown in Fig. 4A and 4G, respectively. For comparison, the dashed line shows direct proportionality to the growth rate. **(B)** Concentration of available RNA polymerases (red symbols, left axis), estimated from the ratio between the measured mRNA synthesis flux, Figure 3G, and 𝒦 · [*Ori*] (using the interpolated curves in Figures 4A and 4G). Note that this quantity shows a stronger dependence on the growth rate compared to 𝒦 · [*Ori*] in panel (A) and has the same growth-rate dependence as the concentration of active ribosomes (grey symbols, right axis). **(C)** The concentrations of various components of the transcription machinery in carbon-limited conditions is plotted against the growth rate. Components of the core enzyme, RpoABC, and the major sigma factor *σ*^70^ are shown as squares. Known modulators of *σ*^70^, Rsd and 6S RNA are shown as triangles. The protein concentrations are determined from mass spectrometry [11], while the concentration of 6S RNA is determined from RNA-sequencing and the concentration of total mRNA concentration (Fig. S4). **(D)** Cartoon illustrating the control of RNA polymerase (RNAP) availability through the known *σ* ^70^-sequestration function of Rsd [48, 74]. **(E)** Comparison of mRNA synthesis fluxes between wild type (open symbols) and *Δrsd* strain (filled symbols). Left axis: total mRNA synthesis flux of *Δrsd* strain (red filled circles) and wild type (red open circles). Right axis: concentration of active ribosomes computed from the measured total RNA for the two strains and the fraction of active ribosomes observed in carbon limited growth [7]. **(F)** The growth defect of *Δrsd* strain, defined as % reduction in growth rate compared to wildtype cells in the same growth condition (black circles, left axis), is plotted against the growth rate of wild type cells for the range of carbon-limited growth conditions. The observed growth reduction matches Rsd expression of wild type cells in the same conditions (red triangles, right axis; same data as in panel C).

A simple scenario explaining the change in the availability of RNA polymerases is one in which the abundance of the transcription machinery itself changes accordingly. However, our quantitative proteomics data shows that the cellular concentrations of the RNA polymerase components, including the house-keeping factor *σ*^70^ (encoded by the gene *rpoD*), are all maintained constant across the growth rate range studied (Fig. 6C, squares). We next checked the expression levels of the two known modulators of *σ*^70^ function, 6S RNA [46, 47] and the anti-*σ*^70^ protein Rsd [48–50]. While the concentration of 6S RNA is 100-fold lower than that of *σ*^70^ (grey triangles, Fig. 6C) and thus unlikely to affect the global transcription flux in these conditions, the Rsd concentration rises to levels comparable to that of *σ*^70^ as growth rate is reduced (Fig. 6C, red triangles). This raises the possibility that Rsd is a regulator of global transcription by sequestering *σ*^70^ during exponential growth (Fig. 6D), even though Rsd is commonly thought to play its role in stationary phase [48, 51]. We tested this scenario by characterizing the total mRNA synthesis flux in a Δ*rsd* strain. Indeed, the mRNA synthesis flux becomes nearly independent of the growth rate (Fig. 6E, filled red circles), well exceeding that of the wild type strain (open red circles), especially at slow growth where Rsd is highly expressed. Without *rsd*, the mRNA synthesis flux is no longer matched to the translational capacity (compare filled symbols), in contrast to the tight matching observed in wild type strain (open symbols). Concomitantly, theΔ*rsd* strain exhibits a growth defect that is proportional to the level of Rsd expression in wild-type cells in slow growth conditions where Rsd is expressed (Fig. 6F). Given the approximate constancy of mRNA turnover across growth conditions for wild type cells (Figure S7E-G), the data in Fig. 6E suggest that Rsd is centrally involved in controlling the total mRNA levels (Figs. 2A and 3F).

## DISCUSSION

Proteome allocation in bacteria is known to adjust in quantitative measures in accordance to different functional demands in different growth conditions and for different genetic perturbations [6, 9, 52, 53]. Yet, how changes at the proteome level arise from the underlying regulatory processes is not clear. In particular, it is not clear how transcriptional and post-transcriptional processes accommodate global constraints governing cellular protein dynamics, i.e. a fixed total protein density (Fig. S2F, [8]) and a limited translation rate for protein synthesis by each ribosome [7].

### Simple rules governing promoter and mRNA characteristics

In this work, through comprehensive transcriptomic and proteomic studies, complemented by quantitative measurement of total mRNA abundance and transcription flux, we determined the absolute mRNA and protein abundances, as well as mRNA degradation rates and promoter on-rates, for 1500+ genes in *E. coli* for many growth conditions during steady-state growth (Table S2, S4). The results lead us to identify two simple rules on the promoter and mRNA characteristics which profoundly shape how *E. coli* responds to environmental changes while coping with global constraints: (1) promoter on-rates span over 3 orders of magnitudes across genes, but vary much less (∼3-fold) across conditions for most genes. Thus, each gene is expressed within an innate abundance range across conditions, e.g., with ribosomal genes belonging to the most abundant and DNA replication proteins, belonging to one of the least abundant classes. (2) The mRNA characteristics, including translation initiation rate and mRNA degradation rate, vary little (<2-fold for half of genes) across genes and conditions. The initiation rates are sufficiently rapid to maintain a high density of ribosomes on the mRNA, resulting in high translational burstiness despite short mRNA half-lives.

Our results suggest a limited effect of post-transcriptional regulation on gene expression across the set of exponential growth conditions examined, even though such effects can be seen for specific perturbations on some genes (Fig. S4, S7HI). Furthermore, the mRNA characteristics are seen to be similar across genes even for a given condition, despite widely studied mRNA-specific effects arising from, e.g., ribosome binding sequence, codon usage, secondary structure [54–59]. For example, different choices of the ribosome binding sequence can easily modulate the protein expression level by several orders of magnitude [60], results which have been widely exploited in synthetic biology applications [61]. Nevertheless, the translation initiation rates are narrowly distributed for most endogeneous genes (Fig. 2CD). Consequently, mRNA levels accurately reflect protein levels (Fig. 1BC, Fig. S3), and the wide range of protein levels observed is predominantly set by the widely different promoter strengths (Fig. 5C), i.e., Rule #1 above. The latter point can actually be seen already in the data of Keren *et al* [62], which exhibited a broad range of activities for different promoter in a given condition through a genome-wide promoter-gfp fusion study, even though the focus of that study was on small changes across conditions, and the connection between promoter activities and protein levels was lacking. Only by quantitatively characterizing the transcriptome and proteome for many conditions are we able to quantitatively connect protein levels to promoter activities, and moreover, separate contributions to promoter activities due to the promoter on-rates (Fig. 5) and available RNA polymerase (Fig. 6).

The rules governing promoter and mRNA characteristics deduced here dictate, to a large extent, *E. coli*’s strategy to implement gene regulation while complying with the constraints on a fixed total protein concentration and a limited translation capacity. This can be cast into two principles of gene expression as summarized below.

### Global coordination between transcription and translation

The similarity of mRNA characteristics across genes and conditions (Rule #2) implies a constant density of translating ribosome on most mRNA (∼4 ribosomes/mRNA, Fig. 2A) across growth conditions. Since the total concentration of translating ribosomes gives the total protein synthesis flux (given limited translational elongation rate across nutrient conditions [7]), the total mRNA concentration must match the total protein synthesis flux, with the latter varying linearly with the growth rate due to the constraint on total protein density [8] (Fig. S2). As the total mRNA pool is specified by the total mRNA synthesis rate (given the constant mRNA turnover rate across conditions), the cell is forced to balance total mRNA synthesis with total protein synthesis, a crucial condition captured by Eq. (11). We refer to this balance as the *principle of transcription-translation coordination*. We showed that *E. coli* implements this coordination across nutrient conditions primarily by adjusting the available RNAP concentration via the anti-*σ*^70^ factor Rsd (Fig. 6D-F).

If this coordination is broken, as in the case of Δ*rsd* mutant (Fig. 6E), then Rule #2 must be broken as long as the constraints on translation capacity and protein density hold. An oversupply of mRNA (with respect to the ribosomes) is expected to decrease the rate of translation initiation (due to competition for limited ribosomes) and/or increase the rate of mRNA degradation (due to reduced protection of mRNA by elongation ribosomes against RNase activity [63, 64]. Aside from the futile cycle involving the synthesis and degradation of useless mRNAs and affecting growth (Fig 6F), breaking Rule #2 would significantly complicate the otherwise simple relation between transcriptional regulation and protein concentrations of the wild type system to be discussed below.

The growth rate dependence of the available RNAP in wild type cells actually affects how transcriptional regulation should be quantified. In the literature, a common measure of transcriptional regulation is the so-called “promoter activity”, defined by the protein synthesis flux obtained as the product of growth rate and concentration of the protein of interest [65]. It is commonly assumed that this promoter activity, which reflects the transcription initiation rate (SI Note), captures the effect of transcriptional regulation. However, since the rate of transcription initiation is dependent on the concentration of available RNAP, and the latter changes with growth condition due to titration by Rsd, promoter activity mixes the global effect due to RNAP availability and gene-specific regulatory effects. As we will discuss below, a much more direct connection exists between protein concentrations and the promoter on-rates (instead of the promoter activities).

### The predominant role of transcription in setting protein concentrations

The similarity of mRNA characteristics (Rule #2) together with the vast disparity of promoter characteristics (Rule #1) across genes in a given condition implies that protein abundances are predominantly set by the promoter characteristics, specifically, the promoter on-rates. Furthermore, as mRNA characteristics remain similar across different growth conditions (Rule #2), changes in protein concentrations across conditions must arise primarily from changes in the promoter on-rates, i.e., via transcriptional regulation (Fig. 5D). We refer to this strong effect of transcription on gene expression as the *principle of transcriptional predominance*.

Given that the promoter on-rates change by only a few-fold across growth conditions for most genes while they differ by up to 3-4 orders of magnitude across genes, we can conclude that the abundances of most proteins are set by the basal promoter on-rate. In this light, transcriptional regulation is seen largely to fine-tune protein abundances which are set in different abundance classes by the “basal level” of their promoters. Fig. S12 provides a cursory look of the different abundance range proteins in different functional classes belong to. They reflect innate differences in the functional demands for growing cells, e.g., the demands for co-factor synthesis and tRNA charging are much smaller than those for ribosomes and glycolysis. An in-depth analysis of promoter sequences for these different abundance classes may lead to new understanding of the sequence determinant setting the wildly different basal expression levels. It will also be interesting to see whether the classes of abundance distribution found here for *E. coli* is conserved across microbes.

We next turn to gene regulation, which as mentioned above, becomes largely reduced to the “fine-tuning” promoter on-rates via transcriptional regulation. Although setting protein levels transcriptionally appears simple at a qualitative level, quantitative relation between promoter on-rates and protein concentrations is complicated by the constraint on total protein density: The two cannot be simply proportional to each other as commonly assumed, since total protein density is a constant across growth conditions while sum of promoter on-rates is generally not constrained. As indicated by the conundrum described in Introduction, if a pleiotropic transcriptional activator doubled the promoter on-rate of each gene, it cannot result in the doubling of every protein because the total protein density is fixed. The resolution to this conundrum, as our study revealed, is that as long as mRNA characteristics are similar, the promoter on-rate of a gene as a fraction of the total promoter on-rate (), is equal to the fractional mRNA and protein abundances; see Eq. (12). Thus, a doubling of all promoter on-rates will have no effect on mRNA and protein concentrations, because it does not change the on-rate of one promoter relative to another. Although the appearance of total promoter on-rate in Eq. (12) couples transcriptional regulation globally, with non-intuitive results discussed below, the simplicity of the form is still striking. In particular, this relation entirely bypasses the activities of the macromolecular machineries (RNAP and ribosomes) which underly gene expression, as long as these machineries are coordinated according to the principle of transcription-translation coordination. We view Eq. (12) as a quantitative formulation of the Central Dogma, providing a quantitative link from DNA to RNA and proteins.

### Global transcriptional coupling and its consequences

Since the protein output of a given promoter depends on the total rate 𝒦 in Eq. (12), non-intuitive relations between promoters and protein concentration can arise whenever 𝒦 changes across conditions. The latter is almost guaranteed whenever there is a substantial change in growth rate, since the abundance of ribosomal proteins change with growth rate, and ribosomal proteins belong to the most abundant protein class, and changes in their on-rate exert a significant effect on the total on-rate 𝒦 (Fig. S12H, 5H). Fortunately, changes in the total rate 𝒦 can be deduced without the need to measure the on-rate of each gene. As illustrated in Fig. 4FG, 𝒦 can be deduced from quantifying the expression of constitutively expressed genes together with the knowledge of gene dose.

Given the unavoidable changes in the total rate 𝒦 across growth conditions, it is generally dangerous to infer regulatory activities directly from changes in mRNA and protein levels; see the simple examples illustrated in Fig. 4E. This effect of global coupling would hardly affect the result of most classical studies, which typically involved changes of many tens-fold in the output of individual promoters, much larger than the few-fold change in 𝒦 expected across conditions (see e.g., Fig. S11H for C-limited growth). However, this global effect has to be taken into account when examining genes changing by less than a few-fold in expression, which is the case for the vast majority of genes (e.g., 60% of genes below 2-fold for C-limited growth; see Fig. 5G). One way to counter this global effect is to keep growth rate constant during study of gene regulation. This however cannot be done for many physiological studies which intrinsically involve different growth conditions. In this case, the best resort may be to incorporate constitutively expressed reporter genes as internal reference point, as was done in this study.

Our work provides a quantitative framework for researchers examining the expression of individual genes to distill gene-specific regulatory effects from global interactions. The knowledge of the promoter on-rates *k*_*i*_ for individual genes offers a direct, promoter-centric view of regulation across conditions at the genome-scale (Fig. 5). This knowledge will be invaluable in deciphering endogenous genetic circuits, as well as in guiding the design of synthetic circuits operating in different growth conditions [66–68]. The entangled relationship between promoter on-rates and protein levels described in this work stands in stark contrast to the very simple mathematical relations (‘growth laws’) between protein levels and the cell’s growth rate established in earlier studies [6, 9, 10]. For example, the concentrations of ribosomal proteins and catabolic proteins vary linearly with the growth rate (grey circles in Fig. S13A-D and Fig. S14A), while the promoter on-rates show manifestly nonlinear dependences (colored triangles in Fig. S13A-D and Fig. S14A). These results indicate that signals driving the expression of these proteins (e.g., cAMP for catabolic proteins [69] and ppGpp for ribosomal proteins [44]) are not simply reflecting the growth rate; instead, these signaling pathways likely implement autonomous feedback strategies that peg the concentrations of key proteins in simple relation to the growth rate [70, 71].

It will be interesting to extend the current analysis of gene expression to include additional growth conditions (e.g., under stress) as well as to other bacterial species, to see the generality of the rules and principles uncovered here. We note that protein degradation, which has been neglected in the analysis here in comparison to growth-mediated dilution, will play increasingly more important roles for slow-growing cells. We caution that the results described here are specific to bacteria. Eukaryotes are subjected to many more complex post-transcriptional processes; e.g., proteins turnover not only by the well-established ubiquination pathways, but also by processes such as autophagy which turnover substantial fractions of the entire cell [72]. Global constraints are also less understood, in particular the extent to which protein density may vary across conditions. Indeed, even quantifying the cell volume may be highly nontrivial as large portions within a cell may be occupied by sub-cellular compartments (e.g., vacuoles) that do not contribute to the cytosol. Nevertheless, detailed characteristics of gene regulatory processes obtained from this study, along with the insights they reveal on the design principles of bacterial gene expression, suggests that similar feat may be accomplished by performing accurate quantitative measurements for eukaryotic processes, and combining them with quantitative characterization of cell physiology across conditions.

## ACKNOWLEDGEMENTS

We thank Kapil Amarnath, Ralf Bundschuh, Kristen Jespen, Chenli Liu, Hiroyuki Okano and Hai Zheng for helpful discussions, and Irina Artsimovitch, Kurt Fredrick, Hernan Garcia, Luca Gerosa, Ido Golding, Michael Ibba, Karl Kochanowski, Namiko Mitarai, Mary-Ann Moran, Rob Phillips and Matt Scott for critical comments on the manuscript. RNA sequencing was performed at the Institute for Genomic Medicine at UC San Diego. This work was supported by NSF through grant MCB1818384 and NIH through grant R01GM109069.

## AUTHOR CONTRIBUTION

R.B., M.M., T.H. conceived the study, interpreted results and wrote the manuscript. R.B. designed and performed the experiments. M.M. performed the computational and mathematical analysis of the data. Z.Z. generated the strains. I.S. developed the RNA-seq analysis pipeline. M.M., R.A. and C.L. provided and analyzed proteomic data. T.H. supervised the work and acquired funding.

## DATA AVAILABILITY STATEMENT

The proteomic mass spectrometry files as well as data analysis files have been published previously (Ref. [11]) and are accessible through the ProteomeXchange Consortium via the PRIDE partner repository: http://www.ebi.ac.uk/pride/archive/projects/PXD014948. The used *E. coli* spectral library for DIA/SWATH data analysis is available via SWATHAtlas: http://www.peptideatlas.org/PASS/PASS01421. The genome-wide parameters of gene expression generated in this study, including transcription and translation initiation rates, mRNA degradation rates, promoter on-rates, as well as protein, mRNA and gene concentrations for reference condition (glucose minimal medium) and slow glucose-limited growth are reported in Table S4. Additional analysis, including modelling and details on the calculation of the parameters, are included in Supplementary Notes S1 to S5. Raw .fastq files for RNA-sequencing and Supplementary Tables S2-5 are not included in this biorxiv submission. These data are available from the corresponding author (T.H.) upon reasonable request.

## DECLARATION OF INTERESTS

The authors declare no competing interests.

## Notes

### Competing Interest Statement

The authors have declared no competing interest.

### Summary of Updates

This version includes updated reference for Mori et al (2021) and additional descriptions of the proteomics dataset used.

