## Supplemental Information for "Principles of gene regulation quantitatively connect DNA to RNA and proteins in bacteria"


| Section | Page |
|---|---|
| **Supplemental Figures** | 2 |
| **Supplemental Tables** | 28 |
| **Supplemental Methods** | 32 |
| **Supplemental Notes** | 38 |
| **Supplemental Note S1:** Unit Conversions | 38 |
| **Supplemental Note S2:** Protein homeostasis and ribosome density | 44 |
| **Supplemental Note S3:** mRNA homeostasis | 48 |
| **Supplemental Note S4:** Promoter on-rates and quantitative Central Dogma relation | 50 |
| **Supplemental Note S5:** Determination of the absolute concentration of available RNAP | 55 |
| **Supplemental Information References** | 58 |

### SUPPLEMENTAL FIGURES

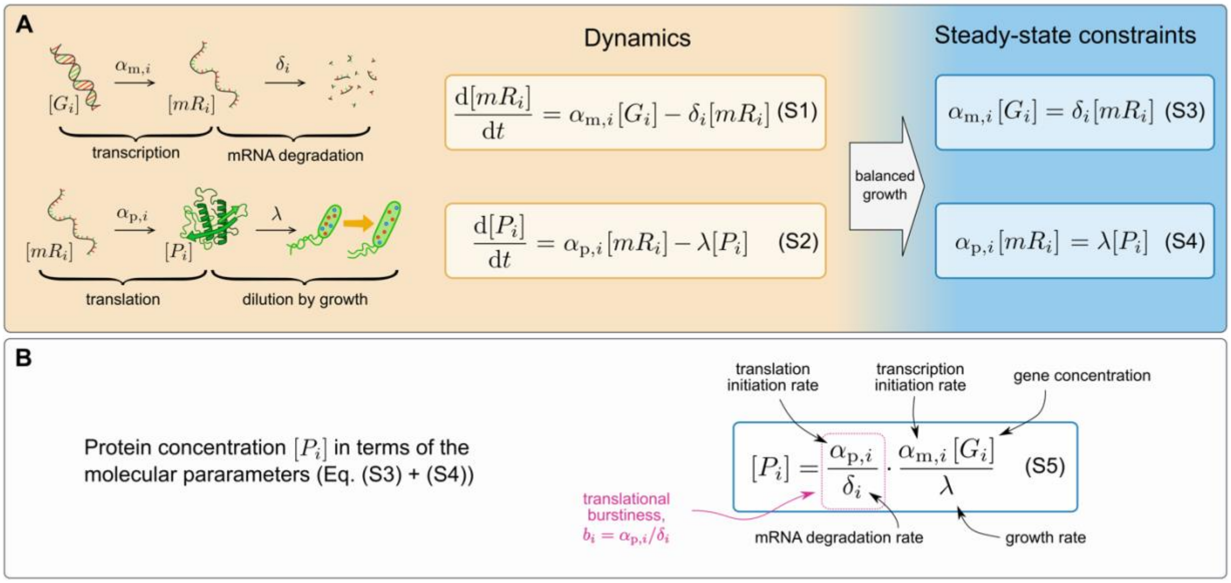


**Figure S1. Models of transcription and translation.**
(A) A widely used framework to model gene expression in bacteria uses simple rate equations describing the production and degradation/dilution of mRNAs (top) and proteins (bottom) [1, 2]. The mRNA concentration of gene $i$, $[mR_i]$ depends on its transcription initiation rate $\alpha_{m,i}$, the gene concentration $[G_i]$, and its rate of degradation, $\delta_i$, as described by Eq. (S1). The concentration of the corresponding protein, $[P_i]$, depends on its translation initiation rate $\alpha_{p,i}$, its mRNA concentration, and the rate at which the protein is diluted due to cell growth $\lambda$, Eq. (S2). In fact, protein turnover is negligible in for exponentially growing *E. coli* except for a few proteins known to be targeted by proteases [3, 4]. In balanced exponential growth, the concentrations of genes, mRNAs and proteins are time-independent, giving rise to the steady state relations (S3) and (S4).
(B) Equations (S3) and (S4) can be combined to obtain the protein concentration $[P_i]$ in terms of the rate parameters (S5), which is widely used in computational models of gene expression. In this expression, the ratio $\alpha_{p,i}/\delta_i$ quantifies the number of proteins produced by each mRNA during its lifetime, and is referred to as the translational burstiness $b_i$. Equation (S5) is not useful to predict gene expression from the microscopic parameters across growth conditions or after major perturbations of gene expression, e.g. induction to high levels of LacZ proteins [5]. For example, according to Eq. (S5), doubling all transcription initiation fluxes $\alpha_{m,i}$ would lead to a doubling of all protein concentrations $[P_i]$, which is not realistic given the limited density of the cell. This suggests the existence of global constraints among the above described rates compatible with a constant protein concentration.

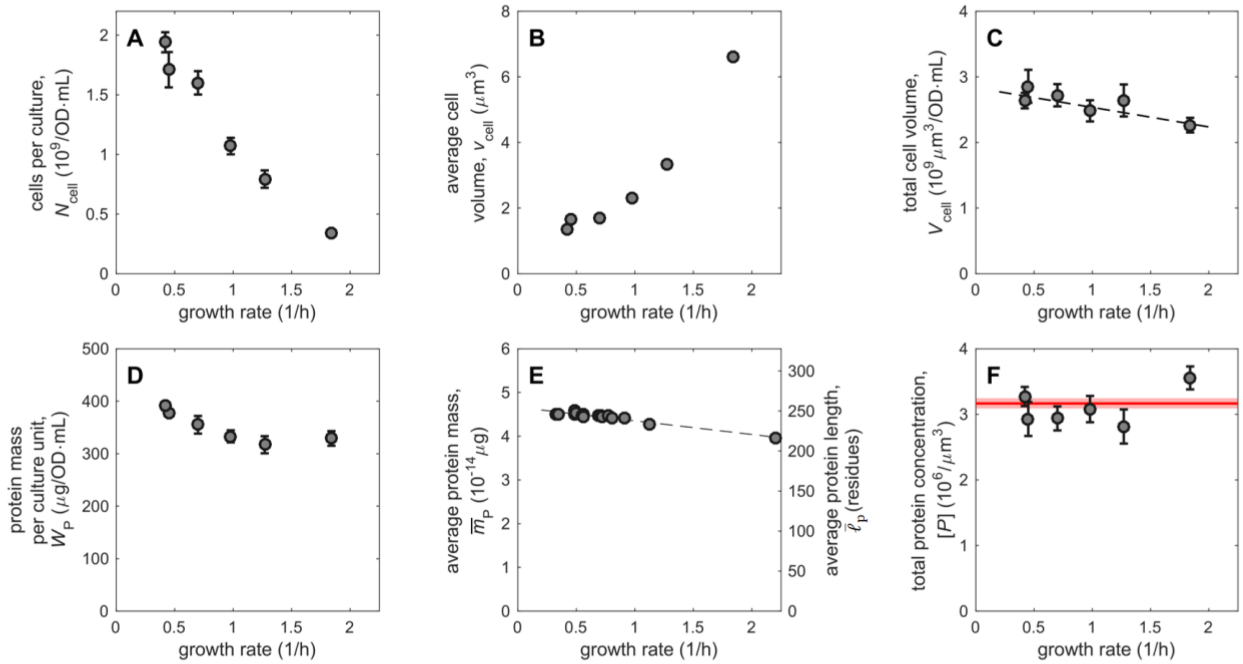


**Figure S2. Cellular volume and protein concentrations.**

**(A)** Number of cells $N_{\text{cell}}$ per culture volume (OD·mL), as determined from Basan et al. [6] for *E. coli* NCM3722 cells under exponential growth in various nutrient conditions; x-coordinate shows the growth rate of the culture.

**(B)** Average volume of individual cells, $v_{\text{cell}}$, as obtained in Basan et al [6] for the same growth conditions as in (A); error bars (standard deviation of the avg. cell volume) are too small to be visible.

**(C)** The total cell volume $V_{\text{cell}}$ per culture volume is obtained as the product of cell number per culture unit and the average cell volume, i.e., $V_{\text{cell}} \equiv N_{\text{cell}} v_{\text{cell}}$; error bars are propagated from those of $N_{\text{cell}}$ and $v_{\text{cell}}$. We observe that $V_{\text{cell}}$ changes only mildly with the growth rate, compared to the strong dependence on growth rate observed for both $N_{\text{cell}}$ and $v_{\text{cell}}$. This quantity is well described by a linear fit $V_{\text{cell}} = a + b\lambda$ (dashed line), with $a = 2.83 \cdot 10^9\ \mu\text{m}^3/\text{OD}/\text{mL}$ and $b = -0.3 \cdot 10^9\ \mu\text{m}^3\text{h}/\text{OD}/\text{mL}$. We used this fit to convert abundances per culture volume (OD·mL) into abundances per cell volume (concentrations) across growth rates.

**(D)** Total protein mass $W_{\text{P}}$ per unit culture volume in the same growth conditions as those shown in (A) and (B); data from ref. [6].

**(E)** Average protein mass $\overline{m}_{\text{P}}$ (left axis) and protein length $\overline{\ell}_{\text{P}}$ (right axis) in carbon limited conditions, estimated from the known length of individual proteins and the proteome composition of *E. coli* grown in glucose titration (C-limitation, growth rates below 1/h), G6P+glucorate (growth rate 1.12/h) and LB medium (growth rate 2.2/h) obtained from Ref. [7] (see Note S1). Throughout this work we approximate protein masses as $m_{\text{P},i} = m_{\text{aa}} \ell_i$ where $m_{aa} = 1.83 \cdot 10^{-16}$ μg is the average residue mass. The black dashed line indicates a linear fit $\overline{m}_{\text{P}} = a + b\lambda$ with $a = 4.66 \cdot 10^{-14}$ μg and $b = -0.31 \cdot 10^{-14}$ μg · h.

**(F)** Total number of proteins per cell volume, $[P] = W_{\text{P}}/\overline{m}_{\text{P}} V_{\text{cell}}$, obtained by combining the total protein mass per culture volume in (D), the total cellular volume in (C) and the average protein mass in (E). Error bars were propagated from $W_{\text{P}}$ and $V_{\text{cell}}$, while $\overline{m}_{\text{P}}$ was linearly interpolated at the same growth rates of the other quantities. The result is approximately constant across growth rates, $[P] = (3.15 \pm 0.09) \times 10^6/\mu\text{m}^3$ (weighted average, in red).

In this work, we refer to the number of proteins per cell volume as "protein concentration". Note that this measure of protein concentration differs in absolute value from alternative definition according to the number of proteins per water volume, because a non-negligible portion of the cell volume is occupied by proteins and other cellular components. However, as the buoyant density of *E. coli* is almost independent of the growth rate [8], we expect the ratio of the cellular water volume and cell volume to be proportional, so that protein concentration is nearly independent of the growth rate for both definition of concentration. Finally, given that the total protein concentration is constant, the protein number fractions $\psi_{\text{p},i} \equiv [P_i]/[P]$, which are directly obtained from mass spectroscopy [7], are directly proportional to protein concentrations independent of growth rates.

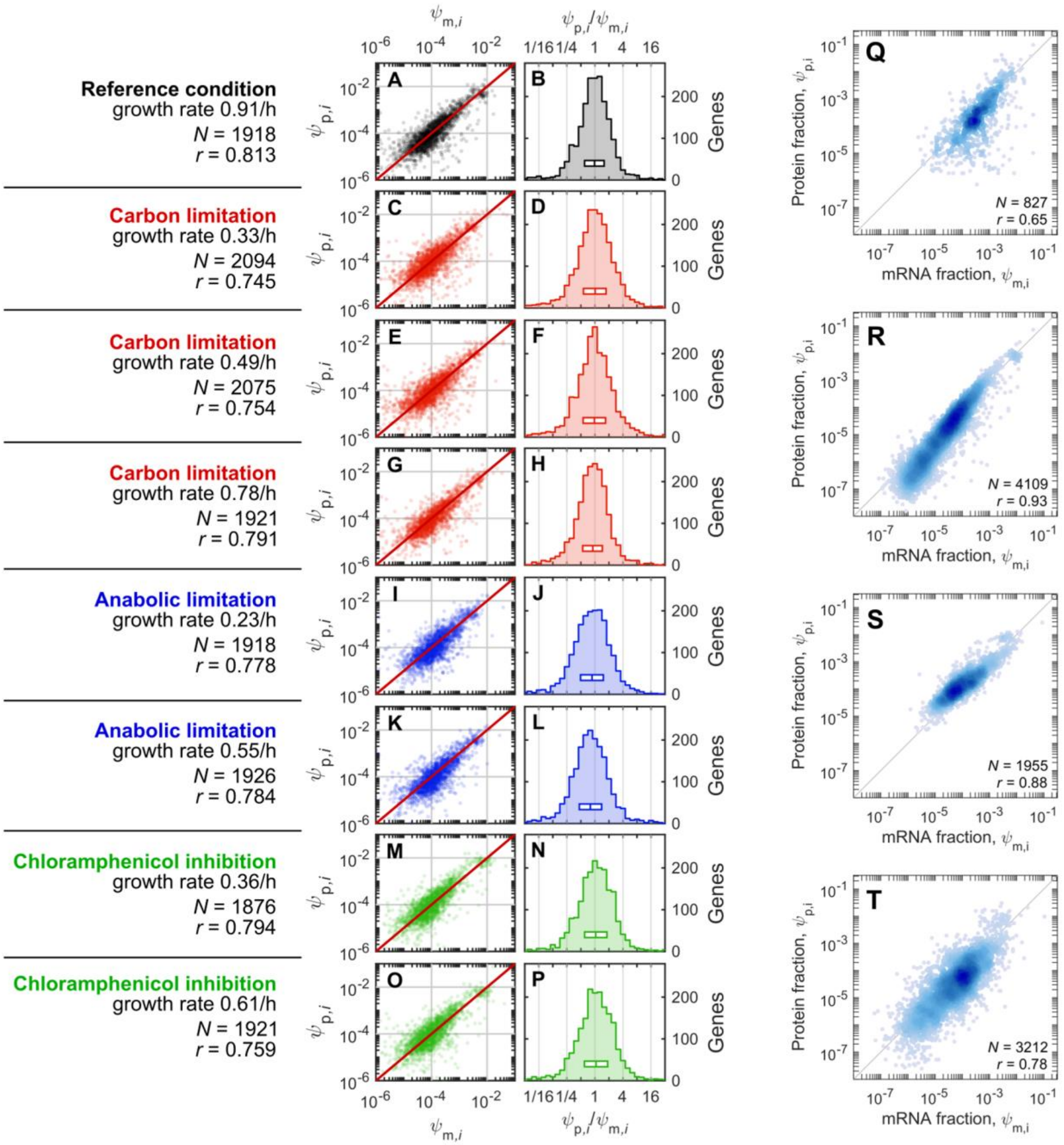


**Figure S3. Comparison of protein vs mRNA abundances.**
**(A-P)** Absolute number fractions of proteins, $\psi_{\mathrm{p},i}$ and mRNAs $\psi_{\mathrm{m},i}$, generated from quantitative mass spectrometry (DIA/SWATH, [7]) or RNA sequencing (RNA-seq; see SI Methods). Number fractions are the natural output of both quantitative proteomics methods (e.g. iBAQ [9] or xTop [7]) and transcriptomics (usually expressed as reads per kilobase per million mapped reads, RPKM), since in both cases the raw output is proportional to either protein or mRNA copy numbers in the sample of interest. In particular, the number fractions are equal to the cellular concentration of the protein/mRNA of interest, divided by the total protein/mRNA concentration, i.e. $\psi_{\mathrm{p},i} = [\mathrm{P}_i]/[\mathrm{P}]$ and $\psi_{\mathrm{p},i} = [\mathrm{mR}_i]/[\mathrm{mR}]$. Here, the scatter plot of protein and mRNA fractions (left) or the distribution of their ratio $\psi_{\mathrm{p},i}/\psi_{\mathrm{m},i}$ (right column) are shown for a variety

of growth conditions: glucose minimal medium (AB), carbon limitation (red, C-H), anabolic limitation (blue, I-L), translation limitation (green, M-P). The growth condition, growth rate ($\lambda$), number of genes plotted ($N$) and Pearson correlation coefficient ($r$) are indicated in the left-most column next to the scatter plots; boxes in the histograms show the median value of the ratio together with the 25-75% percentile interval. The proteomics and transcriptomics dataset used in this work can be found in Table S2.

**(Q-T)** The fractional abundances for proteins are plotted against those of mRNAs using dataset available in the literature. The conversion of these published datasets into fractional number abundances (Note S1), facilitates their direct comparison with the vast range of data analyzed in this work. **(Q)** Protein abundances from a YFP fusion library and RNA-seq for *E. coli* cells grown in M9 medium with glucose, amino acids and vitamins [10]. **(R)** Protein number fractions estimated from protein synthesis rates obtained from ribosome profiling (SI Note S1 for conversion) are plotted against RNA-seq data for cells grown in MOPS complete media [11]. **(S)** Same as panel (R), but for cells grown in MOPS complete media [12]. **(T)** Same as panel (R), using the data in Ref. [13].

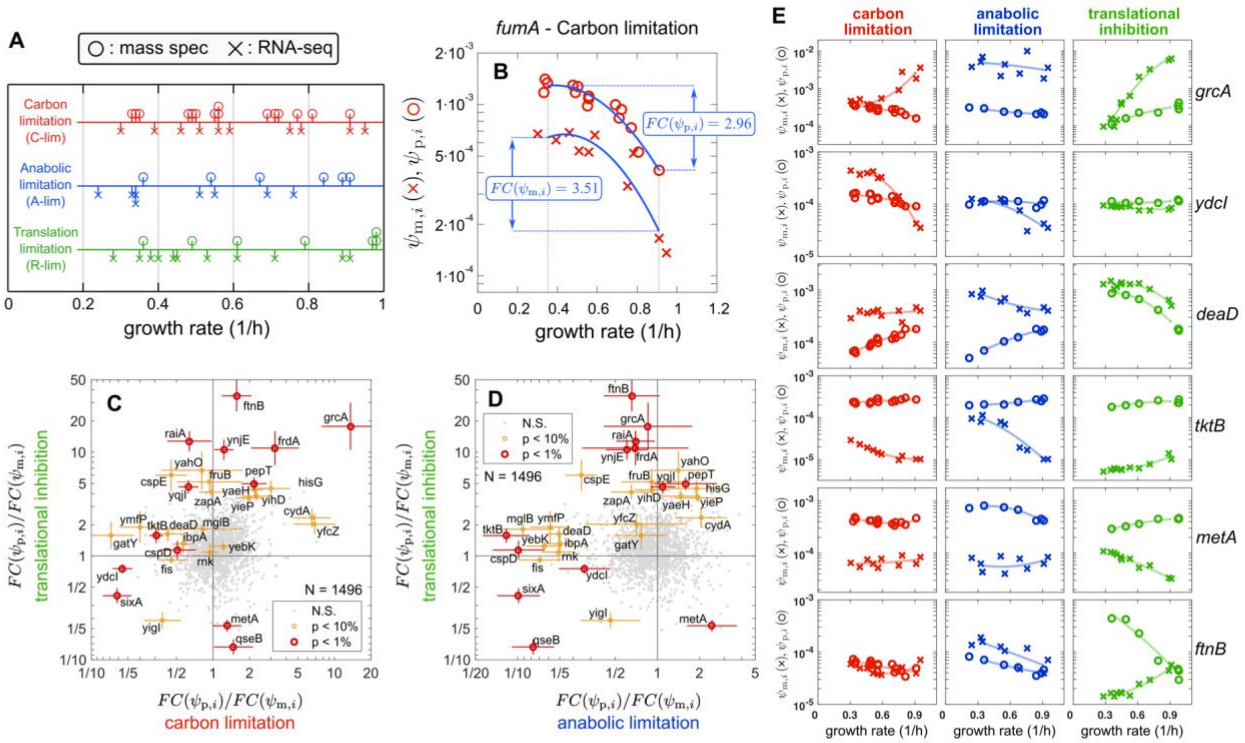


**Figure S4. Genome-wide comparison of protein and mRNA fold-changes across growth conditions.**

**(A)** Illustration of the complete set of proteomic and transcriptomic data used in this work. In total we obtained 29 mass-spec samples and 31 RNA-seq samples for different growth rates ranging from about 0.3/h to 0.9/h by either limiting carbon catabolism (red), anabolism (blue) or translation (green). Circles indicate the growth rates of the samples for which various mass spectrometry [7] measurements were made in each limitation series, while crosses represent the growth rates of the samples for which RNA-seq measurements were made.

**(B)** The quantification of the fold-change (FC) in protein and mRNA across the growth range $\lambda_1 = 0.35/\text{h}$ to $\lambda_2 = 0.91/\text{h}$ in each one of the three growth limitation series is illustrated using the example of the *fumA* gene in carbon-limited growth. The log-transformed protein (red circles) and mRNA (red crosses) fractions were fitted with quadratic functions $\psi_{\mathrm{m},i}^{\mathrm{fit}}(\lambda)$ and $\psi_{\mathrm{p},i}^{\mathrm{fit}}(\lambda)$ respectively (blue lines). The fit allowed precise estimate of the (log-transformed) mRNA and protein abundances at the growth rates $\lambda_1$ and $\lambda_2$, together with their standard error. Errors on $\log_{10} FC(\psi_{\mathrm{p},i}) = \log_{10}\left(\psi_{\mathrm{p},i}^{\mathrm{fit}}(\lambda_2)/\psi_{\mathrm{p},i}^{\mathrm{fit}}(\lambda_1)\right)$, $\log_{10} FC(\psi_{\mathrm{m},i}) = \log_{10}\left(\psi_{\mathrm{m},i}^{\mathrm{fit}}(\lambda_2)/\psi_{\mathrm{m},i}^{\mathrm{fit}}(\lambda_1)\right)$ and $\log_{10}(FC(\psi_{\mathrm{p},i})/FC(\psi_{\mathrm{m},i}))$ are then propagated from the errors on the fitted abundances. The genes for which the corresponding proteins and mRNAs vary by similar extents across growth rates of $\lambda_1$ to $\lambda_2$ in a given growth limitation exhibit values of $FC(\psi_{\mathrm{p},i})/FC(\psi_{\mathrm{m},i}) \sim 1$ for that limitation.

**(C, D)** Comparison of $FC(\psi_{\mathrm{p},i})/FC(\psi_{\mathrm{m},i})$ obtained for $N = 1452$ genes from the three different limitations: (C) translation limitation vs. carbon limitation, (D) translation limitation vs. anabolic limitation. Colored points indicate outlier genes for which the ratio $FC(\psi_{\mathrm{p},i})/FC(\psi_{\mathrm{m},i}) > 3$ in at least one of the three growth limitations, for levels of significance $p \leq 1\%$ (red) and $1\% < p \leq 10\%$ (orange) (two-sided Z-test with Bonferroni correction for triple comparison); the rest of the points are non-significant ($p > 10\%$).

**(E)** The mRNA (crosses) and protein (circle) fractions are plotted against growth rates for each of the three growth limitations for several of the outlier genes in panels (C) and (D), to show the discordance between mRNA and protein levels in at least one of the growth limitations. A complete list of genes exhibiting different growth rate dependencies for mRNA and protein fractions is provided in Table S3, along with potential mechanisms of regulation that could contribute to the discordance between mRNA and protein abundances.

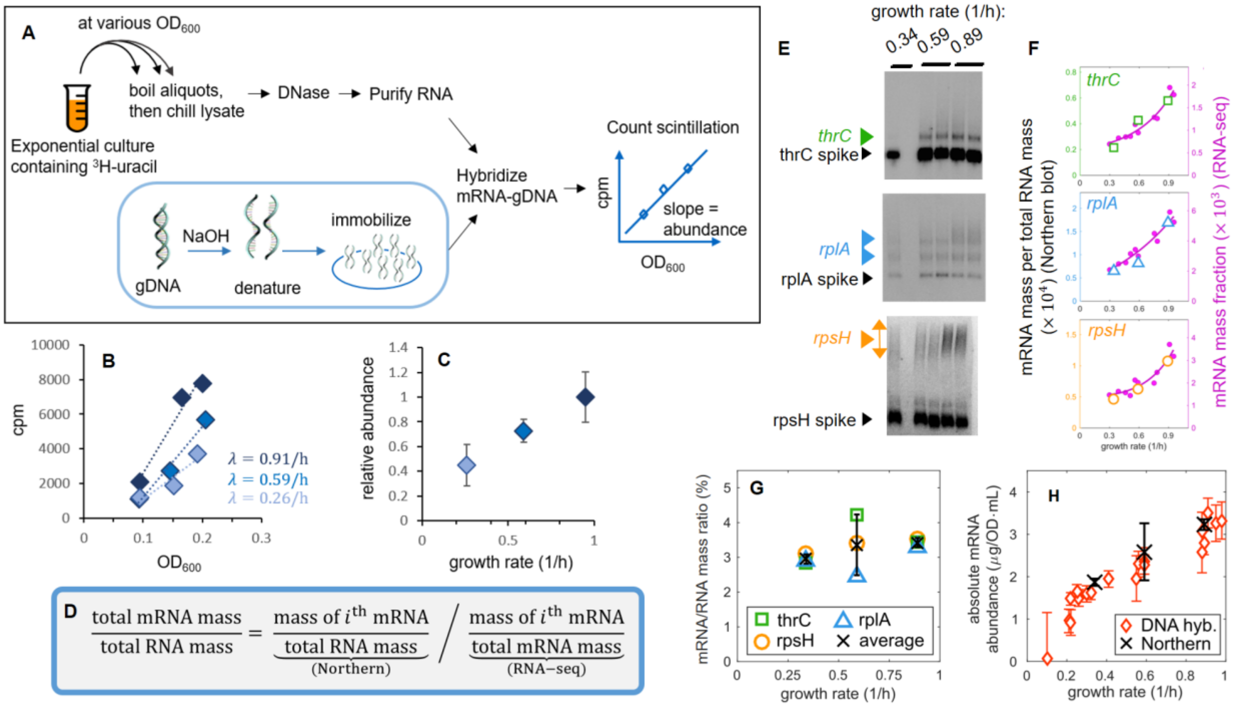


**Figure S5. Measurement of total mRNA mass abundance.**

**(A)** Schematic for the measurement of relative total mRNA mass abundance as described in Ref. [14]. Cultures are continuously labelled with $^3$H-uracil and sampled at different values of culture density ($OD_{600}$). The lysates are treated with DNase and RNA is purified in order to remove un-incorporated nucleotides. The remaining labelled RNA is hybridized to denatured genomic DNA (gDNA) [15] that is immobilized on membrane filters (SI Methods). The label incorporated into the hybridizable RNA (assumed to be predominantly mRNA) is measured by scintillation counting; the slope of counts versus OD gives the estimate of relative total mRNA mass abundance.

**(B)** Example of the relative total mRNA abundance measured for cultures growing on three different carbon sources: glucose (growth rate $\lambda = 0.91/\mathrm{h}$), succinate ($\lambda = 0.59/\mathrm{h}$) and mannose ($\lambda = 0.26/\mathrm{h}$); see Table S1 for growth conditions. Counts per minute are plotted versus the absorbance of the cultures ($OD_{600}$); The slopes yield the relative total mRNA mass abundance per culture volume (OD·mL).

**(C)** The slopes from panel B relative to that of the reference condition (NCM3722 in MOPS glucose, $\lambda$ = 0.91/h) are plotted against the growth rate of each culture. Error bars indicate the uncertainty in fitting of the slopes from panel B.

**(D)** Strategy to obtain the absolute mass fraction of mRNA per total RNA by normalizing quantitative Northern Blotting with RNA-sequencing data. Probing the blots for the $i^{\mathrm{th}}$ endogenous mRNA yields the abundance of that mRNA per total RNA. Normalizing this quantity with the mass fraction of the $i^{\mathrm{th}}$ mRNA per total mRNA (as obtained by RNA-seq data; see Supplementary Note S1 for conversion between number and mass fractions) allows us to compute the proportion of total RNA that is mRNA. Importantly, this quantity can be determined using different specific mRNAs as probes as cross-checks.

**(E)** Examples of quantitative Northern blots. RNA samples extracted from three different carbon-limited cultures (growth rates indicated above the respective lanes) were mixed with known amounts of *in vitro*-transcribed spike-in RNA fragments and resolved on formaldehyde

agarose gels. The gels were blotted onto nylon membranes and probed with radiolabeled oligos specific for either *thrC* mRNA (top), *rplA* mRNA (middle) or *rpsH* mRNA (bottom). The hybridization of the probes to endogenous mRNA as well as the spike-in sequences allows for the estimation of the amount of each of these target mRNAs per μg of total RNA *(*SI Methods).
**(F)** Abundance of the target mRNAs (*thrC, rplA* and *rpsH*) obtained as mass fraction of the total RNA by Northern Blotting (open symbols, left vertical axis) and as mass fraction of total mRNA by RNA-sequencing (purple dots, right vertical axis) at different growth rates are shown for comparison. Conversion between mRNA number and mass fractions is described in detail in SI Note S1.
**(G)** The ratios of the total mRNA mass to total RNA mass, computed as described in (D) by combining the measurements shown in (F, left and right axes) for each of the probes as indicated by the colored symbols, are plotted against the growth rate of the culture. The average ratio (black crosses; error bars represent standard deviations), ranges between 3 and 3.5% with a possible mild growth rate dependence, in agreement with previous measurements [15, 16].
**(H)** The absolute mRNA abundance, in units of μg/OD/ml, is computed using previously measured total RNA content (in total RNA mass per culture volume) across growth rates [6], and the average ratio of mRNA to total RNA from Northern Blotting (black crosses). The absolute mRNA abundance in the reference condition (growth rate 0.91/h) is then used to rescale the relative total mRNA abundances measured by the DNA hybridization method (panels A-C) for the range of carbon limitation conditions to obtain the absolute mRNA mass per culture volume for these growth conditions (red points). The data was finally converted to the units shown in Fig. 2A using the total cell volume per culture volume, Fig. S2C, and the average mRNA length (see Supp. Note S1 for details).

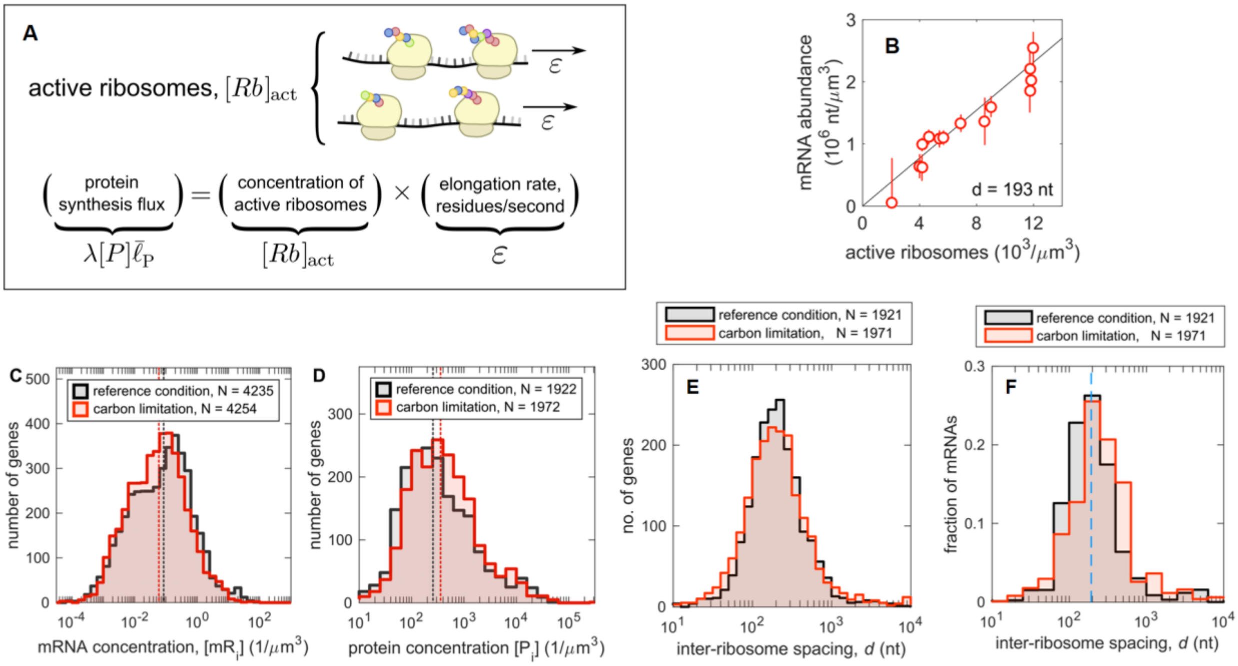


**Figure S6. Conversion of quantitative proteomics and transcriptomics data into concentrations.**

**(A)** The total flux of synthesized proteins, in units of residues per cell volume per time, is obtained as the total protein flux $\lambda[P]$ times the average protein length $\bar{\ell}_{\mathrm{P}}$. The same quantity can be expressed the product of the concentration of active ribosomes in the culture $[Rb]_{\mathrm{act}}$ and the translation elongation rate $\varepsilon$, yielding Eq. (5) in the Main Text. Both the elongation rate and the abundance of active ribosomes have been characterized [17] for a broad range of carbon-limited growth; see SI Note S2 for details.

**(B)** The total abundance of mRNA in terms of number of nucleotides per cell volume is obtained from the measured mRNA mass per culture unit (Figure S5H), the total cellular volume (Fig. S2C) and the average nucleotide mass $m_{\mathrm{nt}} = 5.38 \cdot 10^{-16}$ μg. The result is plotted against the concentration of active ribosomes, interpolated to the growth rates of the mRNA measurements. The black solid line represents the best fitting line passing through the origin; its slope provides the average inter-ribosome spacing, $d \approx 193$ nt.

**(C)** Distribution of mRNA copy number per cell in reference condition (black) and carbon-limiting condition (red; growth rate $0.33/\mathrm{h}$). Absolute concentrations were computed using the measured total mRNA mass abundances (Fig. S5H) and the total cellular volume per culture unit (Fig. S2C) (see Note S1 for details). The median mRNA levels (dashed vertical lines of the corresponding colors) are low, 0.085 copies per cell in reference condition and 0.06 per μm$^3$ at slow growth.

**(D)** Same as panel C, but for protein concentrations in the two growth conditions. Conversion to absolute units was done assuming $[P] = 3.15 \cdot 10^6/\mu\mathrm{m}^3$ across all conditions (see Figure S2F). Mass spectroscopy data [7] covers more than 1900 proteins with concentrations spanning between 100 and $10^5$ copies per μm$^3$; The median protein levels are 248 and 344 copies per cell in reference condition or slow growth, respectively (dashed lines).

**(E)** Distribution of inter-ribosome spacing, $d_i = \left(\psi_{\mathrm{m},i}/\psi_{\mathrm{p},i}\right) \cdot \bar{d}$, computed across >1900 genes. The distributions are centered around $d \approx 200$ nt. Fewer than 4% of genes have spacings $<$

$40$ nt, the nominal distance if ribosomes are “close-packed” on the mRNAs [18], indicating the self-consistence of the data. Note also the lack of genes (< 5.6%) with less than 1 ribosome per kb (the average gene length).
**(F)** Same as panel (E), but weighing each gene by mRNA abundance. The dashed line indicates the average inter-ribosome spacing, $d = 193$ nt.

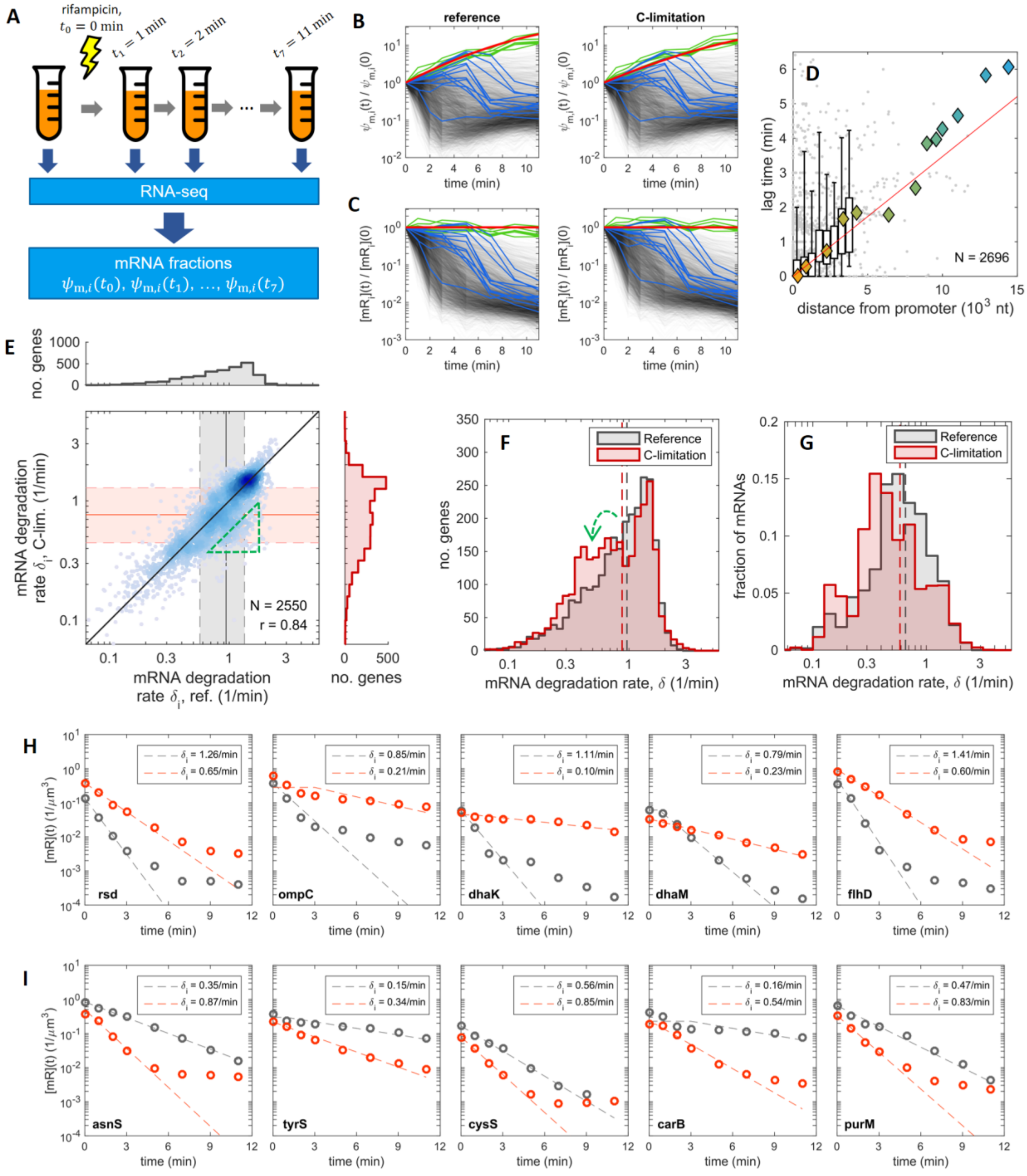


**Figure S7. mRNA degradation rates.**
**(A)** Sketch representing the experimental procedure for measurement of mRNA degradation rates. Transcription initiation was stopped (at $t = 0$) by applying rifampicin to exponentially growing culture, and RNA-sequencing was performed for samples collected at $t = 0$, 1, 2, 3, 5, 7, 9 and 11 minutes, allowing us to obtain the mRNA number fractions $\psi_{\mathrm{m},i}(t)$ at each time point for >3500 genes in reference condition (growth rate = 0.95/h) and carbon limiting condition (growth rate = 0.34/h) (SI methods; Table S2).

**(B)** Grey lines indicate fractional abundances of mRNA relative to the pre-rifampicin condition, $\psi_{\mathrm{m},i}(t)/\psi_{\mathrm{m},i}(0)$ for reference condition (left plot) and slow growth (right plot). In order to convert this quantity to the relative mRNA abundance $[\mathrm{mR}_i(t)]/[\mathrm{mR}_i(0)]$, the data has to be corrected for the reduction over time of the total mRNA abundance, $[\mathrm{mR}(t)]/[\mathrm{mR}(0)]$, which modulates the mRNA fractions globally. Such modulation (red curve) is estimated using the time dependence of the most stable mRNAs (top 0.2% of genes shown in green) and the abundance of mRNA in polycistronic operons far from the promoters (more than 10 kbp, shown in blue); see SI Methods for details.
**(C)** Relative mRNA concentrations, $[\mathrm{mR}_i(t)]/[\mathrm{mR}_i(0)]$, obtained after removing the global modulation observed in the previous panel (red horizontal line). These are then fitted with a lagged exponential decay, using the mRNA degradation rate and a lag time as fitting parameters (see SI Methods). After filtering out low-quality fits, we estimated mRNA degradation rates for 2696 genes in reference condition and 2883 genes in carbon limiting condition.
**(D)** The lag-time between rifampicin addition and the start of mRNA degradation obtained by fitting the data in panel (C) with a lagged exponential is plotted for 2696 genes against their distance from their promoters in the reference condition. Data for genes within 4 kbp from their promoters is binned to aid visualization; boxes and whiskers indicate 50% and 90% of the lag times in each bin, respectively. Overall, the RNAP elongation rates agree with the *in vivo* measured transcription elongation rate of 48/min [19] which is represented by the red line. Diamond symbols indicate the genes of the *nuo* operon, with the same color scheme as in Fig 3A.
**(E)** Genome-wide mRNA degradation rates are plotted for the reference condition (horizontal axis) and the carbon limited condition (vertical axis); the two are strongly correlated (Pearson coefficient $r = 0.84$ between the log-transformed quantities). The distributions of the degradation rates for each condition are shown beside the respective axes, with half of the genes (25$^{\mathrm{th}}$ to 75$^{\mathrm{th}}$ percentiles, colored bands) between approximately 0.5/min and 1.3/min in each condition, and the median value (dashed lines) around 0.8/min for both conditions. The triangular region enclosed by the green dashed lines contains a number of mRNAs whose stability is increased at slow growth compared to reference condition.
**(F)** Distribution of mRNA degradation rates (same as those plotted on the side bars of panel D). Vertical dashed lines indicate the average mRNA degradation rates across the genome, $\sum_i \delta_i/N$, which equals 0.964/min in reference condition (black, $N = 2696$) and 0.879/min in carbon-limited growth (red, $N = 2883$). The subset of genes being stabilized at slow growth (triangle enclosed by green dashed line in panel D) is visible as a hump in the distribution of degradation rates in carbon-limited growth (red), see green dashed arrow.
**(G)** Distribution of mRNA degradation rates, weighted by mRNA abundance. The dashed lines indicate the average mRNA degradation rate, $\bar{\delta} = \sum_i \delta_i \psi_{\mathrm{m},i}$, equal to 0.655/min in reference condition (growth rate 0.96/h) and to 0.586/min in carbon-limited growth (growth rate 0.34/h).

**(H-I)** Examples of genes that show different mRNA degradation dynamics for cells grown in reference (black) and carbon limited (red) conditions, either more stable in carbon limted growth (H) or in reference condition (I). Circles represent mRNA concentrations over a duration of 11 minutes following blockage of transcriptional initiation (see Methods; dashed lines represent the fitted model (lagged exponential decay) with the best-fit decay rate ($\delta_i$) shown in the legend. Many of these genes are known to be regulated by small RNAs which could explain the difference in stabilities. For example, *purM* is regulated by CsrA [20]; *ompC* by MicC [21];

*dhaM* and *dhaK* by RyhB [22] and *flhD* by ArcZ, OmrA, OmrB, OxyS [23] and McaS [24]. While no specific post transcriptional regulation of *rsd* has been reported, our results show an increased stability of its mRNA at slow growth which could result in its higher expression that is discussed in Fig. 6.

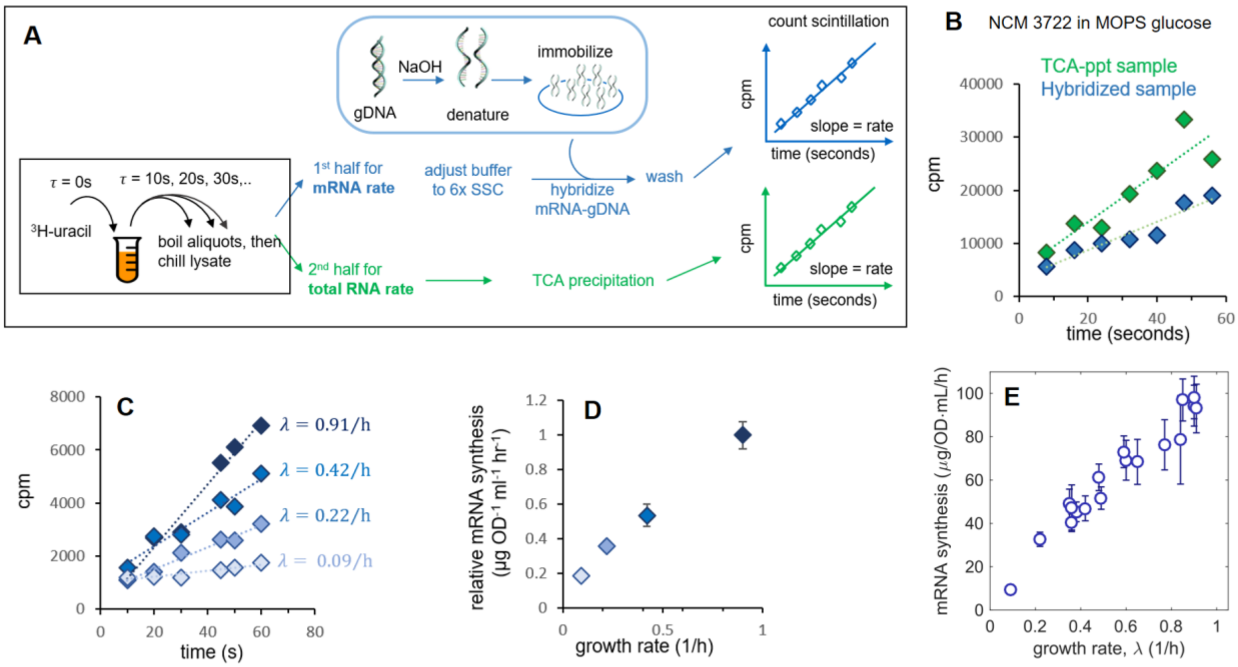


**Figure S8. Measurement of the absolute rates of mRNA synthesis.**

**(A)** Schematic illustration of procedure. Exponentially growing cultures are pulse-labelled with $^{3}$H-uracil and aliquots are sampled every 10 seconds over the ensuing minute. The samples are lysed and split into two. In one half (blue), the RNA is hybridized to *E. coli* genomic DNA that is denatured and immobilized on cellulose filters. The filters are then washed and the radioactivity of the hybridized RNA is measured by scintillation counting. The slope of scintillation counts versus time gives the relative rate of mRNA synthesis. In the second portion of the lysate (green), total RNA is precipitated using trichloroacetic acid (TCA) and washed. Radiolabel incorporation into the total RNA is followed by scintillation. The relative mRNA synthesis rates were determined for cells growing in various growth conditions using the hybridization method (blue). Next, these measurements were mapped to an absolute scale by determining the absolute mRNA synthesis rate at the reference growth condition (glucose minimal medium), as described in (B).

**(B)** $^{3}$H-uracil incorporation kinetics into the TCA-precipitated total RNA fraction (green) and DNA-hybridized mRNA fraction (blue) are shown for cells growing in reference condition. Dashed lines represent linear fits to the data. By comparing the slopes of the $^{3}$H-uracil incorporation kinetics in the total RNA and mRNA (corrected for 75% hybridization efficiency) we estimate that mRNA synthesis flux to be 52% of the total RNA synthesis flux for growth in the reference condition, with the remainder being stable RNA. This approximate equal synthesis of mRNA and stable RNA for such growth conditions is consistent with earlier reports [15, 16, 25, 26]. The value of the synthesis rate of stable RNA can be determined as the product of the total RNA abundance and the growth rate, measured in Ref. [6] for a range of carbon-limiting conditions. Assuming the stable RNA synthesis flux of $93.7\ \mu g/OD_{600}/mL/h$ in our condition, the value for mRNA synthesis rate can be determined to be $101.5\ \mu g/OD_{600}/mL/h$ in the reference growth condition.

**(C)** $^{3}$H-uracil incorporation kinetics into mRNA fractions (hybridization experiment) is shown for growth in different conditions including glucose (0.91/h), acetate (0.42/h), aspartate +$NH_4Cl$ (0.22/h) and glutamate+$NH_4Cl$ (0.09/h), see table S1 for growth conditions.

**(D)** The slopes of $^3$H-uracil incorporation over time obtained from panel (C) are normalized to that of the reference condition (0.91/h), and plotted against the respective growth rates. Uncertainty in the fits of the slopes is indicated by error bars.
**(E)** Absolute rate of mRNA synthesis per culture volume per hour, measured across growth rates in carbon-limited conditions (see Table S1 for a list of growth conditions). Relative data were calibrated using the absolute mRNA synthesis rate in reference condition, panel (B). Data was then converted to cellular units by dividing by the total cellular volume per culture volume (Fig. S2C) and by the average mRNA mass across conditions (SI Note S1) to yield the data shown in Fig. 3FG.

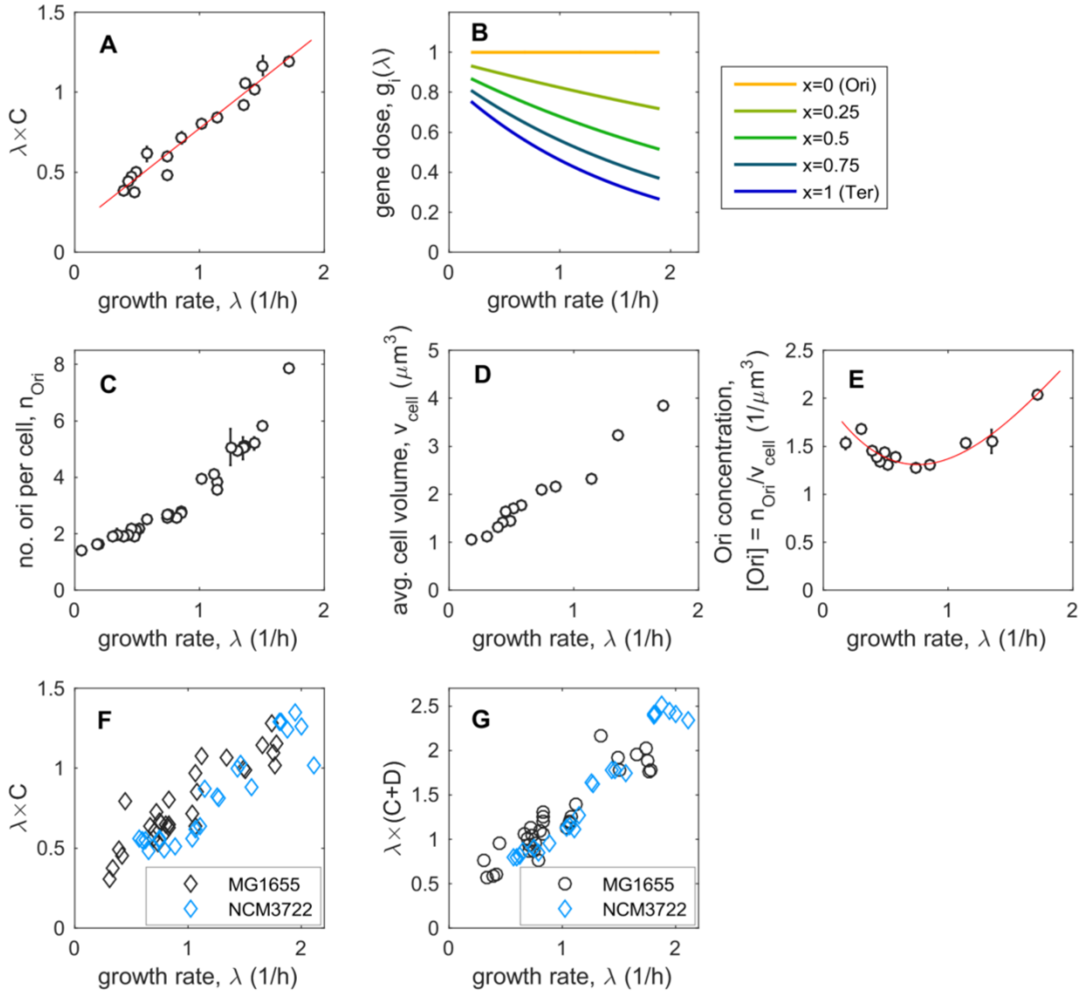


**Figure S9. Growth rate dependence of gene concentrations.**
**(A)** The product of the growth rate $\lambda$ and the C-period ($C$) obtained for *E. coli* MG1655 [27] is well described by a linear function of the growth rate (red line).
**(B)** The gene dose $g_i = [G_i]/[Ori]$ can be expressed in terms of $\lambda C$ and the distance $x$ of the gene from Ori as $g_i = \exp(\lambda C x_i)$ [28]. Here we show the estimated $g_i$ as a function of growth rate for genes at various distances from Ori using the linear fit for $\lambda C$ shown in panel (A).
**(C)** Average number of Ori per cell $n_{\text{Ori}}$, from Ref. [27].
**(D)** Average cell volume $v_{\text{cell}}$, from Ref. [27].
**(E)** By combining the average number of Ori per cell and the average cell volume (panels C and D), we computed the Ori concentration $[Ori] = n_{\text{Ori}}/v_{\text{cell}}$. The red line represents a fit with a third-degree polynomial, which can then be combined with the gene density (panel B) to yield the protein concentrations $[G_i]$ across all growth conditions, as shown in Fig. 4A.
**(F)** Measurement of $\lambda C$ from Ref. [29] for *E. coli* strains MG1655 and NCM3722. The data shows that the C-period is similar for both strains.
**(G)** Measurement of $\lambda(C + D)$ from Ref. [29] for *E. coli* strains MG1655 and NCM3722. The quantity $C + D$ represents the cell cycle duration, and sets the number of Ori per cell, $n_{Ori} =$

$\exp(\lambda(C + D))$ [28, 29]. The data is similar for both strains. Together, the data here and in panel (F) suggest that the DNA replication dynamics is similar between these two strains.

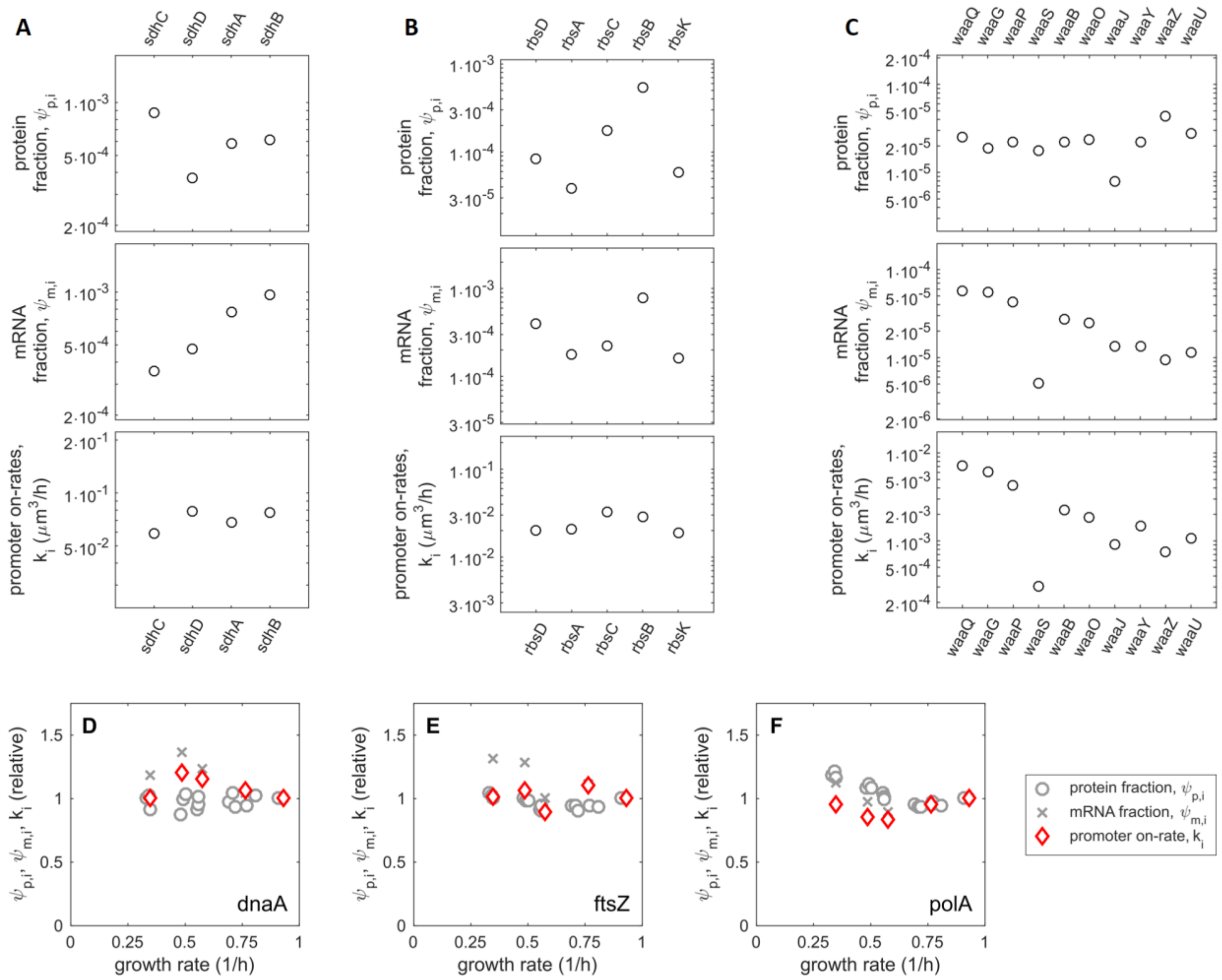


**Figure S10. Promoter on-rates within operons.**

Protein fractions $\psi_{\mathrm{p},i}$ (top), mRNA fractions $\psi_{\mathrm{m},i}$ (middle) and promoter on-rates $k_i$ (bottom) for three exemplary operons: *sdh* (A, left), *rbs* (B, center) and *waa* (C, right). Genes are sorted so that the distance from the common promoters increases from left to right. *sdhD* and *rbsD* mRNA are known targets of small RNA regulators which could potentially explain the discrepancy between the promoter on-rates, mRNA and protein abundances; see text.

**(D-F)** The sum of the promoter on-rates for subsets of genes are plotted as red diamonds as a function of growth rate. For comparison, we also show the total protein fraction (grey circles) and total mRNA fractions (grey crosses) for these genes; crosses are averages between two replicates. Examples of genes exhibiting approximately constant promoter on-rate across the growth conditions examined.

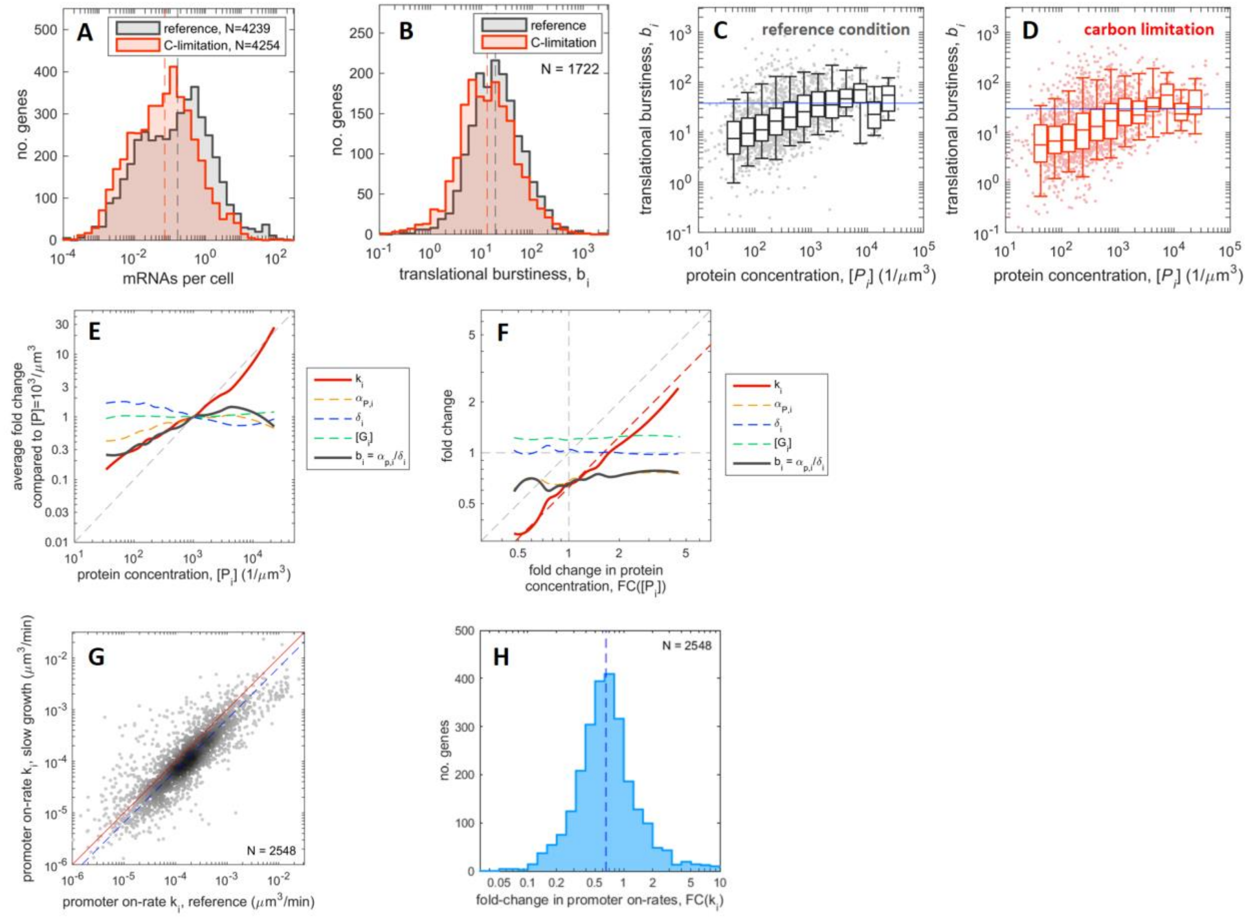


**Figure S11. Translational burstiness and gene expression parameters.**

**(A)** Copy number of mRNAs per cell in reference condition (grey) and slow growth (red), computed by dividing the mRNA concentrations (Fig. S6C) by the average cell volume at the corresponding growth rate, Fig. S2B. The lower abundance of mRNAs at slow growth is visible as a shift in the distributions. Dashed lines indicate the median values (0.165/cell in reference condition, 0.072/cell at slow growth).

**(B)** The translational burstiness parameter $b_i$ is defined for each gene $i$ as the ratio of translational initiation rate $\alpha_{\mathrm{p},i}$ and the mRNA degradation rate, i.e., $b_i = \alpha_{\mathrm{p},i}/\delta_i$. It represents the average number of proteins synthesized from a single mRNA during its lifetime. For both reference condition and slow growth, the distribution of burstiness parameters is concentrated around ~10; the median burstiness parameters in the two conditions are indicated as dashed vertical lines (18.9 in reference condition, 13.1 at slow growth).

**(C)** Scatter plot of the burstiness parameter $b_i$ versus protein concentrations $[P_i]$ in reference condition. Boxes and whiskers indicate the 50% and 90% central intervals of the binned data; the central line indicates the median burstiness in each bin. The solid blue line indicates the median burstiness parameter (38.4) for protein concentrations larger than $10^3/\mu\mathrm{m}^3$. The data shows a gradual reduction in burstiness for low-expressed proteins, i.e., those with $[P_i] < 10^3/\mu\mathrm{m}^3$.

**(D)** Same as panel (C), but for slow growth in carbon-limited conditions. The median translational burstiness for protein concentrations larger than $10^3/\mu\mathrm{m}^3$ is 29.7.

**(E)** The moving averages shown in Figure 5C for the promoter on-rates $k_i$, translation initiation rates $\alpha_{\mathrm{p},i}$, mRNA degradation rates $\delta_i$ and gene concentrations $[G_i]$ are rescaled by their values

at $[P_i] = 10^3/\mu\text{m}^3$ and re-plotted here. Additionally, the average burstiness is obtained as the ratio of the (rescaled) moving averages of $\alpha_{\text{p},i}$ and $\delta_i$ (solid black line). The expression of proteins with concentrations above $10^3/\mu\text{m}^3$ can be accounted for primarily by setting the promoter on-rate (red solid line): higher expression levels are associated with proportionally higher promoter on-rates (compared the red line to the dashed grey, with slope 1). Instead, expression levels lower than $10^3/\mu\text{m}^3$ are accounted for by using a combination of setting in transcriptional regulation ($k_i$) and translational burstiness ($b_i$), as shown by the overlapping red and black lines.
**(F)** The moving averages shown in Figure 5D for the fold changes in rates and concentrations across conditions (slow growth compared to reference condition) are shown here on the same plot. In this case, the fold change in protein concentration is proportional to the fold change in promoter on-rate (red) across the whole range (slope $\approx 1$), indicating that changes in protein levels are primarily set at the transcription level. The global shift in the promoter on-rates, Fig. 5A, is visible as a gap between $FC(k_i)$ and the diagonal (grey dashed line); the dashed red line indicates the median reduction of 37% of $FC(k_i)$ compared to $FC([P_i])$.
**(G)** Scatter plot comparing promoter on-rates for >2500 genes between the reference growth condition and carbon limited growth. The red line represents the diagonal (x=y). The dashed blue line represents the median ratio 0.65 of the promoter on-rates, i.e. an overall 35% reduction of promoter on-rates at slow growth compared to reference condition.
**(H):** Histogram representing the fold-change in promoter on-rates in carbon-limited growth compared to that in the reference condition. The dashed line indicates the median value 0.65 (same as in panel G).

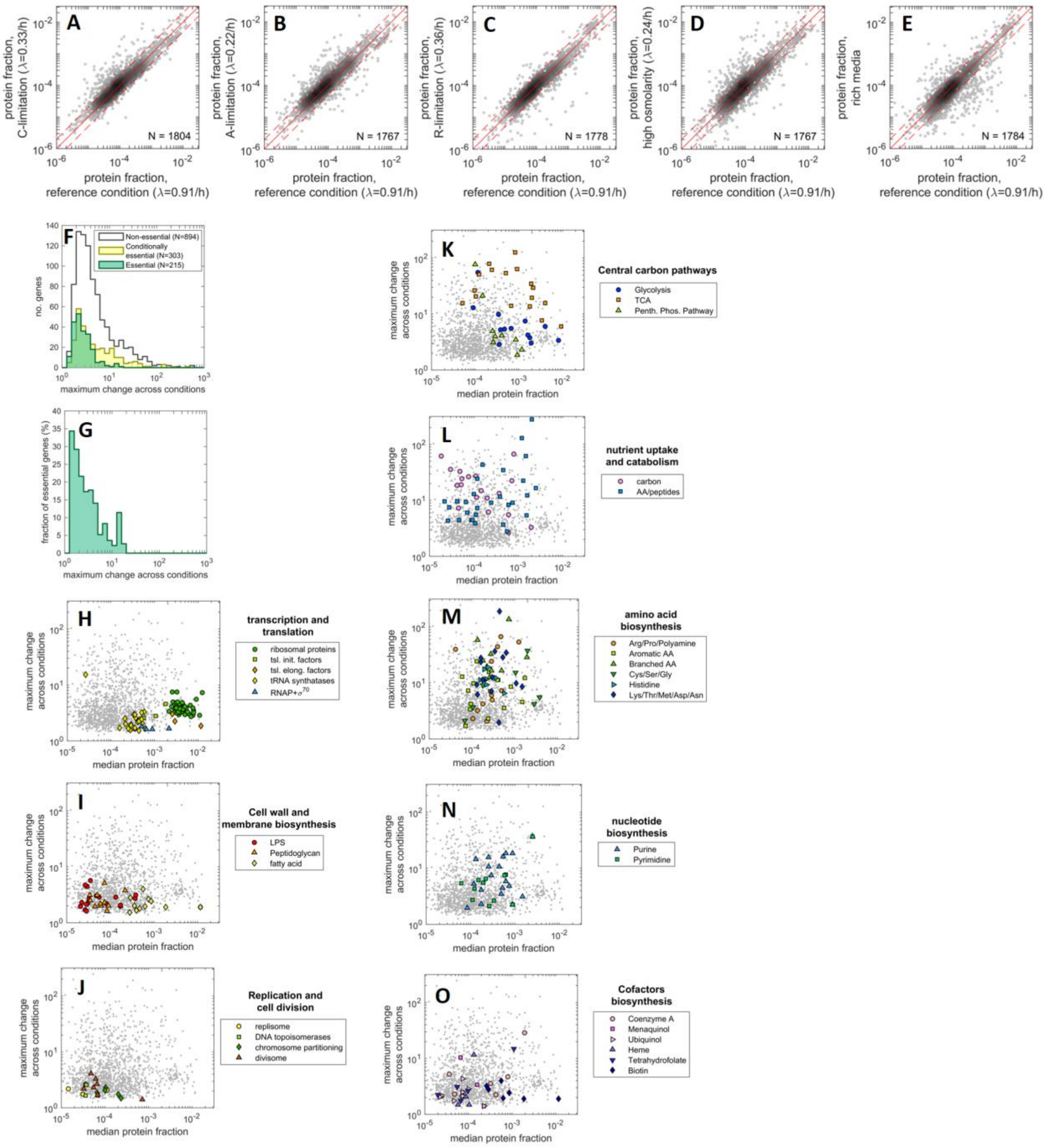


**Figure S12. Growth condition-dependent changes in protein abundances.**
**(A-E)** Comparison of protein abundances between the reference growth condition and other conditions including carbon limitation, anabolic limitation, ribosome limitation, high osmolarity or rich-media. The data is obtained from the proteomic library recently published [7]. The solid red lines represent the diagonal while the dashed red lines represent 2-fold changes across conditions.
**(F)** The distribution shows the maximum difference in the protein fraction abundances between any 2 growth conditions among a set of 19 different growth conditions [7]. The 1412 proteins detected in all 19 samples were classified as either non-essential (894 proteins), conditionally

essential (i.e. essential in certain conditions, 303 proteins), and essential (215 proteins), using the annotation available on Ecocyc [30]. While many non-essential and conditionally-essential proteins show changes over 5-fold across conditions, proteins encoded by essential genes are largely excluded from this region.
**(G)** For each bin in panel F, we show the fraction of proteins with given maximum change that is an essential protein. For maximum fold-changes above 5-fold, a small fraction (~10% or less) of the genes are annotated to be essential.
**(H-J)** Same as Fig 5E, but highlighting different groups of genes displaying little variability in their protein abundances across the 19 conditions tested [7]. Ribosomal proteins (green, panel H) show ~4-fold maximum change in their abundance across conditions. Other translation associated proteins such as initiation factors and tRNA synthetases are less abundant and show even smaller fold-changes across growth conditions, highlighting differences in their regulatory mechanisms. Proteins belonging to the biosynthesis of cell-wall components (I) such as lipopolysaccharides (LPS), peptidoglycan and fatty acids occupy different ranges in protein abundances but show little variability across conditions. The low abundant proteins involved in DNA replication and cell division are one of the groups showing the least variabilities across conditions.
**(K-O)** In contrast to panels H-J, in these panels we show examples of proteins showing large fold-change differences across growth conditions. These are enzymes of the central carbon metabolism pathways (K), transporters involved in carbon and peptide uptake and catabolism (L), or enzymes involved in amino acid (M), nucleotide (N) and cofactor (O) biosynthesis. Most of these pathways are known to be associated with highly specialized regulatory mechanisms. Another example is provided by motility-associated genes, whose expression changes by orders of magnitude across conditions [7]; however, the corresponding proteins are not shown here as their minimum protein level is below detection threshold.

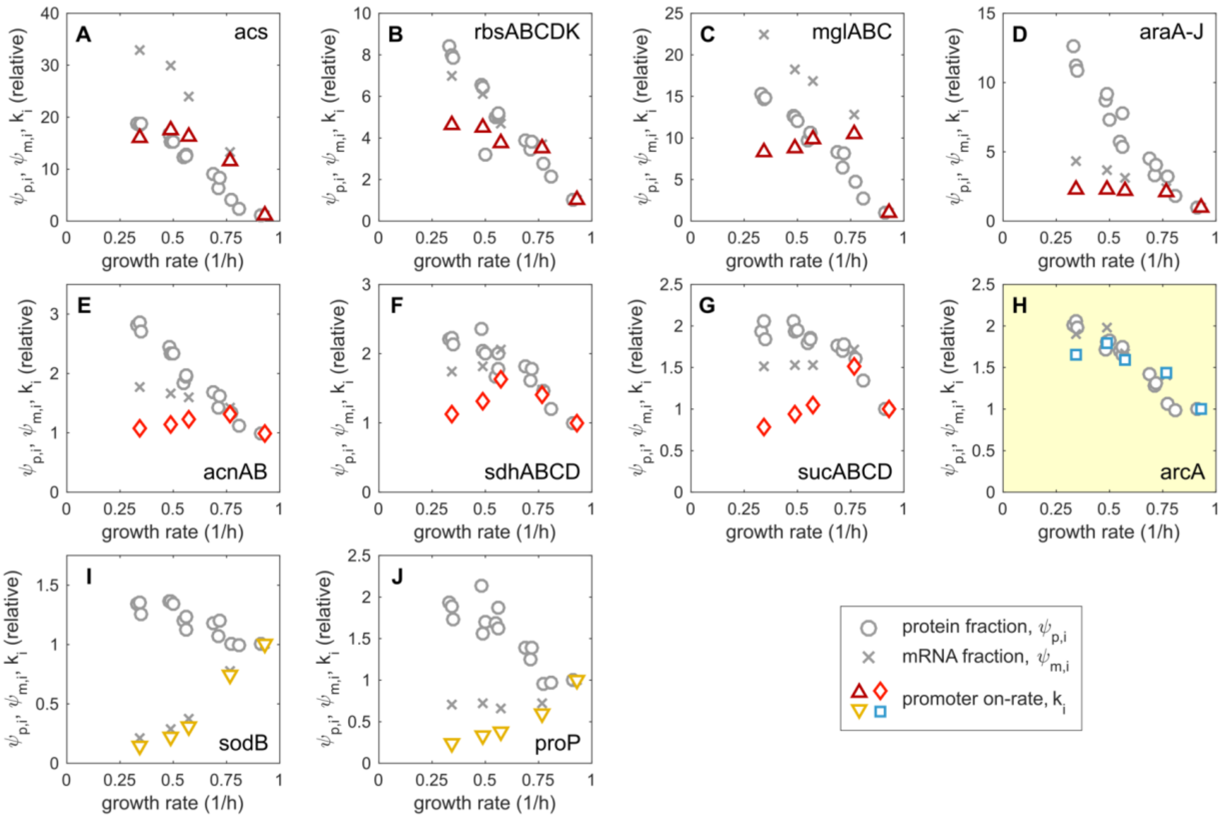


**Figure S13. Promoter on-rates for proteins upregulated in carbon limitation.**
Same as in Fig. S10D-F, but for proteins upregulated in carbon limitation. Promoter on-rates are shown with colored symbols (matching those in Fig. 5F, except for ArcA).
**(A-D)** The promoter on-rates for carbon-uptake proteins increase rapidly and then stabilized as growth rate is lowered from that of reference condition due to carbon limitation. These responses are consistent with the activation of these genes by cAMP-Crp, whose activity increases under carbon limitation due to increased cAMP level [31–33]. Note that these operons are selected out of the many tens of Crp-regulated genes because these outputs are much larger than the rest, presumably because most Crp-regulated genes are inhibited by their sugar-specific regulators [7]. In the low-growth regime, the lack of increase in the promoter on-rate may be due to saturation of cAMP-Crp activity: the activation of Crp-dependent promoters saturates above a few μM of cAMP [34], which is the range of cAMP concentrations in vivo in poor carbon conditions [35]. Further increase in the protein and mRNA fractions at slow growth is attributed to reduction in the total regulatory activity $\mathcal{K}$ according to Eq. (12) of the Main Text.
**(E-G)** Promoter on-rates for TCA cycle enzymes exhibit non-monotonic growth rate dependence, with maximal expression close to growth rate $\sim 0.7/h$.
**(H)** The high levels of transcription factor ArcA at slow growth might be the cause of the reduced promoter on-rates for the TCA genes (panels E-G), which are activated by cAMP-Crp and repressed by (phosphorylated) ArcA [36, 37].
**(I-J)** Two other genes, *sodB* and *proP*, whose protein concentrations increase in carbon-limitation, but the promoter on-rates decrease. These cases may reflect instances of post-transcriptional regulation.

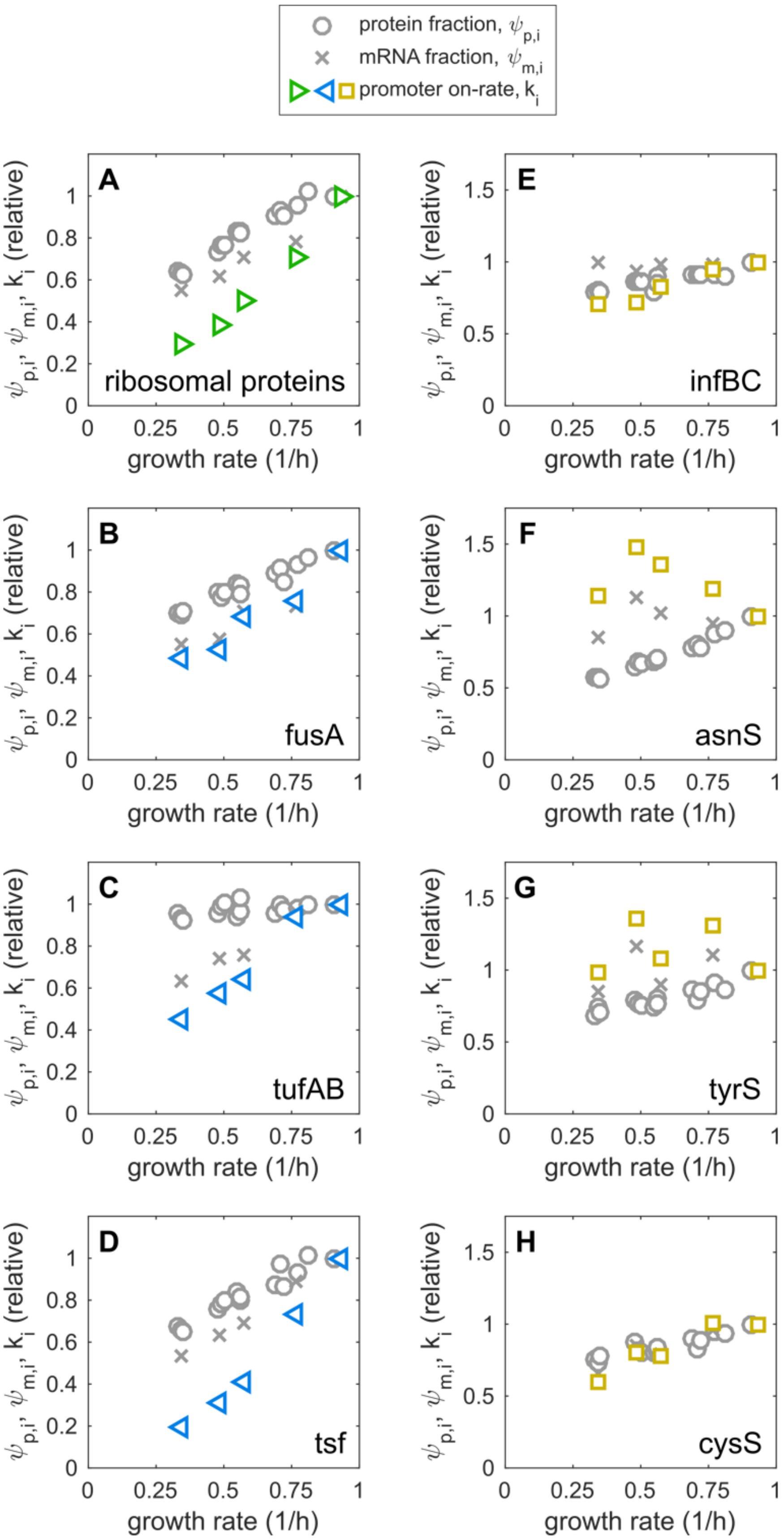
protein fraction, $\psi_{p,i}$
mRNA fraction, $\psi_{m,i}$
promoter on-rate, $k_i$
A
ribosomal proteins
B
fusA
C
tufAB
D
tsf
E
infBC
F
asnS
G
tyrS
H
cysS
$\psi_{p,i}$, $\psi_{m,i}$, $k_i$ (relative)
growth rate (1/h)

**Figure S14. Promoter on-rates for ribosomal proteins.**
Same as in Fig. S10D-F, but for groups of ribosomal and ribosome-associated proteins. Promoter on-rates are shown with colored symbols.
**(A)** The sum of the 54 ribosomal proteins and **(B-D)** translational elongation factors (symbols matching those in Fig. 5G).
**(E-H)** Other translation-associated proteins for which the growth rate dependence of the promoter-on rates is markedly different from those observed for ribosomal proteins and rRNA.

### SUPPLEMENTAL TABLES

**Table S1a** Strains used in this study

**Table S1b** Growth conditions used in this study.

**Table S2.** Transcriptomics and proteomics data used in this work. Sheet 1: Description of each proteomics sample. Sheet 2: mass spectrometry data, in units of protein fractions $\psi_{\mathrm{m},i}$. Sheet 3: Description of each RNA-seq sample for steady state mRNA abundances. Sheet 4: RNA-seq data, in units of mRNA fractions $\psi_{\mathrm{m},i}$. Sheet 5: description of each RNA-seq sample for the determination of mRNA degradation rates. Sheet 6: RNA-seq data for the determination of mRNA degradation rates, in units of mRNA fractions $\psi_{\mathrm{m},i}$.

**Table S3.** Quantification of growth-dependent transcription initiation. Sheet 1: Fold change in relative translation efficiency ($\psi_{\mathrm{p},i}/\psi_{\mathrm{m},i}$) for all genes in each of the three growth limitations (see Figure 1E and Figure S4). Sheet 2: List of genes with the largest variation in the relative translation efficiency.

**Table S4.** Summary of gene expression parameters for each gene, including protein and mRNA concentrations, mRNA degradation rate, translational burstiness, transcription and translation initiation rates, gene concentration and promoter-on rate. Sheet 1: reference condition ($\lambda \sim 0.9/\mathrm{h}$). Sheet 2: carbon-limited slow growth ($\lambda \sim 0.3/\mathrm{h}$).

**Table S5.** Proteomics data (from Ref. [7]) associated to Figure 5E and Figure S13. Sheet 1: the detailed list of the samples used in the analysis. Sheet 2: protein mass fractions, median and maximum fold change across all conditions for the set of 1415 proteins detected across all conditions.

Tables S2-5 are not included in this biorxiv submission. These tables are available from the corresponding author (T.H.) upon reasonable request.

| strain name | Description | purpose | reference |
|---|---|---|---|
| NCM3722 | Wild-type *E. coli* K12 | reference and parental strain | Soupene et al 2003 |
| NQ1243 | NCM3722, Pu-ptsG | titration of glucose uptake | Basan et al 2015 |
| NQ1390 | NCM3722, Pu-ptsG | titration of glucose uptake | Basan et al 2015 |
| NQ393 | NCM3722, ΔgdhA, $P_{Llac-01}$-gltBD | titration of Nitrogen assimilation (A-limitation) | Hui et al 2015 |
| HE725 | NCM3722, Δrsd:cat | rsd deletion strain | this work |
| HE805 | NQ1390, Δrsd:cat | titration of glucose uptake in Δrsd | this work |
| HE806 | NQ1243, Δrsd:cat | titration of glucose uptake in Δrsd | this work |
| HE818 | NCM3722, $P_{oid}$-lacZ@nth | constitutive reporter near ter | this work |
| HE819 | NCM3722, $P_{oid}$-lacZ@essQ | constitutive reporter near ter | this work |
| HE820 | NCM3722, $P_{oid}$-lacZ@atpI | constitutive reporter near oriC | this work |
| HE821 | NCM3722, $P_{oid}$-lacZ@yeiN | constitutive reporter nearOriC | this work |
| NQ1389 | NCM3722, $pZE11_{ptetstab}$-LacZ | LacZ overexpression | Hui et al 2015 |

**Table S1a:** List of strains used in this study.

| **growth condition** | **strain** | **medium** | **growth rate ($h^{-1}$)** |
|---|---|---|---|
| Reference* | NCM3722 | M9, glucose | 0.91 |
| Catabolic limitation by titrating glucose uptake* | NQ1243 | M9, glucose, 400µM 3-MBA | 0.77 |
| | | M9, glucose, 100µM 3-MBA | 0.69 |
| | | M9, glucose, 0µM 3-MBA | 0.56 |
| | NQ1390 | M9, glucose, 400µM 3-MBA | 0.48 |
| | | M9, glucose, 40µM 3-MBA | 0.35 |
| | HE806 | M9, glucose, 400µM 3-MBA | 0.64 |
| | | M9, glucose, 0µM 3-MBA | 0.51 |
| | HE805 | M9, glucose, 40µM 3-MBA | 0.26 |
| Catabolic limitation by varying carbon sources | NCM3722 | MOPS, 20mM succinate | 0.59 |
| | | MOPS, 6mM mannose | 0.38 |
| | | MOPS, 4mM mannose | 0.32 |
| | | MOPS, 3mM mannose | 0.26 |
| | | MOPS, 20mM aspartate | 0.22 |
| | | MOPS, 20mM glutamate | 0.1 |
| | HE725 | MOPS, glucose | 0.87 |
| | | MOPS, glycerol | 0.55 |
| | | MOPS, 20mM mannose | 0.3 |
| | | MOPS, 4mM mannose | 0.2 |
| | | MOPS, 3mM mannose | 0.18 |
| | | MOPS ,20mM glutamate | 0.04 |
| | HE818, HE819, HE820, HE821 | MOPS, glucose, 20µg/ml kan | 0.91 |
| | | MOPS, 20mM succinate, 20µg/ml kan | 0.62 |
| | | MOPS, 30mM acetate, 20µg/ml kan | 0.43 |
| | | MOPS, 4mM mannose, 20µg/ml kan | 0.34 |
| Anabolic limitation | NQ393 | M9 glucose, 90µM IPTG | 0.77 |
| | | M9 glucose, 40µM IPTG | 0.69 |
| | | M9 glucose, 30µM IPTG | 0.49 |
| | | M9 glucose, 25µM IPTG | 0.32 |
| | | M9 glucose, 20µM IPTG | 0.27 |

| | | | |
|---|---|---|---|
| Translation limitation using chloramphenicol (Cm) | NCM3722 | M9 glucose, 0µM Cm | 0.91 |
| | | M9 glucose, 2µM Cm | 0.79 |
| | | M9 glucose, 4µM Cm | 0.61 |
| | | M9 glucose, 6µM Cm | 0.49 |
| | | M9 glucose, 8µM Cm | 0.36 |

**Table S1b:** List of growth conditions used in this study. *represents those titration series for which the biochemical measurements for mRNA synthesis rates were performed in MOPS media in addition to M9 medium.

### SUPPLEMENTAL METHODS

#### STRAINS AND GROWTH CONDITIONS

The *Escherichia coli* K-12 strain NCM3722 [38] was the parental strain used in this study. Strains and growth conditions used in this work are listed in Table S1. For all measurements, cultures were grown exponentially in the experimental condition for at least 6 generations before sampling. For the measurement of mRNA abundance by the hybridization method, the respective growth media were supplemented with 10 µg/ml uracil. For the growth conditions marked with (*), the biochemical measurements of mRNA and RNA synthesis rates and abundances were also confirmed in MOPS-buffered media [39] in addition to M9 media [40].

#### METHOD DETAILS

##### Absolute protein fractions

Quantitative measurements of *E. coli* proteomes under various growth conditions and determination of protein mass fractions were performed in Ref. [7]. In this work, we made use of the samples corresponding to exponential growing cultures in minimal media (C-limitation, A-limitation and R-limitation samples). Protein mass fractions were converted to protein number fractions (in Table S2) as described in Supplementary Note S1. Briefly, E. coli cell pellets were lysed with 2% sodium deoxycholate and digested with LysC and trypsin. An iRT peptide mix (Biognosys) was added to all samples for retention time alignment. Tryptic peptides were measured in SWATH mode (64 variable windows) on a TripleTOF 5600 mass spectrometer (Sciex). The DIA/SWATH data was analyzed using OpenSWATH (www.openswath.org). Protein mass fractions were computed from the detected peptide intensities using the xTop algorithm and a final calibration of xTop protein intensities with ribosome profiling data.

##### RNA extraction

Cells were pelleted from 8-10 ml cultures at $OD_{600}$ of ~ 0.4, resuspended in 200 µl of Lysis buffer A: (10 mM Tris-HCl, pH7.7 and 1 mg/ml lysozyme from OmniPur, cat. no. 596010GM), transferred to an RNase-free microtube, and incubated at 65°C for 1 minute. RNA was extracted by adding 200 µl of TRIzol reagent (Invitrogen, cat. no. 15596026) followed by incubation at room temperature for 5 minutes. 20 µl of chloroform was added and the samples were further incubated for 5 minutes, followed by centrifugation at 4,000 rpm for 30 seconds. The RNA-containing aqueous phase was transferred to a new RNase-free microfuge tube containing 20 µl of 3M sodium acetate, pH5.3 following which 1 ml ethanol was added, and RNA was precipitated at -20°C overnight. RNA was pelleted by centrifugation at 16,100 g for 20 minutes at 4°C, rinsed with 70% ethanol and dissolved in 30 µl of RNase-free water.

##### RNA sequencing

rRNA was depleted using the Ribo-Zero kit (MRZMB126, Illumina) as per the manufacturer's protocol. RNA-sequencing libraries were generated from the rRNA-depleted samples using TruSeq Stranded kit (Illumina) as per the manufacturer's protocol and each sample was barcoded uniquely. Libraries were pooled, single-end sequenced using Illumina Hi-seq 2500 and demultiplexed. Reads were aligned to the *E.coli* MG1655 U00096.3 genome using bowtie v2.2.6 [41]. Read counts were obtained using Python HTSeq-count (HTSeq v0.6.1p2) script [42].

**Northern Blotting**

1 μg each of RNA sample extracted as described above was mixed with known amounts (0.05 - 2 ng) of the appropriate spike RNA fragment (see below). The volume was adjusted to 9 μl with RNA sampling buffer (10 mM MOPS, 1 mM EDTA, 5 mM sodium acetate, 50% (vol/vol) formamide, and 2.2 M formaldehyde, pH7.0) and 1 μl loading dye (a faint amount of Bromophenol Blue (BPB) dissolved in water). The mixture was incubated at 72°C for 2 minutes, and then loaded onto formaldehyde agarose gels (containing 1.2% agarose, 20 mM MOPS, pH 7.0, 1 mM EDTA, 5 mM sodium acetate, 2.2 M formaldehyde, and 0.1 μg/ml ethidium bromide). Electrophoresis was carried out at 65 volts and stopped when the BPB band migrated to about three-quarters of the gel.

Following electrophoresis, the RNA was transferred to a nylon-based membrane filter (Zeta-Probe blotting membrane; cat #: 162-0153; Bio-Rad) by capillary action, using 20 x SSC buffer (3 M NaCl, 0.34 M sodium citrate, pH 7.0). The blots were fixed onto the membrane by exposure to UV for 2 minutes. The membrane was rinsed with water at room temperature for 5 minutes and then incubated at 42°C for 1.5 h in the pre-hybridization buffer (50 mM Tris, pH7.5, 1 M NaCl, 1 mg/ml sodium pyrophosphate, 50% (vol/vol) formamide, 0.1 mg/ml denatured salmon sperm single-stranded DNA and 2 x Denhardt reagent, which consists of 0.4 mg/ml Ficoll, 0.4 mg/ml polyvinylpyrrolidone, 0.4 mg/ml bovine serum albumin).

Meanwhile, 1 pmol of single-stranded DNA oligos probes were phosphorylated at 37°C for 1.5 h in a 20 μl reaction mixture containing 10 units of T4 polynucleotide kinase (Cat #: M0201, New England BioLabs), 1 x final concentration of the supplied buffer, and 100 μCi of γ-[$^{32}$P] ATP. The labeled probe was purified with a spin column (illustra MicroSpin G-50 columns, GE Healthcare) and diluted by the addition of 1 ml RNase-free water. 150 μl of the diluted probe was added to the membrane filter in the pre-hybridization buffer and incubated at 42°C overnight. The filter was washed twice in 0.1 x SSC containing 0.1% SDS at 42°C for 30 min, exposed overnight to FujiFilm BAS-MS4020 imaging plate scanned using a Biorad PMI personal molecular imager FX. Image analysis was performed using Image J software.

**In-vitro transcription of target mRNA spike-in**

The first 400 base pairs of the different target genes were amplified from the *E.coli* genomic DNA to generate linear DNA templates and gel-purified. Using these linear templates, in-vitro transcription was carried out using the Ampliscribe T7-flash kit (Epicenter) according to the manufacturer's protocol. The in-vitro synthesized RNA was gel-purified and quantified using the Qubit fluorometer (Invitrogen) as well as Tapestation (Agilent).

**Genomic DNA extraction, denaturation and immobilization**

*E. coli* genomic DNA (gDNA) was extracted from 12 ml cultures of NCM3722 growing in minimal media with 4 mM mannose as the sole carbon source. Cells were pelleted and lysed in 1 ml gDNA Lysis buffer B: (10 mM Tris HCl at pH 7.7, 1 mM EDTA, 1% SDS and 1mg/ml proteinase K) at 37°C for 1 hr. Equal volume of 1:1 phenol:chloroform mixture was added to the lysate and incubated for 10 minutes at room temperature. The aqueous phase was transferred to a fresh tube and the extraction using 1:1 phenol chloroform was repeated. The gDNA was then precipitated with ethanol in the presence of 300 mM sodium acetate (pH 5.3) at -20°C overnight, rinsed with 70% ethanol and dissolved in nuclease-free water.

The gDNA was first denatured as described by Pigott 1968 [43] and immobilized on nitrocellulose membranes as described by Gillespie and Spiegelman 1965 [44], with some minor adaptations. The gDNA was diluted to 100 μg/ml with 0.01 x SSC and denatured by adding 1N NaOH to a final strength of approximately 0.15N and incubating at room temperature for 10 minutes. The samples were then cooled on ice for 5 minutes and then neutralized using 5N acetic acid, verified using a pH strip. Ice cold denatured gDNA was passed through Whatman cellulose nitrate membrane filter (GE healthcare, cat. 7184-009) at a slow rate of approximately 40 minutes using a Schleicher and Schuell dot-blot assembly such that each well received 5 μg of gDNA. The membrane was then exposed to UV for 2 minutes, dried and the area corresponding to each well was cut out in the form of discs.

**Measurement of total mRNA abundance**

Overall, the method was adopted from Freissen 1966 [14], with minor adaptations. The procedure is illustrated in Fig. S5A. Cultures for various growth conditions were typically started at $OD_{600}$ ~ 0.006 in the presence of 1 μg/ml unlabeled uracil. When the $OD_{600}$ reached 0.05 (approximately 3 generations), 2 μCi of $^3$H-uracil (at 30-50 Ci/mmol, Perkin Elmer) was added to a culture volume of 600 μl and growth was allowed to continue. At various values of $OD_{600}$, 100 μl of labelled cultures were removed into 100 μl of boiling Lysis buffer C (0.1 M NaCl, 0.01 M Tris HCl at pH 7.7, 0.02 M EDTA, 0.5% SDS) and incubated at 100°C for 2 minutes. The lysates were then cooled on ice, treated with 40 units of RNase-free DNase I (Roche) for 10 minutes followed by purification of RNA using the RNeasy kit (Qiagen) as per the vendor's protocol. The resulting RNA samples devoid of excess un-incorporated label were then brought to a total volume of 100 μl with nuclease free water. 150 μl of Hybridization solution (280 μl Denhardt reagent as described above, 20 μl of salmon sperm single stranded DNA that is heated at 95°C for 3 minutes, volume adjusted to 2 ml with 6x SSC) was added to the samples and the RNA was hybridized to the gDNA by immersing a filter disc with immobilized gDNA and incubating overnight at 65°C. The hybridization reactions were cooled on ice and the filter discs were washed twice with 1 ml cold 6x SSC for 45 minutes each. The washed filter discs were dissolved in 2ml Liquiscint scintillation cocktail (National Diagnostics) and counts were read in a Beckman LS 6500 scintillation counter. The counts were corrected for the hybridization efficiency, which was estimated to be ~75%, comparable to that observed previously [43]. The relative slopes of counts versus $OD_{600}$ between different growth conditions indicate the relative mRNA abundance per $OD_{600}$ of the cultures. While only three or four growth conditions could be measured on a given day, the reference growth condition (NCM3722 in MOPS-glucose) was always included, allowing for the data integration across all growth conditions. Finally, by using the mRNA/RNA ratio measured by Northern blotting and RNA-seq (described above), and total

RNA abundance [6], the relative mRNA abundances could be scaled to absolute abundances (Fig. S5H).

**Measurement of total mRNA and total RNA synthesis rates**

The procedure, adopted from Pigott 1968 [43] with some modification is illustrated in Fig. S9. Cultures were grown in the appropriate growth condition until OD600 ~ 0.3 and 10 μCi of $^{3}$H-uracil (at 30-50 Ci/mmol, Perkin Elmer) was added to 2 ml culture, following which 100 μl of aliquots were removed at 10 second intervals into 100 μl of boiling Lysis buffer C (0.1 M NaCl, 0.01 M Tris Hcl, pH 7.7, 0.02 M EDTA, 0.5% SDS) over a period of 1 minute. Each aliquot was incubated at 100°C for 2 minutes following which they were cooled on ice. The lysates were divided into 2 equal fractions, one for the measurement of mRNA synthesis rate and the other for total RNA synthesis rate. 50 μl of 20x SSC was added to the first fraction for mRNA synthesis and hybridization to gDNA was carried as described above in the measurement of mRNA abundance. To the fraction saved for measuring the total RNA synthesis, 50 μl of ice cold 10% TCA was added and the RNA was allowed to precipitate for 30 minutes. The precipitated RNA was collected by spinning at 13,000 rpm for 15 minutes, dissolved in 100 μl water, added to 2 ml of Liquiscint scintillation cocktail (National Diagnostics) and counted in a Beckman LS 6500 scintillation counter. The relative slopes of counts incorporated over time for different growth conditions indicate either the relative mRNA or the total RNA synthesis rates across growth conditions. In order to integrate data for different growth conditions collected over several days, a reference growth condition (NCM3722 growing in MOPS glucose) was included every time.

**Measurement of mRNA turnover**

The procedure of Chen 2015 [45] was employed. Briefly, cultures were grown to $OD_{600}$ ~ 0.4 and Rifampicin (Sigma) dissolved in DMSO was added at a final concentration of 500 ng/μl. 10 ml cultures were removed at 0, 1, 2, 3, 5, 7, 9 and 11 minutes of Rifampicin addition and cultures were inhibited with 10% volume of 9:1 ethanol:phenol solution. RNA preparation, RNA-sequencing and analysis was carried as described above.

**LacZ reporter assay for constitutive expression**

Beta-galactosidase activity was measured as described by You *et al* [33].

#### QUANTIFICATION AND STATISTICAL ANALYSIS

**Quantification of mRNA degradation rates**

In order to determine the mRNA degradation rates, we first computed the change in mRNA concentration over time, $[mR_i](t)/[mR_i](0)$. By writing the mRNA concentrations as the product of the mRNA fractions and the total mRNA concentration, this quantity can be expressed as follows:

$$\frac{[mR_i](t)}{[mR_i](0)} = \frac{\psi_{\mathrm{m},i}(t)}{\psi_{m,i}(0)} \cdot \frac{[mR](t)}{[mR](0)}$$

The mRNA fractions $\psi_{\mathrm{m},i}(t)$ at time $t$ are directly computed via RNA-sequencing. Instead, the change in the total mRNA concentration over time, $[mR](t)/[mR](0)$, has to be estimated separately. The logarithm of this ratio was modeled with a quadratic polynomial:

$$\log_2\left(\frac{[mR](t)}{[mR](0)}\right) = -t/\tau_{\mathrm{HL}} + ct^2$$

In this expression, $\tau_{\mathrm{HL}}$ corresponds to the half-life of the total mRNA concentration; the quadratic term takes into account the fact that as time passes by, the mRNA pool is progressively dominated by long-lived mRNAs, and hence the decay of total mRNA is not expected to be a simple exponential. The coefficients $\tau_{\mathrm{HL}}$ and c were determined using two criteria. First, the concentration of stable mRNAs should be time-independent. Therefore, the mRNA fractions of stable mRNAs should be inversely related to the change in total mRNA. Second, the abundance of mRNAs corresponding to genes distant from the promoter should be constant on a short time-scale, and then decrease. We used for the parameters the following values:

- Reference condition: $\tau_{\mathrm{HL}} = 1.8$ min, $c = 0.015\ \mathrm{min}^{-2}$
- Carbon limitation: $\tau_{\mathrm{HL}} = 2.2$ min, $c = 0.01\ \mathrm{min}^{-2}$

The corresponding curves are shown in red in Figure S7B. The trajectories for stable mRNAs and genes far from promoters are also shown for comparison. Using these parameters, we computed the scaled mRNA concentrations, $[mR_i](t)/[mR_i](0)$, which were fitted to a function describing a lagged exponential decay. More precisely, we modeled the mRNA degradation as follows:

$$\log\frac{[mR_i](t)}{[mR_i](0)} = \begin{cases} 0 & t < t_{\mathrm{lag},i}\ , \\ -\delta_i(t - t_{\mathrm{lag},i}) & t \geq t_{\mathrm{lag},i}\ , \end{cases}$$

where $\delta_i$ is the mRNA degradation rate and $t_{\mathrm{lag},i}$ is a lag time between rifampicin injection and the start of mRNA degradation. We only fitted genes for which the mRNA mass fraction (mass of the $i^{\mathrm{th}}$ mRNA over the total mRNA mass) was larger than $10^{-5}$ before adding rifampicin (this value corresponds roughly to the detection limit of our proteomics dataset, see Ref. [7], and with at least three data points with mass fractions above $10^{-8}$.

As can be seen in Figure S7C, for most genes the mRNA abundance drops by about two orders of magnitude within a few minutes, but degradation appears to stop afterwards. This behavior is not captured by the function used to model mRNA degradation. Instead, we performed a weighted fit with weights $w$ set according to the relative mRNA abundance, $y_i(t) \equiv [mR_i(t)]/[mR_i(0)]$:

$$w(y_i) = \begin{cases} 10^{-3} & y_i < 2\cdot 10^{-2} \\ 10^{-3} + (1 - 10^{-3})\cdot\left(\dfrac{y_i - 2\cdot 10^{-2}}{5\cdot 10^{-2} - 2\cdot 10^{-2}}\right) & 2\cdot 10^{-2} \leq y_i < 5\cdot 10^{-2} \\ 1 & y_i \geq 5\cdot 10^{-2} \end{cases}$$

With this choice, the fit prioritizes fitting the initial part of the mRNA decay, while mostly ignoring the long-term behavior of the relative abundances. Given the discontinuous nature of the function, it is necessary to perform the fit separately for lag times within each sampling time interval (between 0 and 1 minute, between 1 and 2 minutes, and so on), including the case $t_{\mathrm{lag},i} = 0$. After performing all fits, the one that minimizes the (weighted) sum of the squared residuals was selected. In order to improve the quality of the resulting dataset, results were discarded if such sum of squared residuals was larger than 1 or if the fitted mRNA degradation rate was larger than 5/min. Results for the mRNA degradation rates are reported in Table S4.

**Quantification of stable RNA synthesis flux and rRNA promoter strength**

The promoter strength of ribosomal RNA genes was computed as follows. The synthesis flux of ribosomal RNA, $A_{\mathrm{rR}}$, was computed as $A_{\mathrm{rR}} = 0.9 \cdot \lambda W_{\mathrm{R}}$, where $\lambda$ is the growth rate, $W_{\mathrm{R}}$ the total RNA abundance in units of RNA mass per culture density [17], and the factor 0.9 accounts for the fact that about 90% of the cellular RNA is ribosomal RNA, with the remainder being tRNA and mRNA [5]. Each transcription initiation leads to the synthesis of all main RNA components of the ribosome, namely the 16S, 23S and 5S rRNA. In total, these rRNAs correspond to a total length of $\ell_{\mathrm{rR}} = 4566$ nt. Therefore, in order to obtain the total transcription rate, $J_{\mathrm{rR}}$, we divided the rRNA synthesis flux by the mass corresponding to the 16S+23S+5S RNA, i.e. $J_{\mathrm{rR}} = A_{\mathrm{rR}}/m_{\mathrm{nt}}\ell_{\mathrm{rR}}$. This initiation flux is generated from the seven ribosomal operons *rrnABCDEGH*, for which we assume the same dependence on promoter on-rates and available RNAP polymerase as for the protein-coding genes. This allows to write $J_{\mathrm{rR}} = \sum_i k_i[G_i]\,[RNAP]$, with the sum running over the seven *rrn* operons. Further assuming identical regulation for the seven operons, $k_i \equiv k_{\mathrm{rrn}}$, we computed the promoter on-rate as $k_{\mathrm{rrn}} = J_{rR}/[RNAP]_{av}\sum_i[G_i]$, with the RNAP concentration shown in Fig. 6B and the gene concentrations computed from the known locations of the *rrn* operons and the concentration of Ori.

### SUPPLEMENTAL NOTES

#### Supplemental Note S1 – Unit conversions

In this Note we discuss the conversions across different concentrations units.

##### Protein and mRNAs fractions

We define the number fractions of the $i^{\text{th}}$ mRNA ($\psi_{\text{m},i}$) or protein ($\psi_{\text{p},i}$) as the number of mRNAs or proteins ($N_{\text{mR},i}$, $N_{\text{P},i}$) in the culture over the total number of mRNAs and proteins ($N_{\text{mR}}$, $N_{\text{P}}$):

$$\psi_{\text{m},i} = \frac{N_{\text{mR},i}}{\sum_k N_{\text{mR},k}} = \frac{N_{\text{mR},i}}{N_{\text{mR}}} \quad , \quad \psi_{\text{p},i} = \frac{N_{\text{P},i}}{\sum_k N_{\text{P},k}} = \frac{N_{\text{P},i}}{N_{\text{P}}} \qquad \text{(N1.1)}$$

Since the number fractions are ratios of abundances, these can also be expressed as ratios of cellular concentrations by dividing both numerator and denominator by the total cellular volume, $V_{\text{cell}}$. The number fractions are often the most direct output of –omics measurements. For instance, RNA sequencing output reported in TPM are proportional to mRNA number fractions (see below); similarly, protein intensities are usually defined so that they yield protein number fractions when normalized to the total protein intensity.

While –omics measurements allow to compute fractional quantities, total or bulk measurements of protein and mRNA abundances most usually yield total mass per culture unit. For instance, the Biuret method, commonly used for the determination of protein abundance, provides a signal proportional to the number of peptide bonds in the sample, which in turn is proportional to the total number of protein residues in the culture. Therefore, it is convenient to introduce mass fractions of mRNA and proteins:

$$\phi_{\text{m},i} = \frac{m_{\text{m},i} N_{\text{mR},i}}{\sum_k m_{\text{m},k} N_{\text{mR},k}} = \frac{m_{\text{m},i} N_{\text{mR},i}}{W_{\text{mR}}} \quad , \quad \phi_{\text{p},i} = \frac{m_{\text{p},i} N_{P,i}}{\sum_k m_{\text{p},k} N_{P,k}} = \frac{m_{\text{p},i} N_{P,i}}{W_P} \qquad \text{(N1.2)}$$

where $m_{\text{m},i}$ and $m_{\text{p},i}$ are the mass of the $i^{\text{th}}$ mRNA and protein, respectively, and $W_{\text{mR}}$ and $W_P$ are the total mRNA and protein mass per culture unit (units: μg/$\text{OD}_{600}$mL), respectively.

In this work we approximated the mass of each protein as $m_{\text{p},i} \approx m_{\text{aa}} \ell_i$, where $\ell_i$ is the number of residues of the protein and $m_{\text{aa}}$ is the average mass of an amino acid in the proteome, $m_{\text{aa}} = 1.83 \cdot 10^{-16}$ μg. Similarly, we approximated the mass of each mRNA as $m_{\text{m},i} = 3 m_{\text{nt}} \ell_i$, where $m_{\text{nt}} = 5.38 \cdot 10^{-16}$ μg and the factor 3 is due to the coding ratio of three nucleotides per residue. In doing so, we are neglecting regions upstream and downstream of the protein coding region of the mRNAs, which are generally small compared to the mRNA length. This approximation allows us to very simply convert between number and mass fractions, and vice versa. Mass fractions are obtained as:

$$\phi_{\mathrm{m},i} = \frac{\ell_i \psi_{\mathrm{m},i}}{\sum_k \ell_k \psi_{\mathrm{m},k}} \quad , \quad \phi_{\mathrm{p},i} = \frac{\ell_i \psi_{\mathrm{p},i}}{\sum_k \ell_k \psi_{\mathrm{p},k}} \ , \tag{N1.3}$$

while number fractions are obtained from number fractions as:

$$\psi_{\mathrm{m},i} = \frac{\phi_{\mathrm{m},i}/\ell_i}{\sum_k \phi_{\mathrm{m},k}/\ell_k} \quad , \quad \psi_{\mathrm{p},i} = \frac{\phi_{\mathrm{p},i}/\ell_i}{\sum_k \phi_{\mathrm{p},k}/\ell_k} . \tag{N1.4}$$

The denominators of the expressions in Eq. N1.3 also appear frequently when converting total amounts of proteins or mRNAs into concentrations. It is hence convenient to define the mRNA-weighted and protein-weighted gene lengths (expressed in codons):

$$\bar{\ell}_{\mathrm{mR}} \equiv \sum_k \ell_k \psi_{\mathrm{m},k} , \qquad \bar{\ell}_{\mathrm{P}} \equiv \sum_k \ell_k \psi_{\mathrm{p},k} \tag{N1.5}$$

In particular, the average protein size is given by $\bar{\ell}_{\mathrm{P}}$; it is shown, together with the average protein mass $\bar{m}_{\mathrm{P}} = m_{\mathrm{aa}} \bar{\ell}_{\mathrm{P}}$, in Fig. S2E for carbon limitation and rich media, and has a very mild dependence on growth rate.

**Calculating mRNA fractions from common RNA-sequencing units**

For single-end sequencing, common units used to report mRNA abundances are RPM (read counts per million), RPKM (read counts per million, per kilobase) and TPM (read counts per kilobase, per million). Their relation with number and mass mRNA fractions, which are more convenient for our purposes, is summarized as follows. (The analogous relations between protein intensities generated from mass-spectrometry pipelines and protein number or mass fractions are explained in detail in Ref. [7].

Comparing mRNA of different lengths but present in the same number, it is clear that longer genes will have a proportionally more mapped reads (in absence of other sources of bias). Therefore, we take the read counts to be proportional to the total length (mass) of each mRNA in the sample of interest: it then follows that RPM (reads per million) are proportional to the mRNA mass fractions. Defining $R_i$ to be the number of reads mapped to gene $i$:

$$\mathrm{RPM}_i \equiv 10^6 \frac{R_i}{\sum_k R_k} = 10^6 \phi_{\mathrm{m},i} . \tag{N1.6}$$

Similarly, TPM (reads per kilobase, per million) are defined by first normalizing the read counts by the gene length (the unit for the gene length does not matter) and then normalizing so that the sum of the TPM counts equals $10^6$. Since the read counts are themselves proportional (in a given sample) to the mass fractions, $R_i \propto \phi_i$, and comparing the definition of TPMs with Eq. (N1.4), it is clear that TPM counts are proportional to the mRNA number fractions:

$$\mathrm{TPM}_i \equiv 10^6 \frac{R_i/\ell_i}{\sum_k R_k/\ell_k} = 10^6 \psi_{\mathrm{m},i} . \tag{N1.7}$$

The $10^6$ factors (the "per million" in the unit names) are usually convenient for human readability, but not when the abundances of mRNAs have to be combined mathematically with other quantities, e.g. with the total mRNA concentration to yield the concentrations of individual mRNAs. For this reason we use number and mass fractions instead of TPM and RPM counts, respectively. The similitude between Eq. (N1.7) and Eq. (N1.4) above is not incidental: the conversion between RPM and TPM mirrors exactly the conversion between mass and number fractions summarized in Eq. (N1.3)-(N1.4). Finally, RPKM counts are defined as reads per million, per kilobase:

$$\mathrm{RPKM}_i \equiv \frac{10^6}{\ell_i} \frac{R_i}{\sum_k R_k} = \frac{10^6 \phi_{\mathrm{m},i}}{\ell_i} . \tag{N1.8}$$

From the definition, it is clear that RPKM counts are a non-normalized version of the mRNA number fractions, as can be seen by comparing with Eq. (N1.4). Since this quantity is not normalized, it cannot be straightforwardly used to compare mRNA abundances across different samples, and is therefore less useful than RPM and TPM.

**Cellular concentrations and copy numbers per cell**

Optical density per culture volume (OD·mL) is a convenient tool to quantify the abundance of cell culture in batch tube experiments. Optical density is known to correspond well to cellular dry mass [6], however it is not *per se* an absolute unit. For example, the measured optical density can vary, e.g., depending on the instrument used for the measurement. Rather, the ratio of two quantities measured per unit of culture volume (e.g. the total protein mass and the total dry weight) does not depend on the culture volume itself.

For any quantity $X$ (including proteins and mRNAs), we indicate abundances per culture unit and "average" concentrations as $N_X$ and $[X]$, respectively. The ratio of the two is given by the total cell volume per OD mL, $V_{\mathrm{cell}}$:

$$\frac{N_X}{[X]} = V_{\mathrm{cell}} . \tag{N1.9}$$

This latter quantity can be broken down into the product of the average cell volume, $v_{\mathrm{cell}}$, and the number of cells per OD mL, $N_{\mathrm{cell}}$:

$$V_{\mathrm{cell}} = N_{\mathrm{cell}} \cdot v_{\mathrm{cell}} . \tag{N1.10}$$

The two have been measured independently [6], and are reported in Fig. S2A-B. Together suggest a very weak growth rate dependence of $V_{\mathrm{cell}}$ across conditions, see Fig. S2C.

In this work we only consider the "average" cellular concentrations (copy number per cell volume). However, it should be noted that average concentrations can differ from the "actual" concentrations needed for biochemical calculations, for a couple of reasons. A first issue is the presence of compartments: proteins can be localized into either the cytoplasm or the periplasm, and the two cellular compartments have different (condition-dependent) volumes [46]. This implies that the average concentration of a protein across the whole cell can be quite different from the actual concentration in the cellular compartment. Additional, concentrations of membrane- or wall-associated proteins have to be calculated with either the cytoplasmic or the periplasmic volume, depending on the application. In this work we do not distinguish among cellular compartments, and we hence adopt the "average" concentration as concentration unit.

A second problem is that both cytosol and periplasm are crowded environment, which limits the volume accessible to the solvent (water). Therefore, concentrations should only take into account the volume of water, without considering the excluded volume of macromolecules (proteins, RNA, DNA); furthermore, the water volume can potentially depend on the growth condition, incuding the osmolarity of the medium. However, it can be seen that average concentrations of cytoplasmic proteins are indeed proportional to cytoplasmic concentrations. Indeed, the buoyant cell density of *E. coli* does not depend on the growth rate [8], implying the approximate proportionality between dry mass and the cytoplasmic water volume. Together with the approximate constancy of cell dry mass per culture volume [6], and the weak growth dependence of the total cell volume per culture unit (Fig. S2C), we deduce that the cytoplasmic water volume per cell volume to depend weakly on the growth rate. This directly leads to an approximate proportionality across growth rates between average and cytoplasmic concentrations of cytoplasmic proteins.

Finally, "per cell" quantities are instead obtained as $N_X/N_{\mathrm{cell}} = [X]/v_{\mathrm{cell}}$. This normalization can be used, for instance, to express mRNA and protein copy numbers per cell. However, the cell size depends strongly on growth conditions (Fig. S2B), and therefore a copy number of mRNAs and proteins will correspond to very different concentrations across growth conditions. For this reason, we do not use the "per cell" normalization, and stick to concentration units throughout the study.

**Protein concentration**

The concentrations of individual proteins can be expressed as the product of the total protein concentration $[P]$ and the protein fraction $\psi_{\mathrm{p},i}$, i.e.

$$[P_i] = \psi_{\mathrm{p},i}[P]\,. \tag{N1.11}$$

The total protein concentration was determined by combining the measured values of total cell volume per culture unit ($V_{\mathrm{cell}} = N_{\mathrm{cell}}v_{\mathrm{cell}}$, Fig. S2C), protein mass per culture unit ($W_{\mathrm{P}}$, Fig. S2D) and average protein mass $\bar{m}_{\mathrm{P}}$ (Fig. S2E) as

$$[P] = \frac{W_{\mathrm{P}}}{\bar{m}_{\mathrm{P}}V_{\mathrm{cell}}}\ . \tag{N1.12}$$

The resulting values, shown in Fig. S2F, do not show a significant dependence on growth rate. Hence, we take its average value across conditions as basic constant, which we use to derive protein concentrations across all conditions in this work:

$$[P] \approx (3.15 \pm 0.1) \cdot 10^6/\mu m^3\ . \tag{N1.13}$$

Eq. N1.8 and N1.10 allow to readily convert protein fractions $\psi_{\mathrm{p},i}$ into protein concentrations. For example, a fraction $\psi_{\mathrm{p},i} = 10^{-4}$ is equivalent to a concentration of about $300$ proteins per $\mu\mathrm{m}^3$.

**mRNA concentration**

The total concentration of mRNAs shown in Fig. 2A is obtained in a manner similar to what done for the protein abundances. In this work we measured the total mRNA mass per culture unit, $W_{\mathrm{mR}}$, by combining relative measurements of uracil incorporation and DNA hybridization and the absolute measurements yielded by the joint use of RNA-seq and quantitative Northern blotting (Fig. S5 and Methods). The resulting values for $W_{\mathrm{mR}}$ shown in Fig. S5H were converted to total mRNA concentrations using the total cellular volume per culture $V_{\mathrm{cell}}$ and the average mRNA mass $\bar{m}_{\mathrm{mR}} = 3m_{\mathrm{nt}} \sum_i \ell_i \psi_{\mathrm{m},i}$ as follows:

$$[mR] = \frac{W_{\mathrm{mR}}}{\bar{m}_{\mathrm{mR}} V_{\mathrm{cell}}} . \tag{N1.14}$$

Notably, while the total protein concentration can be taken to be constant in our conditions, the total mRNA concentration depends strongly on the growth rate. The concentration of individual mRNAs is hence obtained as:

$$[mR]_i = \psi_{\mathrm{m},i}[mR] . \tag{N1.15}$$

An alternative, but perfectly equivalent, way of computing the mRNA concentrations is to multiply the total mRNA mass per cell volume, $W_{\mathrm{mR}}/V_{\mathrm{cell}}$, with the mRNA mass fraction $\phi_{\mathrm{m},i}$ to obtain the mass of $i^{\mathrm{th}}$ mRNA species per cell volume; this can then be divided by the mass of a single mRNA (which is approximately given by $m_{\mathrm{mR},i} = 3m_{\mathrm{nt}}\ell_i$) to yield the concentration of mRNA.

**Gene concentration**

The gene concentrations are set by the so-called C and D periods. These quantities are central in the study of cell replication, and have been studied in detail in the last decades [27, 29].

The C period sets the relative abundance of genes at different locations on the chromosome. Since the chromosome replication proceeds from the OriC locus to Ter, genes close to OriC will have higher copy numbers compared to genes close to Ter. To quantify this effect, it is convenient to introduce $x$ as the distance of a gene from the origin of replication OriC, with $x = 0$ corresponding to OriC itself and $x = 1$ corresponding to the maximal distance for genes close to Ter. With this definition, the concentration of a gene $i$, $[G_i]$, is given by:

$$[G_i] = [Ori] \cdot e^{-x_i \cdot (\lambda C)} . \tag{N1.16}$$

This can also be expressed in terms of the gene dose, defined as $g_i \equiv e^{-x_i \cdot (\lambda C)}$:

$$[G_i] = [Ori] \cdot g_i . \tag{N1.17}$$

While the gene dose depends only on the growth rate $\lambda$, the C period and the distance from Ori of the gene under consideration ($x_i$), the concentration of Ori further depends on the D period. The latter is the segregation time between the completion of chromosome replication and cell division and, together with the C-period, sets the average number of chromosome copies per cell when averaging over a cellular population:

$$n_{\mathrm{Ori}} = e^{\lambda(C+D)}\ . \tag{N1.18}$$

To estimate the gene concentrations, we primarily used data from Zheng *et al.* [27]. In this work, the authors measured a variety of cell replication parameters, including both the C and D periods, the number of Ori per cell $n_{\mathrm{Ori}}$, and the average cell volume $v_{\mathrm{cell}}$, for *E. coli*. In particular, $\lambda C$ was found to be linearly related to the growth rate, $\lambda C = a + b\lambda$, which allows to estimate the gene dose $g_i$ across growth rates (Fig. S9A-B). Furthermore, the measurements of both $n_{\mathrm{Ori}}$ and $v_{\mathrm{cell}}$ allowed to determine the origin concentration $[Ori] = n_{\mathrm{Ori}}/v_{\mathrm{cell}}$ (Fig. S9C-E). Together, the gene dose and the concentration of Ori allow to compute the concentration of each gene $[G_i]$ via Eq. N1.17.

### Supplemental Note S2 – Protein homeostasis and ribosome density

This note complements and extends Main Text Fig. 2 and Fig. S6, providing additional details on the derivation of the constraint between mRNA abundance and translating ribosomes (or alternatively, between translation initiation and elongation rates) discussed in the Main Text. The starting point of the analysis is the formulation of protein kinetics (Eq. (S1) in Fig. S1) in terms of translation initiation flux, $\alpha_{\mathrm{p},i}[\mathrm{mR}_i]$, and protein turnover, $\lambda[P_i]$:

$$\frac{\mathrm{d}[P]}{\mathrm{d}t} = \alpha_{\mathrm{p},i}[\mathrm{mR}_i] - \lambda[P_i]\,. \qquad \text{(N2.1)}$$

This expression, when evaluated at steady state, yields a constraint relating mRNA and protein abundances, the translation initiation rate and the growth rate:

$$\alpha_{\mathrm{p},i}[\mathrm{mR}_i] = \lambda[P_i]\,. \qquad \text{(N2.2)}$$

A point worth mentioning is that, in individual cells, gene expression is noisy. For instance, for lowly expressed genes, mRNA copy numbers can oscillate between zero and one per cell, leading to bursts of protein synthesis of short duration, followed by lack of protein synthesis. Therefore, for individual cells, the right hand side of Eq. (N2.1) cannot be simply set to zero to yield Eq. (N2.2). However, it is possible to do so if $[mR_i]$ and $[P_i]$ are average concentrations across the whole cell culture. In this case, the steady state assumption is simply that of balanced growth (i.e. average or bulk quantities are time-independent). In this work we will always refer to average concentrations over a population of cells.

**Protein synthesis, dilution and turnover**

A central assumption in our analysis is that the synthesis rate of the *i*th protein is given by $\lambda[P_i]$. Protein turnover for *E. coli* cells in exponential growth is generally very small [3, 4]. As done in Mori et al [7], we assumed that the bulk (average) protein degradation rate is negligible compared to the growth rate, and we approximated the protein synthesis flux for the *i*th protein as the product of its concentration $[P_i]$ and the growth rate $\lambda$. Under this assumption, the protein fraction $\psi_{\mathrm{p},i}$ also equals the fraction of protein synthesis flux (proteins synthesized per unit of time) associated to the *i*th protein.

While this approximation is generally valid for exponentially growing *E. coli*, it underestimates the synthesis rates of proteins subject to fast degradation, or proteins that are excreted extracellularly. This is the case for a few proteins, most notably flagellin (FliC) and the sigma factors $\sigma^S$ and $\sigma^H$ [7] [47]. Also, this assumption is not valid in organisms in which protein degradation happens at rates comparable to growth, for instance in eukaryotic cells.

**Translation initiation rate, translation efficiency**

We rewrite Eq. N2.2 rewriting the protein and mRNA concentrations in terms of fractions and total concentrations, as follows:

$$\alpha_{\mathrm{p},i}\psi_{\mathrm{m},i}[mR] = \lambda[P]\psi_{\mathrm{p},i}\,. \qquad \text{(N2.3)}$$

Solving for the translation initiation rate $\alpha_{\mathrm{p},i}$ yields:

$$\alpha_{\mathrm{p},i} = \frac{\psi_{\mathrm{p},i}}{\psi_{\mathrm{m},i}} \times \frac{\lambda[P]}{[mR]} \cdot \qquad \text{(N2.4)}$$

This expression can be broken down into two terms. The first term, $\psi_{\mathrm{p},i}/\psi_{\mathrm{m},i}$, is the ratio of protein and mRNA number fractions, and it is gene-specific. It provides a relative measurement of the initiation rate $\alpha_{\mathrm{p},i}$, i.e. it allows to compare the initiation rates of *different* mRNAs in the *same* growth condition. For this reason, the ratio of protein (synthesis rates) and mRNA number fractions has been simply named "translation efficiency" [2].

The second term provides the absolute scale of the mRNA initiation efficiencies. Its meaning can be appreciated by summing Eq. N2.3 over all genes, yielding

$$\bar{\alpha}_{\mathrm{p}}[mR] = \lambda[P]\,, \qquad \text{(N2.5)}$$

where we defined

$$\bar{\alpha}_{\mathrm{p}} \equiv \sum_i \alpha_{\mathrm{p},i}\psi_{\mathrm{m},i} = \frac{\lambda[P]}{[mR]}. \qquad \text{(N2.6)}$$

Therefore, the second term in Eq. N2.4 is equal to the average translation initiation rate within the mRNA population. Together, Eq. (N2.4) and (N2.6) allow to write the initiation rates $\alpha_{\mathrm{p},i}$ in terms of the average initiation rate $\bar{\alpha}_{\mathrm{p}}$ and the translation efficiency $\psi_{\mathrm{p},i}/\psi_{\mathrm{m},i}$:

$$\alpha_{\mathrm{p},i} = \bar{\alpha}_{\mathrm{p}} \cdot \frac{\psi_{\mathrm{p},i}}{\psi_{\mathrm{m},i}}. \qquad \text{(N2.7)}$$

Since $\bar{\alpha}_{\mathrm{p}}$ is in principle condition-dependent, it has to be measured for cells growing in different conditions in order to compare initiation rates $\alpha_{\mathrm{p},i}$ of the *same* gene *across conditions*. In our study, we provided extensive mass spectroscopic and RNA-sequencing data which show that most translation efficiencies $\psi_{\mathrm{p},i}/\psi_{\mathrm{m},i}$ are condition independent, and for the bulk of the genes they reside within a 2-3 fold range around unity (Fig. 2, Supp. Fig. S1-S2). The initiation rates $\alpha_{\mathrm{p},i}$ are consequently proportional, and numerically similar, to their average value $\bar{\alpha}_{\mathrm{p}}$. On the other hand, $\bar{\alpha}_{\mathrm{p}}$ depends weakly on the growth rate (Fig. 2B).

**Ribosomal activity and ribosome density**

The process of protein synthesis requires two main components: mRNAs and ribosomes. Equations N2.1 and N2.2 (same as Eq. S1 and S3 in Supp. Fig. S1) describe translation through a "microscopic", mRNA-centric point of view, which allows to study the protein synthesis of individual proteins from the corresponding mRNAs, including the quantification of their translation efficiencies. However, these expressions overlook the global effect of macromolecular machinery (ribosomes) on translation. In order to relate protein synthesis to the concentration of translating ribosomes, we note that the number of residues elongated per unit of time in a cell equals the elongation rate of the ribosomes $\varepsilon$ (in codons/second; the elongation rate is mostly gene-independent, despite the occasional presence of slow codons [17] times the concentration of actively translating ribosomes $[Rb]_{\text{act}}$ (Supp. Fig. S6A). On the other hand, the total protein flux can be written as the sum over all genes of the protein synthesis rate, $\lambda[P_i]$, and the protein length $\ell_i$, yielding:

$$\sum_i \lambda[P_i]\ell_i = \lambda[P]\bar{\ell}_{\text{P}} = \varepsilon[Rb]_{\text{act}}\,, \qquad \text{(N2.8)}$$

where $\bar{\ell}_{\text{P}} = \sum_i \psi_{\text{p},i}\ell_i$ is the average protein length. Using Eq. N2.5, and defining $\bar{r}$ as the ratio between the concentration of translating ribosomes and that of mRNAs, we obtain the following simple relation between the average initiation rate and the number of mRNAs per mRNA:

$$\bar{r} \equiv \frac{[Rb]_{\text{act}}}{[mR]} = \frac{\bar{\ell}_{\text{P}}\bar{\alpha}_{\text{P}}}{\varepsilon}\,. \qquad \text{(N2.9)}$$

To appreciate the meaning of this expression, consider that the density of ribosomes on an mRNA is given by the ratio of the mRNA initiation rate, $\alpha_{\text{p},i}$, and the elongation rate $\varepsilon$, with the average spacing (in codons) between ribosomes given by the reciprocal of this quantity, $d_i = \varepsilon/\alpha_{\text{p},i}$. Therefore, the ratio $\bar{\alpha}_{\text{P}}/\varepsilon$ represents the average density of ribosomes on mRNAs, to which corresponds an average spacing

$$\bar{d} \equiv \frac{\varepsilon}{\bar{\alpha}_{\text{P}}} = \frac{\bar{\ell}_P[mR]}{[Rb]_{\text{act}}}\,. \qquad \text{(N2.10)}$$

The value of this spacing is obtained in Fig. S6B where we compared the total mRNA length, $\bar{\ell}_P[mR]$ (multiplied by a factor 3 to express the length in nucleotides instead of codons) and the concentration of active ribosomes $[Rb]_{\text{act}}$, yielding a value $\bar{d} \sim 193$ nt across conditions. It is worth noting that the average mRNA length really is $\bar{\ell}_{\text{mR}} \equiv \sum_i \ell_i\psi_{\text{m},i}$, and not $\bar{\ell}_{\text{P}} = \sum_i \ell_i\psi_{\text{p},i}$. The appearance of $\bar{\ell}_{\text{P}}$ in Eq. (N2.10) reflects the fact that mRNA lengths contribute in proportion to their translation efficiecies. However, we find $\bar{\ell}_{\text{P}} \approx \bar{\ell}_{\text{mR}}$ across all conditions studied, which is a consequence of the very similar translation initiation rates across the genome.

**Allowed range of initiation rates and ribosome densities**

At the quantitative level, the average initiation rate is estimated to be ~15/min in reference condition and ~12/min at slow growth (Fig. 2B). This reduction matches the growth-rate dependence of the translation elongation rate, so that the ratio of the two corresponds to a constant ribosome density (or spacing). The observed average spacing of ~200 nt is only about 5-fold of the maximum initiation rate given the minimum inter-ribosome distance of ~40 nt. This value can be justified as follows: due to the stochastic nature of ribosome translocation, high ribosome densities promoted by high initiation rates will lead to high collision frequency and reduced protein synthesis flux [48], in particular in the presence of additional mRNA sequence features slowing down elongation [12, 49–51]. In the opposite case, low initiation rates would lead to low ribosome densities and hence increase mRNA turnover, due to a combination of loss of transcriptional processivity [19] and weakening of ribosome protection against mRNA degradation [52, 53]. The resulting over-synthesis and degradation of mRNA would lead to not only a wasteful futile cycle for the mRNAs, but likely also futile protein synthesis, as prematurely terminated transcripts would leave behind partially synthesized proteins which are known to be rapidly degraded [54, 55]. We thus believe that the observed average translation initiation rates and ribosome densities strike a compromise between maximal protein synthesis rates and minimal futile gene expression.

#### Supplemental Note S3 – mRNA homeostasis

In this section we will discuss the mRNA synthesis and decay kinetics. As done previously for the protein kinetics, the goal is to build the relationship between theoretical/mechanistic quantities and experimentally accessible quantities.

Both synthesis and degradation of mRNA are taken to be first-order kinetics. The mRNA synthesis flux is proportional to the gene concentration $[g_i]$ through the rate $\alpha_{\mathrm{m},i}$ expressing the promoter activity; when multiple promoters are present, we take $\alpha_{\mathrm{m},i}$ to reflect the combined activity of all promoters. The mRNA degradation flux is instead proportional to the concentration of the mRNA, $[\mathrm{mR}_i]$, through a decay rate $\delta_i$. Since the decay rate is much smaller than the growth rate, $\delta_i \ll \lambda$, we neglect here the dilution term due to growth. Together, the mRNA kinetics take the form:

$$\frac{\mathrm{d}[\mathrm{mR}_i]}{\mathrm{d}t} = \alpha_{\mathrm{m},i}[G_i] - \delta_i[\mathrm{mR}_i]\,. \tag{N3.1}$$

Since we focused on steady state/balanced growth, we set the l.h.s. to zero, leading to the steady state constraint on the mRNA initiation flux $J_{mR,i}$:

$$J_{\mathrm{mR},i} \equiv \alpha_{\mathrm{m},i}[G_i] = \delta_i[mR_i]\,. \tag{N3.2}$$

Summing over all genes, we obtain:

$$J_{\mathrm{mR}} \equiv \sum_i \alpha_{\mathrm{m},i}[G_i] = \sum_i \delta_i[mR_{\mathrm{i}}]\,. \tag{N3.3}$$

The right hand side of this expression can be recast in terms of the average mRNA degradation rate $\bar{\delta} = \sum_i \delta_i \psi_{\mathrm{m},i}$, providing a relation between the total mRNA synthesis flux and the total mRNA concentration:

$$J_{\mathrm{mR}} = \bar{\delta}[mR]\,. \tag{N3.4}$$

By comparing Eq. (N3.2) and (N3.4), it is possible to see that the fraction of mRNA initiation flux associated to the i$^{\mathrm{th}}$ gene can be obtained from the measured mRNA fractions and mRNA degradation rates:

$$\psi_{\mathrm{Jm},i} \equiv \frac{J_{\mathrm{mR},i}}{J_{\mathrm{mR}}} = \frac{\delta_i[mR_i]}{\sum_k \delta_k[mR_{\mathrm{k}}]} = \frac{\delta_i}{\bar{\delta}}\psi_{\mathrm{m},i}. \tag{N3.5}$$

Since mRNA degradation rates were measured only in reference condition and at slow, C-limited growth, the mRNA degradation rates were linearly interpolated at the growth rates at which the mRNA fractions were measured. This allowed to obtain the mRNA synthesis fractions $\psi_{\mathrm{Jm},i}$ across the whole range of growth conditions.

**Total mRNA synthesis flux**

The absolute value of the mRNA synthesis fluxes, $J_{\mathrm{mR}}$, was obtained by directly measuring the total mRNA mass produced per unit culture per unit time ($A_{\mathrm{mR}}$) by pulse-labelling cultures with 3H-uracil and hybridizing the labelled RNA to genomic DNA over short time intervals (Fig. S8E). The total mRNA initiation flux $J_{\mathrm{mR}}$ (shown in Fig. 3FG) can then be obtained from $A_{\mathrm{mR}}$

using an approach very similar to what has been used to the total mRNA concentration (see Eq. N1.14 in Supplementary Note 1) as follows:

$$J_{\mathrm{mR}} = \frac{A_{\mathrm{mR}}}{3m_{\mathrm{nt}}\bar{\ell}_{\mathrm{JmR}}V_{\mathrm{cell}}}\,, \tag{N3.6}$$

where $3\bar{\ell}_{\mathrm{JmR}}$ is the average mRNA length (in nucleotides), weighted by the fraction of mRNA synthesis flux associated to each gene:

$$\bar{\ell}_{\mathrm{JmR}} \equiv \sum_i \ell_i \psi_{\mathrm{Jm},i}\,. \tag{N3.7}$$

These values can be directly compared to those obtained using the r.h.s. of Eq. N3.4, i.e. using the independently measured mRNA concentration and the average mRNA degradation rates. The result (crosses in Fig. 3F) show a good agreement between these two very different methods of determining the total mRNA synthesis flux.

In our calculations, we use the total mRNA synthesis flux determined from the pulse-labelling procedure to obtain $J_{\mathrm{mR}}$; synthesis fluxes of individual mRNAs $J_{\mathrm{mR},i}$ are then obtained by combining $J_{\mathrm{mR}}$ (interpolated across growth rates) with the mRNA synthesis fractions $\psi_{\mathrm{Jm},i}$ obtained as described above. These values can then be used to calculate the mRNA initiation rates as $\alpha_{\mathrm{m},i} = J_{\mathrm{mR},i}/[G_i]$ using the gene concentrations computed as described in Supplementary Note S1.

#### Supplemental Note S4 – Promoter on-rates and the quantitative Central Dogma relation

This note will explore the relation between promoter on-rates and gene expression, as well discuss in more detail the approximations assumed in the formulation of the quantitative Central Dogma relation, Eq. (12) in the Main Text. The starting point are the steady state constraints for mRNA and protein abundances. First, by taking the ratio of Eq. (N2.2) and the corresponding version summed over all genes, Eq. (N2.5), we obtain a relation linking protein and mRNA fractions:

$$\psi_{\mathrm{p},i} = \frac{\alpha_{\mathrm{p},i}}{\bar{\alpha}_{\mathrm{p}}} \psi_{\mathrm{m},i} \, . \tag{N4.1}$$

A similar relation, Eq. (N3.5), relates the mRNA fraction to the corresponding fraction of mRNA synthesis flux, as follows:

$$\psi_{\mathrm{m},i} = \frac{\bar{\delta}}{\delta_i} \cdot \frac{J_{\mathrm{mR},i}}{J_{\mathrm{mR}}} \, , \tag{N4.2}$$

where $J_{\mathrm{mR},i} = \alpha_{\mathrm{m},i}[G_i]$ and $J_{\mathrm{mR}} = \sum_i J_{\mathrm{mR},i}$. However, we want to manipulate this expression by introducing a "mechanistic" expression of the mRNA initiation rates in terms of the promoter on-rates and the concentration of RNA polymerases available to initiate transcription, $\alpha_{m,i} = k_i \cdot [\mathrm{RNAP}]_{\mathrm{av}}$, (Eq. (8) in the Main Text). (The quantification of the promoter on-rates is discussed below.) We also express the gene concentrations in terms of the Ori concentration, $[Ori]$, and the gene dose relative to Ori, $g_i$, i.e. $[G_i] = g_i[Ori]$. Using these relations, Eq. (N4.X) can be recasted as:

$$\psi_{\mathrm{m},i} = \frac{\bar{\delta}}{\delta_i} \cdot \frac{k_i g_i}{\sum_j k_j g_j} \equiv \frac{\bar{\delta}}{\delta_i} \cdot \frac{k_i g_i}{\mathcal{K}} \, , \tag{N4.3}$$

Together, Eq. (N4.1) and (N4.3) also directly connect protein fractions to the promoter on-rates, as follows:

$$\psi_{\mathrm{p},i} = \frac{\alpha_{\mathrm{p},i}}{\bar{\alpha}_{\mathrm{p}}} \psi_{\mathrm{m},i} = \frac{\alpha_{\mathrm{p},i}/\delta_i}{\bar{\alpha}_{\mathrm{p}}/\bar{\delta}} \cdot \frac{k_i g_i}{\mathcal{K}} \, . \tag{N4.4}$$

These relations represent the most general version of the quantitative Central Dogma relation, Eq. (12) in the Main Text, which we also show here below:

$$\psi_{\mathrm{p},i} \approx \psi_{\mathrm{m},i} \approx \frac{k_i g_i}{\mathcal{K}} \, . \tag{N4.5}$$

The Central Dogma relation, Eq. (N4.5), can be seen as a specialization of the more general Eq. (N4.4), obtained by assuming that translation initiation rates are similar, $\alpha_{\mathrm{p},i} \approx \bar{\alpha}_{\mathrm{p}}$, and mRNA degradation rates are similar, $\delta_i \approx \bar{\delta}$. This assumption is based on the observed similarity of mRNA characteristics for the majority of *E. coli* genes across conditions (Fig. 1, Fig. S2-S3 for the transcription initiation rates and Fig. 3D, Fig. S7E-G for the mRNA degradation rates). Our data also suggest that the relative translation initiation rates, $\alpha_{\mathrm{p},i}/\bar{\alpha}_{\mathrm{p}}$, which set the ratio between protein and mRNA fractions, and the relative mRNA degradation rates $\delta_i/\bar{\delta}$, setting the ratio between mRNA fractions and the share of transcription flux, are condition-independent for the vast majority of genes, except for a few cases attributed to post-transcriptional regulation (Fig. S4 and S7HI). The constancy of these ratios implies that changes in protein fractions, mRNA fractions and in mRNA synthesis fractions match tightly across conditions. This statement, expressed in terms of fold changes of each quantity across conditions, is captured by Main Text Eq. (13).

**Determination of promoter on-rates**
Central to the quantitative Central Dogma relation is the "molecular" relation between the mRNA transcription initiation rates, the promoter on-rates and the concentration of available RNAP. Here we summarize the determination of both genome-wide promoter on-rates and the concentration of available RNAP.

First, we considered expression from a constitutive promoter, for which $k_i$ is assumed to be constant. Under the assumption that the mRNA characteristics do not change across conditions, the Central Dogma relation relates the protein fraction to transcription as $\psi_{\mathrm{p},i} \propto g_i/\mathcal{K}$. The measured protein abundances shown in Fig. 4F, plus the knowledge of the gene dose for the constitutive promoter, allows to compute the total regulatory activity, i.e. the change in the total promoter on-rates (weighted by gene dose) $\mathcal{K}$ across condition as $\mathcal{K} \propto g_i/\psi_{\mathrm{p},i}$.

Second, the absolute value of the product $\mathcal{K}[RNAP]_{\mathrm{av}}$ can be easily obtained as the ratio between the mRNA synthesis flux $J_{\mathrm{mR}}$ and the Ori concentration $[Ori]$ as:

$$\mathcal{K}[RNAP]_{\mathrm{av}} = J_{\mathrm{mR}}/[Ori]. \tag{N4.6}$$

In order to estimate the absolute values of $\mathcal{K}$ and $[RNAP]_{\mathrm{av}}$ separately, we estimated $[RNAP]_{\mathrm{av}}$ in a single condition (reference condition, i.e. growth on glucose minimal medium, growth rate $\lambda \sim 0.9/h$) as explained in detail in Supplementary Note S5. Knowledge of the concentration of available RNAP allowed us to also compute the total promoter on-rate $\mathcal{K}$ in the same condition.

Combining the value of $\mathcal{K}$ obtained in reference condition with the growth-rate dependence estimated using constitutive promoter, the absolute value of $\mathcal{K}$ was obtained across all conditions (as reported in Fig. 4G). In turn, the knowledge of the absolute value of $\mathcal{K}$ across conditions led us to compute $[RNAP]_{\mathrm{av}}$ across conditions, as shown in Fig. 6B.

Finally, promoter on-rates for individual genes could be computed from the previously calculated fractions of mRNA synthesis flux, $\psi_{\mathrm{Jm},i} \equiv J_{\mathrm{mR},i}/J_{\mathrm{mR}}$, which are computed from the mRNA fractions and mRNA degradation rates as described in Supplementary Note S3. From the relations $J_{\mathrm{mR},i} = k_i g_i [Ori][RNAP]_{\mathrm{av}}$ and $J_{\mathrm{mR}} = \mathcal{K}[Ori][RNAP]_{\mathrm{av}}$, it is straightforward to see that $\psi_{\mathrm{Jm},i} = k_i g_i/\mathcal{K}$. Hence, the promoter on-rates can be calculated as:

$$k_i = \mathcal{K}\frac{\psi_{\mathrm{Jm},i}}{g_i}. \quad \text{(N4.7)}$$

**Quantification of determinant of gene expression**

After the determination of the promoter on-rates, we performed a global cross-gene comparison determine the main determinants of gene expression, i.e. which of the central rates and concentrations contribute the most to the vast range of protein concentrations observed in all conditions. The analysis validates quantitatively the approximation of similar mRNA characteristics performed in Eq. (N4.4) to yield the Central Dogma relation, Eq. (N4.5). The starting point is the expression of the protein concentrations from the fundamental rates described by Eq. (S5) in Figure S1, which is obtained by combining the steady state equations for mRNA and protein concentrations. We report here the expression for convenience of the reader:

$$[P_i] = \frac{\alpha_{\mathrm{p},i}}{\delta_i}\frac{\alpha_{\mathrm{m},i}[G_i]}{\lambda}. \quad \text{(N4.8)}$$

Introducing once again the "mechanistic" relation for the transcription initiation rates, $\alpha_{m,i} = k_i \cdot [\mathrm{RNAP}]_{\mathrm{av}}$, we obtain:

$$[P_i] = \alpha_{\mathrm{p},i} \cdot \frac{1}{\delta_i} \cdot k_i \cdot [G_i] \cdot \frac{[RNAP]_{\mathrm{av}}}{\lambda}. \quad \text{(N4.9)}$$

This expression expresses the protein concentration as the product of four gene-specific factors: the translation initiation rate $\alpha_{\mathrm{p},i}$, the inverse of the mRNA degradation rate $\delta_i$, the promoter on-rate $k_i$, and the gene concentration $[G_i]$. This expression can be used to study the extent by which each of these four factors determine the observed protein abundances across different genes, as illustrated in Fig. 5C. Note that the term $[RNAP]_{\mathrm{av}}/\lambda$ acts as a global multiplicative factor, and therefore it is inconsequential when comparing the protein concentrations of different genes in the same growth condition. In this analysis, it is important that the four terms are determined from the available data in a manner that is consistent with their product being proportional to the protein concentrations. Since $\alpha_{\mathrm{p},i}$ are computed from the measured ratios of protein and mRNA fractions, $\psi_{\mathrm{p},i}/\psi_{\mathrm{m},i}$, and the promoter on-rates $k_i$ are proportional to $\delta_i\psi_{\mathrm{m},i}/[G_i]$, it can be verified that the product of the four terms simplifies exactly to the protein concentrations (or protein fractions), apart from a global multiplicative factor.

Equation (N4.9) also provides the basis for the analysis shown in Figure 5D, where we compare fold changes of protein concentrations between reference condition and slow growth to the fold changes of each of the four gene-specific terms. Again, the product of the fold changes in each of the four quantities is guaranteed to yield the fold change in the protein concentrations, apart from a global factor. The result of the analysis highlights how changes in the concentration of one protein relative to the change of another one are primarily due to differential promoter-level regulation.

**Promoter activity**

The protein synthesis flux $\lambda[P_i]$, given as the product of the growth rate $\lambda$ and the protein concentration $[P_i]$, is a convenient measure of gene expression that has been widely used in the literature as a proxy of gene regulation under the name of "promoter activity" [56, 57]. Slightly recasting the expression relating the protein concentrations to the fundamental gene expression rates, Eq. (N4.1), yields:

$$\lambda[P_i] = \frac{\alpha_{\mathrm{p},i}[G_i]}{\delta_i} \cdot \alpha_{\mathrm{m},i}. \tag{N4.10}$$

In this expression, both the translation initiation rates, the mRNA degradation rates and the gene concentrations have little (<2-fold) gene-to-gene variation. Furthermore, the mRNA degradation rates are mostly condition-independent, while the average translation initiation rates and gene concentrations have mild and opposite dependences on the growth rates, leading to a partial compensation. Hence, in absence of post-transcriptional regulation, the protein synthesis flux is a good estimate of the transcription initiation rate $\alpha_{\mathrm{m},i}$, i.e.

$$\lambda[P_i] \propto \alpha_{\mathrm{m},i} \tag{N4.11}$$

across genes and conditions. However, an important consequence of Rsd-mediated regulation is the growth rate dependence of the available RNAP, which imposes a global growth rate dependence on the transcription initiation rate $\alpha_{m,i} = k_i[RNAP]_{av}$ of each promoter $i$. This global effect is on top of the specific regulatory effects acting on the promoter, represented by the promoter on-rate $k_i$. Because of the strong growth-rate dependence of $[RNAP]_{\mathrm{av}}$, the quantity $\lambda[P_i]$ does not faithfully represent changes in the regulatory activity of a gene.

Remarkably, the growth-rate dependence of $[RNAP]_{\mathrm{av}}$ appears to be not far off a direct proportionality with the growth rate itself $\lambda$ (Fig. 6B). A perfect match between the availability of RNA polymerases and the growth rate would simplify Eq. (N4.11), leading to a direct proportionality between protein concentrations and promoter on-rates, $[P_i] \propto k_i$. This cannot be quite true given the constancy of the total protein concentration vs. the global change in promoter on-rates observed across conditions. To obtain a better estimate of the regulatory activity of a promoter, we instead return to the general relation between protein concentrations and the fundamental rates of gene expression in Eq. (N4.9). Using the balance between protein synthesis flux and the number of elongating ribosomes, $\lambda\bar{\ell}_P[P] = \varepsilon[\mathrm{Rb}]_{\mathrm{act}}$ (Eq. (5) or Eq. (N2.8)), it is possible to express the growth rate in terms of the abundance of translating ribosomes. Using this relation in Eq. (N4.9) yields:

$$[P_i] = \frac{\alpha_{\mathrm{p},i}}{\delta_i \varepsilon} \cdot \frac{[\mathrm{RNAP}]_{\mathrm{av}}}{[\mathrm{Rb}]_{\mathrm{act}}} \cdot \left(\bar{\ell}_{\mathrm{P}}[P]\right) \cdot k_i[G_i]\,. \tag{N4.12}$$

Let us analyze the terms individually. The first term, $\alpha_{\mathrm{p},i}/\delta_i\varepsilon$, is the inverse of the product of the mRNA decay rate and the mRNA inter-ribosome spacing $d_i = \varepsilon/\alpha_{\mathrm{p},i}$. Both of these quantities are mostly condition independent (Fig. S6 and S7), and do not vary by more than 2-fold across genes; furthermore, both the average inter-ribosome spacing and the mRNA degradation rates are remarkably condition-independent (Fig. S6F, Fig. 3D). The second term is the ratio $[\mathrm{RNAP}]_{\mathrm{av}}/[\mathrm{Rb}]_{\mathrm{act}}$ of the available RNA polymerases and the active ribosomes. As seen in Figure 6B, the two are proportional to each other, with about 11 elongating ribosomes per available RNAP. Similarly, the third term $\bar{\ell}_{\mathrm{P}}[P]$ is also mostly growth-independent, given the constancy of total protein concentration and the very weak dependence of the average protein size on the growth rate (Fig. S2). Given the constancy of all these quantities, the protein abundances are approximately proportional across conditions to the product of the gene concentration $[G_i]$ and the promoter affinity $k_i$:

$$[P_i] \propto k_i[G_i] \tag{N4.13}$$

Indeed, the quantity $\mathcal{K}[Ori] = \sum_i k_i[G_i]$ has a much milder dependence on the growth rate compared to $[\mathrm{RNAP}]_{\mathrm{av}}$ (compare Fig. 6A and Fig. 6B). Therefore, in our growth conditions, promoter-level regulation is better represented by the ratio of protein and gene concentrations, $[P_i]/[G_i]$, compared to the protein synthesis flux.

### Supplemental Note S5 - Determination of the absolute concentration of available RNAP

In this Note, we discuss our estimate of the absolute concentration of available RNAP, necessary to assign an absolute scale to the promoter on-rates. The total mRNA flux can be expressed in terms of the available RNA polymerases, DNA concentration and total promoter on-rates as described by Main Text Eq. (10), which we also report here below in terms of the total mRNA flux $J_{\mathrm{mR}}$:

$$J_{\mathrm{mR}} = \mathcal{K}[Ori][RNAP]_{\mathrm{av}} \,. \tag{N5.1}$$

Of these quantities, $J_{\mathrm{mR}}$ and [Ori] are known across growth conditions (Fig. 3G, 4A). The relative value of $\mathcal{K}$ has been determined in Fig. 4G by measuring the expression of LacZ driven by a constitutive promoter. Using these quantities as inputs in Eq. (N5.1), it is possible to calculate the change of $[RNAP]_{av}$ across conditions. However, the absolute magnitude of $[RNAP]_{\mathrm{av}}$ cannot be known without the knowledge of that of $\mathcal{K}$ in at least one condition, and vice versa. The aim of this Note is to estimate $[RNAP]_{av}$ for *E. coli* cells in reference condition, i.e. growth on glucose minimal medium, corresponding to a growth rate $\lambda \sim 0.9/\mathrm{h}$. This allows both to estimate $\mathcal{K}$ and to assign an absolute value to the promoter on-rates $k_i$ as discussed in Supplementary Note S4.

##### The concentration of available RNAP is set by $\sigma^{70}$

Mechanistically, $[RNAP]_{av}$ is set by the cellular concentration of holoenzymes ready for transcription initiation. It is well known that most transcription is driven by the housekeeping sigma factor $\sigma^{70}$ (encoded by *rpoD*), which a limited role played by alternative sigma factors during exponential growth [58]. The crucial role of $\sigma^{70}$ in globally regulating transcription is highlighted by the two following results:

> (*i*) The concentrations of the constituents of the core enzyme (RpoA, RpoB and RpoC) are constant across growth rates, and in excess of that of $\sigma^{70}$ (Fig. 6C). This suggest that the core RNAP is not limiting transcription.
>
> (*ii*) The deletion of the *rsd* gene, encoding for the anti-$\sigma^{70}$ factor Rsd, leads to a dramatic increase in the synthesis of total RNA compared to wild-type at slow growth (Fig. 6E), when Rsd is expressed at levels comparable to those of $\sigma^{70}$ (Fig. 6C). This suggests that transcription is limited by the available $\sigma^{70}$ in the cell.

We can therefore take as a first estimate of the concentration of available RNAP the concentration of $\sigma^{70}$ in reference condition:

$$[RNAP]_{\mathrm{av}} \sim [\sigma^{70}] \sim 1473/\mu m^3 \,. \tag{N5.2}$$

##### Impact of TEC-$\sigma^{70}$ binding

The simple picture above is complicated by the fact that $\sigma^{70}$ can bind to transcription elongation complexes (TECs) for some time after transcription initiation, or ever detach and re-bind stochastically to TECs [59, 60]; furthermore, transcription elongation can pause on promoter-like regions in a $\sigma^{70}$-dependent manner [61]. This phenomenon can potentially limit the pool of $\sigma^{70}$ available for transcription. Kinetic experiments in vitro have suggested timescales for the release of $\sigma^{70}$ after initiation ranging from <5 seconds [62] to more than 30 minutes [63, 64]. This large spread in timescales can be reconciled with a fraction of $\sigma^{70}$ detaching immediately after initiation, and the remainder of TEC-$\sigma^{70}$ complexes being long-lived. Both in vivo [65, 66] and in vitro data [67] suggest that about 70% of $\sigma^{70}$ are quickly released from TECs after initiation.

We estimated the concentration of elongating complexes using the measured mRNA synthesis fluxes and the observed abundance of stable RNA. The number of elongated nucleotides per cell volume can be derived from the measured mRNA synthesis flux as $A_{\mathrm{mR}}/m_{\mathrm{nt}}/V_{\mathrm{cell}}$ (see also Supplementary Note S3). This quantity can be expressed as the product of the mRNA transcription elongation rate $\varepsilon_{\mathrm{mR}}$ and the concentration of transcription elongation complexes transcribing mRNAs, $[TEC]_{\mathrm{mR}}$:

$$\frac{A_{\mathrm{mR}}}{m_{\mathrm{nt}}V_{\mathrm{cell}}} = \varepsilon_{\mathrm{mR}}[TEC]_{\mathrm{mR}}\,. \qquad \text{(N5.3)}$$

The mRNA transcription elongation rate is similar to the translation elongation rate $\varepsilon$ across growth conditions [19]. Using $\varepsilon_{\mathrm{mR}} = 47\ \mathrm{nt/s}$, and an mRNA synthesis flux of $1.85 \times 10^4\ \mathrm{nt/}$ $\mathrm{\mu m^3/s}$, we obtain $[TEC]_{\mathrm{mR}} = 394/\mu m^3$.

The concentration of TECs associated to the synthesis of stable RNA is obtained in a similar way. Since the vast majority of RNA in the cell is stable, we estimate the synthesis flux as the product of the growth rate $\lambda$ and the abundance of stable RNA ($W_{\mathrm{R}}/V_{\mathrm{cell}}/m_{\mathrm{nt}}$, in units of nucleotides per cell volume). As in the previous case, this quantity is equal to the product of the transcriptional elongation rate $\varepsilon_{\mathrm{sR}}$ and the concentration of elongation complexes, $[TEC]_{\mathrm{sR}}$:

$$\frac{\lambda W_{\mathrm{R}}}{m_{\mathrm{nt}}V_{\mathrm{cell}}} = \varepsilon_{\mathrm{sR}}[TEC]_{\mathrm{sR}}\,. \qquad \text{(N5.4)}$$

The elongation rate of ribosomal RNA is high compared to mRNA operons, with $\varepsilon_{\mathrm{rR}} = 85\ nt/s$ [68]. At a growth rate of $0.9/\mathrm{h}$, we obtain a synthesis flux of $1.69 \times 10^4\ \mathrm{nt/\mu m^3/s}$, leading to $[TEC]_{\mathrm{sR}} = 229/\mu m^3$. Together, the elongation of mRNA and stable RNA lead to $[TEC] = [TEC]_{\mathrm{mR}} + [TEC]_{\mathrm{sR}} = 623/\mu m^3$. Assuming that only 30% of the $\sigma^{70}$ are retained on a long timescale, we obtain a revised estimate:

$$[RNAP]_{av} \sim [\sigma^{70}] - 0.3 \times [TEC] \sim 1286/\mu m^3\,. \qquad \text{(N5.5)}$$

**Anti-sigma factors and final estimate**

We finally consider the effect of anti-sigma factors, specifically Rsd. The number of $\sigma^{70}$ inactivated by Rsd depends on the $\sigma^{70}$-Rsd dissociation constant and the total concentrations of $\sigma^{70}$ and Rsd; in principle, the number of inactivated $\sigma^{70}$ could range between zero and the concentration of Rsd. However, in reference condition, the concentration of Rsd is much lower than that of $\sigma^{70}$. In the extreme case in which each Rsd inactivates a $\sigma^{70}$ unit, the concentration of available RNAP in reference condition would be reduced by $[Rsd] \approx 328/\mu m^3$, leading to the estimate:

$$[RNAP]_{\mathrm{av}} \sim [\sigma^{70}] - 0.3 \times [TEC] - [Rsd] \sim 958/\mu m^3\,. \qquad \text{(N5.6)}$$

We take Eq. (N5.6) as our estimate for the concentration of $[RNAP]_{\mathrm{av}}$ at a growth rate of $\lambda = 0.9/\mathrm{h}$, as seen in Fig. 6B. Other effects could potentially affect this estimate: the actual value might be slightly higher due to incomplete binding of Rsd to $\sigma^{70}$, or lower due to a fraction of

$\sigma^{70}$ not being bound to core RNAP enzymes. These effects are hard to estimate, but their magnitude should be small. Indeed, in this condition the concentration of Rsd is low, and the dissociation constant between core RNAP and $\sigma^{70}$ is low enough (sub-nanomolar range, [69] so that the concentration of free $\sigma^{70}$ can be neglected compared to that of RNAP-$\sigma^{70}$ holoenzyme. From this value, plus $J_{\mathrm{mR}} = 1482\ \mathrm{\mu m^3/min}$ and $[Ori] = 1.33/\mathrm{\mu m^3}$, one derives the value $\mathcal{K} = 1.16/\mathrm{min}$ for the total promoter on-rate and $\mathcal{K}[Ori] = 1.54/\mathrm{min}$ in reference condition; these absolute values across growth conditions of these two quantities are shown in Fig. 4G and Fig. 6A, respectively.

### SUPPLEMENTAL INFORMATION REFERENCES


1. Paulsson, J. “Models of Stochastic Gene Expression” *Physics of Life Reviews* **2**: 157–175 (2005).
2. Li, G. W. “How Do Bacteria Tune Translation Efficiency?” *Current Opinion in Microbiology* **24**: 66–71 (2015).
3. Goldberg, A. L. and St John, A. C. “Intracellular Protein Degradation in Mammalian and Bacterial Cells: Part 2.” *Annual review of biochemistry* **45**: 747–803 (1976).
4. Nath, K. and Koch, A. L. “Protein Degradation in Escherichia Coli. II. Strain Differences in the Degradation of Protein and Nucleic Acid Resulting from Starvation.” *Journal of Biological Chemistry* **246**: 6956–6967 (1971).
5. Scott, M., Gunderson, C. W., Mateescu, E. M., Zhang, Z., and Hwa, T. “Interdependence of Cell Growth” *Science* **330**: 1099–1102 (2010).
6. Basan, M., Zhu, M., Dai, X., Warren, M., Sévin, D., Wang, Y.-P., and Hwa, T. “Inflating Bacterial Cells by Increased Protein Synthesis” *Mol Syst Biol* **11**: 836 (2015).
7. Mori, M., Zhang, Z., Esfahani, A. B., Lalanne, J., Collins, B. C., Schmidt, A., Schubert, O. T., Lee, D., Li, G. W., Hwa, T., and Ludwig, C. “From Coarse to Fine : The Absolute Escherichia Coli Proteome under Diverse Growth Conditions” *Mol Syst Biol* **17**:e9536 (2021)
8. Woldringh, C. L., Binnerts, J. S., and Mans, A. “Variation in Escherichia Coli Buoyant Density Measured in Percoll Gradients” *Journal of Bacteriology* **148**: 58–63 (1981).
9. Schwanhüusser, B., Busse, D., Li, N., Dittmar, G., Schuchhardt, J., Wolf, J., Chen, W., and Selbach, M. “Global Quantification of Mammalian Gene Expression Control” *Nature* **473**: 337–342 (2011).
10. Taniguchi, Y., Choi, P. J., Li, G.-W., Chen, H., Babu, M., Hearn, J., Emili, A., and Xie, X. S. “Quantifying E. Coli Proteome and Transcriptome with Single-Molecule Sensitivity in Single Cells”
11. Li, G.-W., Burkhardt, D., Gross, C., and Weissman, J. S. “Quantifying Absolute Protein Synthesis Rates Reveals Principles Underlying Allocation of Cellular Resources.” *Cell* **157**: 624–35 (2014).
12. Campo, C. Del, Bartholomäus, A., Fedyunin, I., and Ignatova, Z. “Secondary Structure across the Bacterial Transcriptome Reveals Versatile Roles in MRNA Regulation and Function” *PLOS Genetics | DOI* (2015).
13. Choe, D., Lee, J. H., Yoo, M., Hwang, S., Sung, B. H., Cho, S., Palsson, B., Kim, S. C., and Cho, B.-K. “Adaptive Laboratory Evolution of a Genome-Reduced Escherichia Coli”
14. Friesen, J. D. “Control of Messenger RNA Synthesis and Decay in Escherichia Coli” *Journal of Molecular Biology* **20**: 559–573 (1966).
15. Kennell, D. “Titration of the Gene Sites on DNA by DNA-RNA Hybridization. II. The Escherichia Coli Chromosome” *Journal of Molecular Biology* **34**: 85–103 (1968).
16. Norris, T. E. and Koch, A. L. “Effect of Growth Rate on the Relative Rates of Synthesis of Messenger, Ribosomal and Transfer RNA in Escherichia Coli” *Journal of Molecular Biology* **64**: 633–649 (1972).
17. Dai, X., Zhu, M., Warren, M., Balakrishnan, R., Patsalo, V., Okano, H., Williamson, J. R., Fredrick, K., Wang, Y.-P., Hwa, T., Frederick, K., Wang, Y.-P., and Hwa, T. “Reduction of Translating Ribosomes Enables Escherichia Coli to Mantain Elongation Rates during Slow Growth” *Nature Microbiology* **2**: 16231 (2016).

18. Mohammad, F., Green, R., and Buskirk, A. R. "A Systematically-Revised Ribosome Profiling Method for Bacteria Reveals Pauses at Single-Codon Resolution"
19. Zhu, M., Mori, M., Hwa, T., and Dai, X. "Disruption of Transcription-Translation Coordination in Escherichia Coli Leads to Premature Transcriptional Termination" *Nature Microbiology* **4**: (2019).
20. Sowa, S. W., Gelderman, G., Leistra, A. N., Buvanendiran, A., Lipp, S., Pitaktong, A., Vakulskas, C. A., Romeo, T., Baldea, M., and Contreras, L. M. "Integrative FourD Omics Approach Profiles the Target Network of the Carbon Storage Regulatory System" *Nucleic acids research* **45**: 1673–1686 (2017).
21. Chen, S., Zhang, A., Blyn, L. B., and Storz, G. "MicC, a Second Small-RNA Regulator of Omp Protein Expression in Escherichia Coli" *Journal of Bacteriology* **186**: 6689–6697 (2004).
22. Wang, J., Rennie, W., Liu, C., Carmack, C. S., Prévost, K., Caron, M. P., Massé, E., Ding, Y., and Wade, J. T. "Identification of Bacterial SRNA Regulatory Targets Using Ribosome Profiling" *Nucleic Acids Research* **43**: 10308–10320 (2015).
23. Lay, N. De and Gottesman, S. "A Complex Network of Small Non-Coding RNAs Regulate Motility in Escherichia Coli" *Molecular Microbiology* **86**: 524–538 (2012).
24. Thomason, M. K., Fontaine, F., Lay, N. De, and Storz, G. "A Small RNA That Regulates Motility and Biofilm Formation in Response to Changes in Nutrient Availability in Escherichia Coli" *Molecular Microbiology* **84**: 17–35 (2012).
25. Baracchini, E. and Bremer, H. "Control of RRNA Synthesis in Escherichia Coli at Increased Rrn Gene Dosage: Role of Guanosine Tetraphosphate and Ribosome Feedback" *Journal of Biological Chemistry* **266**: 11753–11760 (1991).
26. Ryals, J., Little, R., and Bremer, H. "Control of RRNA and TRNA Syntheses in Escherichia Coli by Guanosine Tetraphosphate" *Journal of Bacteriology* **151**: 1261–1268 (1982).
27. Zheng, H., Bai, Y., Jiang, M., Tokuyasu, T. A., Huang, X., Zhong, F., Wu, Y., Fu, X., Kleckner, N., Hwa, T., and Liu, C. "General Quantitative Relations Linking Cell Growth and the Cell Cycle in Escherichia Coli" *Nature Microbiology* **5**: 995–1001 (2020).
28. Cooper, Stephen and Helmstetter, C. "Chromosome Replication and the Division Cycle of Escherichia Coli B/R" *J. Mol. Biol.* **31**: 519–540 (1968).
29. Si, F., Li, D., Cox, S. E., Sauls, J. T., Azizi, O., Sou, C., Schwartz, A. B., Erickstad, M. J., Jun, Y., Li, X., and Jun, S. "Invariance of Initiation Mass and Predictability of Cell Size in Escherichia Coli" *Current Biology* **27**: 1278–1287 (2017).
30. Karp, P. D., Ong, W. K., Paley, S., Billington, R., Caspi, R., Fulcher, C., Kothari, A., Krummenacker, M., Latendresse, M., Midford, P. E., Subhraveti, P., Gama-Castro, S., Muñiz-Rascado, L., Bonavides-Martinez, C., Santos-Zavaleta, A., Mackie, A., Collado-Vides, J., Keseler, I. M., and Paulsen, I. "The EcoCyc Database" *EcoSal Plus* **8**: 1–19 (2018).
31. Feucht, B. U. and Saier, M. H. "Fine Control of Adenylate Cyclase by the Phosphoenolpyruvate:Sugar Phosphotransferase Systems in Escherichia Coli and Salmonella Typhimurium" *Journal of Bacteriology* **141**: 603–610 (1980).
32. Busby, S. and Ebright, R. H. "Transcription Activation by Catabolite Activator Protein (CAP)" *Journal of Molecular Biology* **293**: 199–213 (1999).
33. You, C., Okano, H., Hui, S., Zhang, Z., Kim, M., Gunderson, C. W., Wang, Y.-P., Lenz, P., Yan, D., and Hwa, T. "Coordination of Bacterial Proteome with Metabolism by Cyclic AMP Signalling" (2013).
34. Kolb, A., Busby, S., Buc, H., Gorges, S., and Adhya, S. "Transcriptional Regulation by CAMP and Its Receptor Protein" *Annual Review of Biochemistry* **62**: 749–795 (1993).

35. Epstein, W., Rothman Denes, L. B., and Hesse, J. "Adenosine 3':5' Cyclic Monophosphate as Mediator of Catabolite Repression in Escherichia Coli" *Proceedings of the National Academy of Sciences of the United States of America* **72**: 2300–2304 (1975).
36. Park, S. -J, Tseng, C. -P, and Gunsalus, R. P. "Regulation of Succinate Dehydrogenase SdhCDAB Operon Expression in Escherichia Coli in Response to Carbon Supply and Anaerobiosis: Role of ArcA and Fnr" *Molecular Microbiology* **15**: 473–482 (1995).
37. Lynch, A. S. and Lin, E. C. C. "Transcriptional Control Mediated by the ArcA Two-Component Response Regulator Protein of Escherichia Coli: Characterization of DNA Binding at Target Promoters" *Journal of Bacteriology* **178**: 6238–6249 (1996).
38. Soupene, E., Heeswijk, W. C. Van, Plumbridge, J., Stewart, V., Bertenthal, D., Lee, H., Prasad, G., Paliy, O., Charernnoppakul, P., and Kustu, S. "Physiological Studies of Escherichia Coli Strain MG1655: Growth Defects and Apparent Cross-Regulation of Gene Expression" *JOURNAL OF BACTERIOLOGY* **185**: 5611–5626 (2003).
39. Neidhardt, F. C., Bloch, P. L., and Smith, D. F. "Culture Medium for Enterobacteria" *Journal of Bacteriology* **119**: 736–747 (1974).
40. Kochanowski, K., Volkmer, B., Gerosa, L., Rijsewijk, B. R. H. Van, Schmidt, A., and Heinemann, M. "Functioning of a Metabolic Flux Sensor in Escherichia Coli" *Proceedings of the National Academy of Sciences of the United States of America* **110**: 1130–1135 (2013).
41. Langmead, B. and Salzberg, S. L. "Fast Gapped-Read Alignment with Bowtie 2." *Nature methods* **9**: 357–9 (2012).
42. Anders, S., Pyl, P. T., and Huber, W. "Genome Analysis HTSeq-a Python Framework to Work with High-Throughput Sequencing Data" **31**: 166–169 (2015).
43. Pigott, G. H. and Midgleyt, J. E. M. "Characterization of Rapidly Labelled Ribonucleic Acid in Escherichia Coli by Deoxyribonucleic Acid-Ribonucleic Acid Hybridization" *Biochem. J* **110**: (1968).
44. Gillespie, D. and Spiegelman, S. "A Quantitative Assay for DNA-RNA Hybrids with DNA Immobilized on a Membrane" *Journal of Molecular Biology* **12**: 829–842 (1965).
45. Chen, H., Shiroguchi, K., Ge, H., and Sunney Xie, X. "Genome-Wide Study of MRNA Degradation and Transcript Elongation in Escherichia Coli" *Mol Syst Biol* **11**: 781 (2015).
46. Schmidt, A., Kochanowski, K., Vedelaar, S., Ahrné, E., Volkmer, B., Callipo, L., Knoops, K., Bauer, M., Aebersold, R., and Heinemann, M. "The Quantitative and Condition-Dependent Escherichia Coli Proteome" *Nature Biotechnology* **34**: 104–110 (2016).
47. Gottesman, S. "Proteases and Their Targets in Escherichia Coli" *Annual Review of Genetics* **30**: 465–506 (1996).
48. Zarai, Y., Margaliot, M., and Tuller, T. "On the Ribosomal Density That Maximizes Protein Translation Rate" (2016).
49. Buskirk, A. R. and Green, R. "Ribosome Pausing, Arrest and Rescue in Bacteria and Eukaryotes"
50. Ude, S., Lassak, J., Starosta, A. L., Kraxenberger, T., Wilson, D. N., and Jung, K. "Translation Elongation Factor EF-P Alleviates Ribosome Stalling at Polyproline Stretches"
51. Thanaraj, T. A. and Argos, P. "Ribosome-Mediated Translational Pause and Protein Domain Organization" *Protein Science* **5**: 1594–1612 (1996).
52. Deana, A. and Belasco, J. G. "Lost in Translation: The Influence of Ribosomes on Bacterial MRNA Decay" *Genes and Development* **19**: 2526–2533 (2005).
53. Hui, M. P., Foley, P. L., and Belasco, J. G. "Messenger RNA Degradation in Bacterial Cells" **7**: 19 (2014).

70. Waldminghaus, T., Gaubig, L. C., Klinkert, B., and Narberhaus, F. “RNA Biology The Escherichia Coli IbpA Thermometer Is Comprised of Stable and Unstable Structural Elements” *RNA Biology* **6**: 455–463
71. Massé, E., Vanderpool, C. K., and Gottesman, S. “Effect of RyhB Small RNA on Global Iron Use in Escherichia Coli Downloaded from MATERIALS AND METHODS” *JOURNAL OF BACTERIOLOGY* **187**: 6962–6971 (2005).
72. Nafissi, M., Chau, J., Xu, J., and Johnson, R. C. “Robust Translation of the Nucleoid Protein Fis Requires a Remote Upstream AU Element and Is Enhanced by RNA Secondary Structure” (2012).
73. Balasubramanian, D., Ragunathan, P. T., Fei, J., and Vanderpool, C. K. “A Prophage-Encoded Small RNA Controls Metabolism and Cell Division in Escherichia Coli” (2016).
74. Modi, S. R., Camacho, D. M., Kohanski, M. A., Walker, G. C., and Collins, J. J. “Functional Characterization of Bacterial SRNAs Using a Network Biology Approach”
75. Waldminghaus, T., Gaubig, L. C., Klinkert, B., and Narberhaus, F. “RNA Biology The Escherichia Coli IbpA Thermometer Is Comprised of Stable and Unstable Structural Elements” *RNA Biology* **6**: 455–463
76. Yaagoubi, A. El, Kohiyama, M., and Richarme, G. “Defect in Export and Synthesis of the Periplasmic Galactose Receptor MglB in DnaK Mutants of Escherichia Coli, and Decreased Stability of the Mg/B MRNA” *Microbiology* **142**: (1996).
77. Zhang, A., Wassarman, K. M., Rosenow, C., Tjaden, B. C., Storz, G., and Gottesman, S. “Global Analysis of Small RNA and MRNA Targets of Hfq” *Molecular Microbiology* **50**: 1111–1124 (2003).
78. Langklotz, S. and Narberhaus, F. “The Escherichia Coli Replication Inhibitor CspD Is Subject to Growth-Regulated Degradation by the Lon Proteasem Mi_7646 1313..1325” (2011).
79. Hatoum, A. and Roberts, J. “Prevalence of RNA Polymerase Stalling at Escherichia Coli Promoters after Open Complex Formation” (2008).
80. Wright, P. R., Richter, A. S., Papenfort, K., Mann, M., Vogel, J., Hess, W. R., Backofen, R., and Georg, J. “Comparative Genomics Boosts Target Prediction for Bacterial Small RNAs”
81. Potts, A. H., Vakulskas, C. A., Pannuri, A., Yakhnin, H., Babitzke, P., and Romeo, T. “Global Role of the Bacterial Post-Transcriptional Regulator CsrA Revealed by Integrated Transcriptomics”
82. Nikolic, N., Moll, I., and Didara, Z. “MazF Activation Promotes Translational Heterogeneity of the GrcA MRNA in Escherichia Coli Populations” (2017).
83. Melamed, S., Peer, A., Faigenbaum-Romm, R., Gatt, Y. E., Reiss, N., Bar, A., Altuvia, Y., Argaman, L., and Margalit, H. “Global Mapping of Small RNA-Target Interactions in Bacteria” *Molecular Cell* **63**: 884–897 (2016).
84. Kolmsee, T. and Hengge, R. “RNA Biology Rare Codons Play a Positive Role in the Expression of the Stationary Phase Sigma Factor RpoS (ΣS) in Escherichia Coli” *RNA Biology* **8**: 913–921
